# Integration of Infinium and Axiom SNP array data in the outcrossing species *Malus* × *domestica* and causes for seemingly incompatible calls

**DOI:** 10.1101/2020.09.01.276758

**Authors:** Nicholas P. Howard, Michela Troggio, Charles-Eric Durel, Hélène Muranty, Caroline Denancé, Luca Bianco, John Tillman, Eric van de Weg

## Abstract

**Background:** Single nucleotide polymorphism (SNP) array technology has been increasingly used to generate large quantities of SNP data for use in genetic studies. As new arrays are developed to take advantage of new technology and of improved probe design using new genome sequence and panel data, a need to integrate data from different arrays and array platforms has arisen. This study was undertaken in view of our need for an integrated high-quality dataset of Illumina Infinium^®^ 20K and Affymetrix Axiom^®^ 480K SNP array data in apple (*Malus* × *domestica*). In this study, we qualify and quantify the compatibility of SNP calling, defined as SNP calls that are both accurate and concordant, across both arrays by two approaches. First, the concordance of SNP calls was evaluated using a set of 417 duplicate individuals genotyped on both arrays starting from a set of 10,295 robust SNPs on the Infinium array. Next, the accuracy of the SNP calls was evaluated on additional germplasm (n=3,141) from both arrays using Mendelian inconsistent and consistent errors across thousands of pedigree links. While performing this work, we took the opportunity to evaluate reasons for probe failure and observed discordant SNP calls.

**Results:** Concordance among the duplicate individuals was on average of 97.1% across 10,295 SNPs. Of these SNPs, 35% had discordant call(s) that were further curated, leading to a final set of 8,412 (81.7%) SNPs that were deemed compatible. Compatibility was highly influenced by the presence of alternate probe binding locations and secondary polymorphisms. The impact of the latter was highly influenced by their number and proximity to the 3’ end of the probe.

**Conclusions:** The Infinium and Axiom SNP array data were mostly compatible. However, data integration required intense data filtering and curation. This work resulted in a workflow and information that may be of use in other data integration efforts. Such an in-depth analysis of array concordance and accuracy as ours has not been previously described in literature and will be useful in future work on SNP array data integration and interpretation, and in probe/platform development.

## Background

Single nucleotide polymorphism (SNP) array technology has been increasingly used to generate large quantities of SNP data for use in genetic studies. Over time, next generation arrays are developed that use new sequence data and/or new genome drafts to either refine or expand upon the set of SNPs used in previous arrays. Additionally, different SNP array technologies have been developed, resulting in different array platforms, creating a need for data harmonization and integration [1].

This need has been faced in apple (*Malus* × *domestica*), where a large amount of SNP array data has been generated using the Infinium^®^ IRSC 8K [2] and 20K apple SNP arrays [3] on thousands of accessions through over thirty published, as well as ongoing, studies on pedigree reconstruction, genetic linkage map construction, identification of polyploids and aneuploids, quantitative trait loci identification, genome-wide association, and genomic selection; this data has also been used in downstream research like de novo genome assemblies and methodology development for the calling of SNP [2,4-33]. These previous and on-going studies have previously relied on a single SNP array platform, however a recent study provided whole genome SNP data on over 1400 mostly old, unique apple cultivars [32] using the Affymetrix Axiom^®^ Apple480K SNP array [34]. Hence, ongoing and following studies on genetic relationships among apple cultivars could greatly benefit from the integration of data across these platforms.

When newer arrays are simply updated arrays with additional SNPs that utilize the same platform, an evaluation of concordance of data among common accessions is straightforward and concordance is often high, such as with the BovineLD BeadChip [35] and the barley 50K iSelect SNP array [36]. Concordance between the Infinium 8K and 20K apple SNP arrays has not been reported, but integration of SNP data across these arrays was seamless in Vanderzande et al. [29]. However, when SNP calls are compared across different platforms that use different technology, such as between the Infinium 20K and the Axiom 480K apple SNP arrays, concordance rates may be more variable because of differences in the chemistry, probe lengths, probe densities used across these platforms, and/or differences in the genotyped germplasm. Concordance rates between the Illumina Infinium 20K and Affymetrix Axiom Apple 480K SNP array data were reported as 96% to 98%, based on 53 common individuals [34]. This high rate is promising and is in line with those found in other organisms: an average concordance of 96% to 98.8% was reported in human [37], sheep [38], and swine [39]. However, levels of concordance were not documented at the individual SNP level, as would be needed for accurate data integration. Also, this nor any of the studies in other organisms reported on the technical or biological reasons for the observed SNP call discordances.

So far, compatibility between array platforms has been determined by evaluating the concordance of SNP calls of genetically duplicate samples genotyped on both platforms, which is usually limited by few individuals. Such evaluations would be made more useful by considering SNP call accuracy via assessments of Mendelian inconsistent and Mendelian consistent errors [40] across direct parent-offspring relationships. This approach could increase the amount of informative comparisons and could also expand the data chain to multiple successive generations. Moreover, the use of inheritance patterns surpasses analyses on duplicate genotypes in determining the precise genotype of an individual, allowing, for instance, the revealing of null alleles. The power of compatibility studies may increase even further by an integrated Mendelian error analyses on a mixed data set, rather than within array analyses. The identification and troubleshooting of Mendelian inconsistent and consistent errors have been previously described using Infinium SNP array data in apple, cherry, and peach [29]. Extensive pedigree information exists for apple from breeding records (e. g. from websites such as https://hort.purdue.edu/newcrop/pri/), pomological textbooks (e. g. [41]), historic pedigree reconstruction studies [6,20,27,29,31,32,33,42-45], and may also be revealed by the available SNP data.

Mendelian inconsistent and consistent errors in SNP array data can result from the presence of secondary polymorphism(s) on probe sites and/or the presence of duplicated or paralogous sequences. Secondary polymorphisms are sequence differences between the probe and intended target genomic sequence. They may impact the affinity by which a probe binds to a genomic sequence, resulting in distinct signal intensities for the same marker allele (e.g. *B* and *b* for high and low intensity respectively). Thus, individuals with an alternate allele(s) at these secondary polymorphisms have distinct clustering patterns (e.g. *Ab* in addition to *AB)*. As individuals may differ for their secondary polymorphism, this gives raise to multiple sub-clusters for the heterozygous genotypes. The location of the *Ab* and *aB* cluster moves towards the *AA* and *BB* homozygous clusters, respectively, with decreasing intensity of the *a* and *b* alleles, which may impact cluster separation and calling accuracy. Secondary polymorphisms may also lead to so called null alleles, where probes completely or nearly completely fail to bind to some of the target genomic sequences [46,47]. When not accounted for, null alleles may lead to unexpected genotype classes in segregating progenies and thereby in detected, but false, Mendelian inconsistent errors [6,9,29]. Duplicated and paralogous sequences may affect cluster separation too because they may also bind to probes, often reducing and compacting the effective cluster space for the target polymorphism [48].

Platforms may differ in their sensitivity to secondary polymorphism and duplicated sequences due to differences in chemistry by each platform [49-51]. The resulting probe hybridization data may also be interpreted differently due to differences in allele calling algorithms used in different genotyping software. The Infinium and Axiom SNP platforms make use of a selective, locus specific primer/probe that runs up to the targeted SNP. The platforms differ on how the targeted SNP is captured. The Infinium platform is based on a polymerase executed extension reaction building in fluorescently labelled nucleotides [49] whereas the Axiom platform is based on an end-to-end hybridization between a locus specific and a dye-labelled and allele specific, but otherwise non-selective probe [51]. In addition, the locus specific probes are 50-mers with the Infinium and 35-mers with the Axiom platform. Furthermore, each Infinium probe is bound in multiple copies to 20-30 beads that are in solution [52], while with Axiom the locus specific probes are located at two defined positions on a two-dimensional substrate (while the non-selective allele specific probe is in solution) [51]. How these details affect calling concordance and SNP call accuracy has yet to be revealed.

This study seeks to qualify and quantify the compatibility of allele calls from both the Illumina Infinium 20K and Affymetrix Axiom 480K apple SNP arrays for the creation of an integrated SNP dataset. We also determine reasons for inaccurate SNP calls that resulted in observed discordant SNP calls and probe failures in order to improve SNP array data interpretation and probe/platform development. Towards these goals, this study included classical concordance evaluations across individuals genotyped on both platforms, as well as accuracy evaluations by detecting and evaluating Mendelian inconsistent and consistent errors across pedigrees on a mixed dataset. We hereby updated the apple integrated genetic linkage (iGL) map [15] and used a subset of SNPs that showed high performance on the Illumina platform as defined in this paper.

## Results

### Genetic map and Infinium data curation

There were 10,295 SNPs that passed the Infinium SNP data curation steps and thus were included in the genetic map. Of these, 94.1% (9,685) were SNPs retained from the iGL map (Table 1; Additional file 1). For 12.4% (1,206) of the SNPs retained from the iGL map, new positions were assigned, and these new positions were all within their respective genetic bins on the iGL, and also within a single centimorgan (cM) of their original position except for six SNPs (Additional file 2), and except for the first 5-6Mb of LG1, where SNPs were ordered according to other ongoing studies on the GDDH13 and HFTH1 whole genome sequences (WGSs) (Van de Weg, personal communication).

**Table 1.**
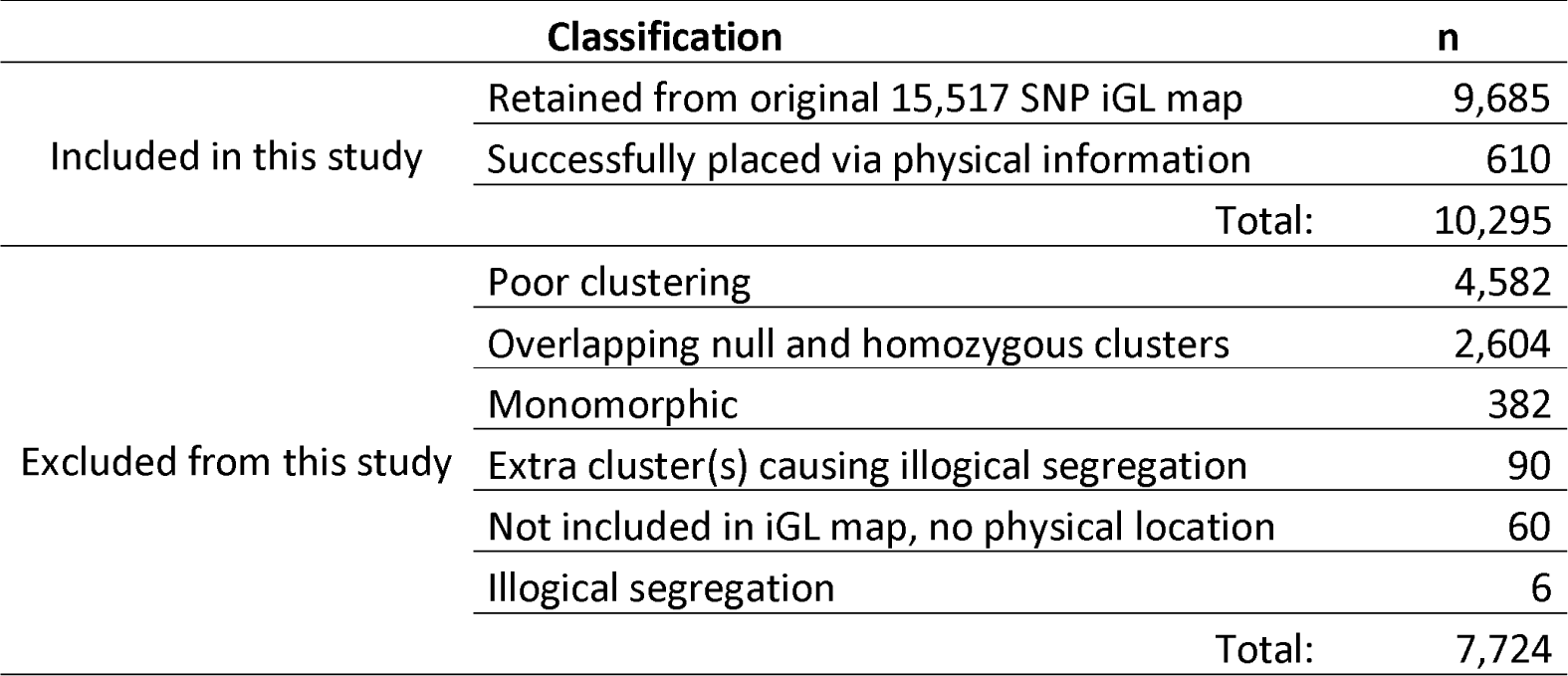
SNP inclusion/exclusion summary from the Illumina Infinium 20K array.

|  | Classification | n |
| --- | --- | --- |
| Included in this study | Retained from original 15,517 SNP iGL map | 9,685 |
|  | Successfully placed via physical information | 610 |
|  | Total: | 10,295 |
| Excluded from this study | Poor clustering | 4,582 |
|  | Overlapping null and homozygous clusters | 2,604 |
|  | Monomorphic | 382 |
|  | Extra cluster(s) causing illogical segregation | 90 |
|  | Not included in iGL map, no physical location | 60 |
|  | Illogical segregation | 6 |
|  | Total: | 7,724 |

The level by which co-segregation patterns could be examined varied per SNP and some SNPs were only polymorphic in a small number of individuals. For example, numerous SNPs were only polymorphic in *Malus floribunda* 821 and a small number of its descendants, particularly its grandchild F2-26829-2-2, which was included in the discovery panel used to create the Illumina Infinium 20K SNP array [3] and which served as a bottleneck in the introgression of the *Rvi6* gene for scab resistance from *Malus floribunda* 821 [53]. Information about minor allele frequencies (MAF) for the 10,295 SNPs included in this study across all individuals except genetic duplicates, can be found in Additional file 1.

### Evaluation of within-platform repeatability

Repeatability of Infinium data was very high, with only 0.0016% and 0.0014% average discordant SNP calls observed per genotyped duplicate when evaluating the 10,295 SNPs included in this study (Subset 1), and when evaluating only 8,412 SNPs that were also concordant in Axiom data (Subset 2), respectively (Table 2). For Axiom data, rates of discordant SNP calls varied among SNP subsets between 0.0117% and 0.3199% and were always higher than those for the Infinium data. Logically, discordancy was least for SNP sets that were first filtered for their performance on the Axiom array. Here, discordancy increased with increasing size of the subset (and thus with decreasing filtering intensity) from 0.0117% for the 253,095 SNP of subset 4 to 0.1516% for the 402,714 SNP of subset 6 (Table 2). Discordancy in Axiom data was highest for the SNPs that passed the Infinium data curation steps (Subset 1) (Table 2).

**Table 2.** Frequency of discordant SNP calls across 16 individuals genotyped twice on each array.

| Array | SNP subset Groups and their percentage (%) discordant SNP calls |  |  |  |  |  |
| --- | --- | --- | --- | --- | --- | --- |
|  | 1 | 2 | 3 | 4 | 5 | 6 |
| Infinium | 0.0016 | 0.0014 | - | - | - | - |
| Axiom | 0.3199 | 0.0706 | 0.0134 | 0.0117 | 0.1516 | 0.0421 |

\*For accessions that were genotyped more than twice on an array, a single average value was used in the across accession analyses.
Subset 1: 10,295 SNPs from the Infinium array that passed the SNP data curation steps in the present study.
Subset 2: 8,412 SNPs considered compatible between the array platforms in this study.
Subset 3: 275,223 SNPs in the Axiom array deemed robust [34].
Subset 4: 253,095 SNPs from subset 3 filtered for absence of more than one Mendelian inconsistent error in Muranty et al. [32].
Subset 5: 402,714 SNPs in the Axiom array that were classified as "Poly High Resolution" or "No Minor Homozygous" by Axiom Analysis Suite software.
Subset 6: 320,761 SNPs from subset 5 with SNPs removed that had Mendelian inconsistent errors in two or more parent-offspring relationships, discordant SNP calls in two or more duplicate pairs, or were heterozygous in doubled haploid accessions from Muranty et al. [32].

### Compatibility of Axiom data with included Infinium SNPs

The 417 individuals genotyped on both platforms (Additional file 3) showed an average concordance level of 97.1% across all 10,295 included SNPs, with a minimum of 96.0% and a maximum of 98.1%. Of the 10,295 included SNPs, 65% (6,691) had no discordant call(s), while 35% had. Following the initial process of reclustering of SNPs with discordant SNP calls that were salvageable, all SNPs just resolved and concordant were again evaluated for Mendelian inconsistent and consistent errors across all germplasm to assess accuracy. Finally, this iterative process resulted in 8,412 (81.7%) SNPs deemed compatible. These compatible SNPs were classified into nine groups based on the type of adjustment needed to make the SNP compatible (Table 3). Examples of each classification can be found in Additional file 4 and examples of each classification for SNPs deemed discordant in Axiom data can be found in Additional file 5. Classifications per SNP are included in Additional file 1.

**Table 3.**
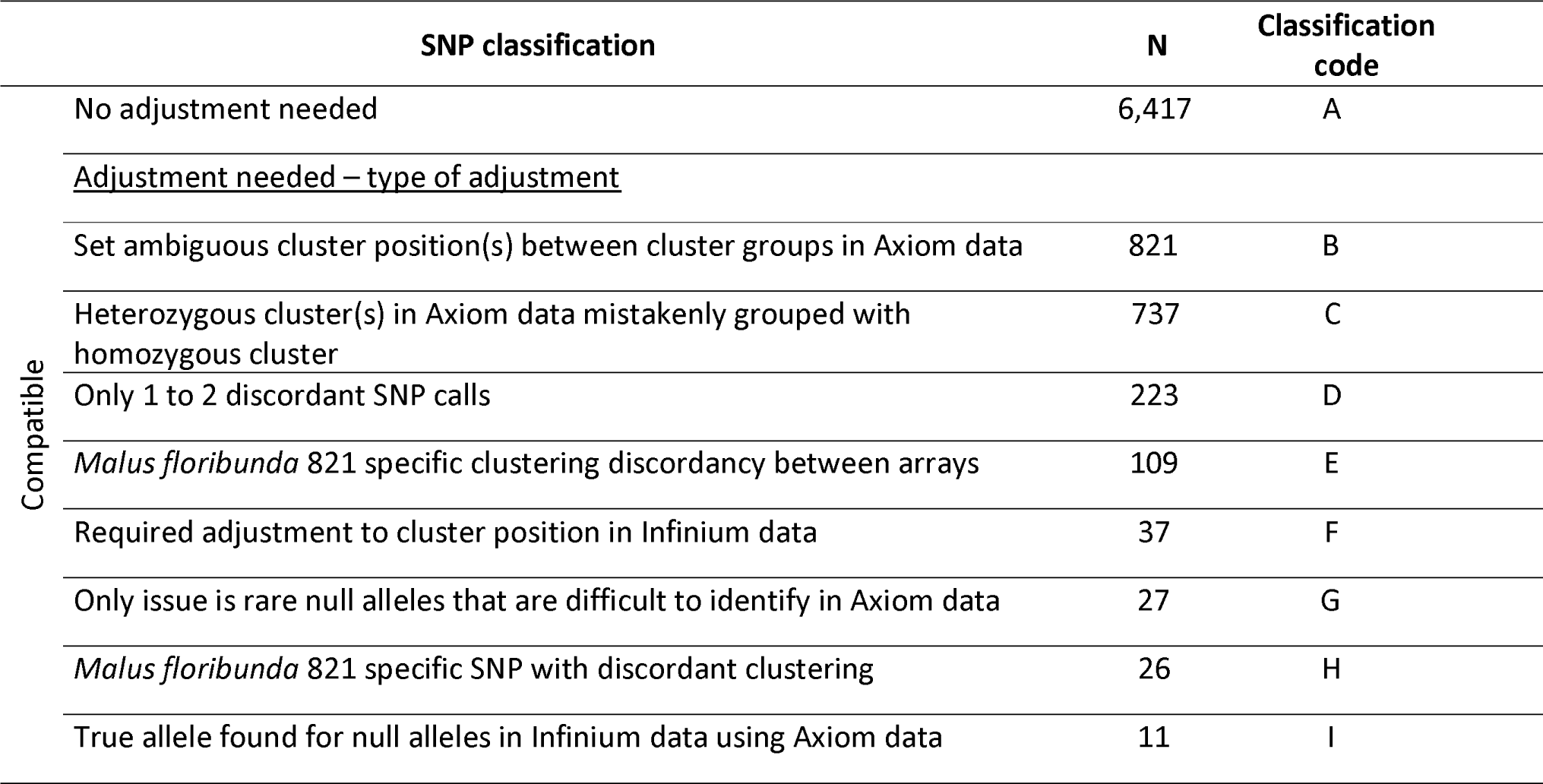
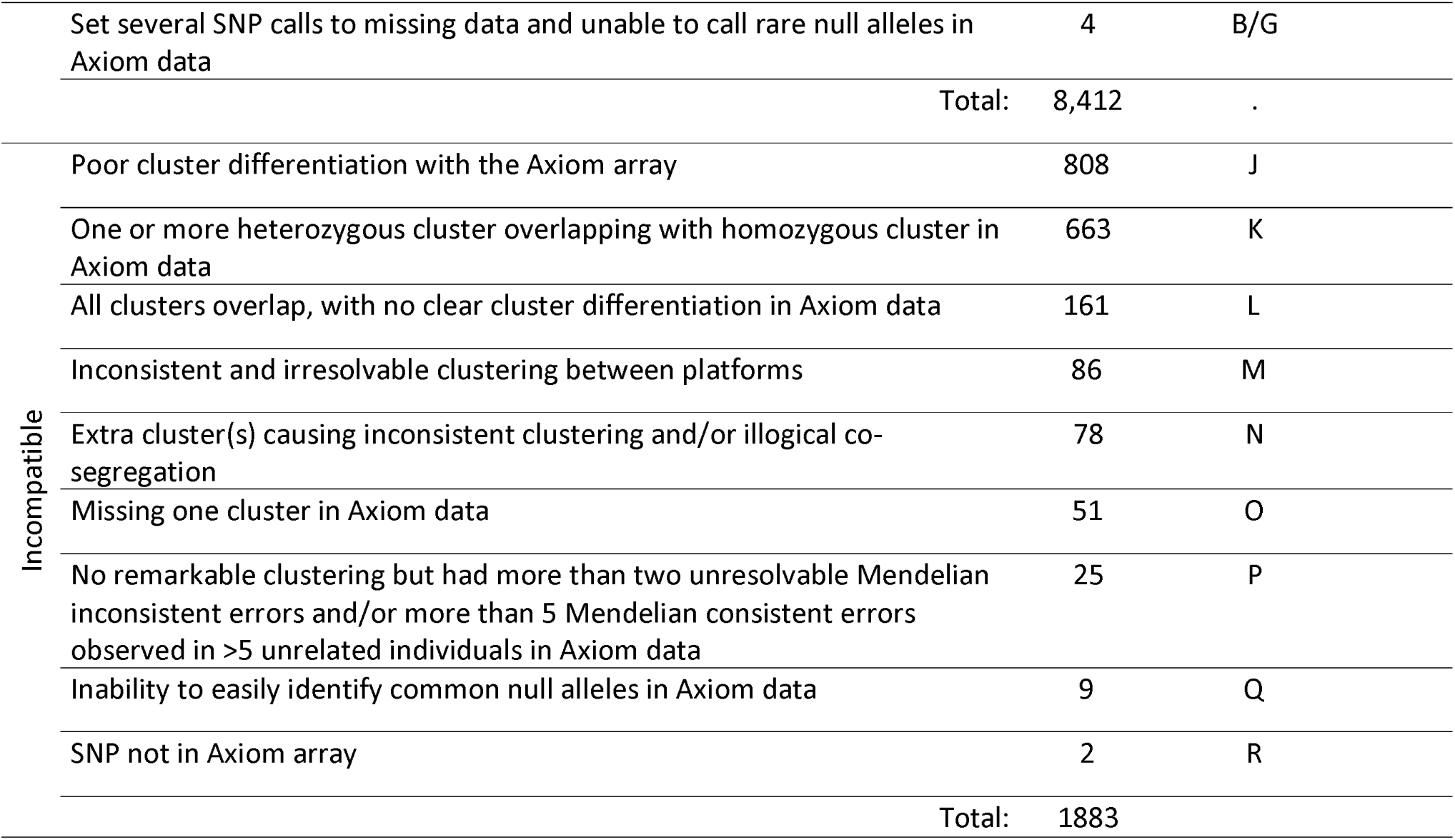
Distributions of SNPs included in the genetic map study grouped by compatible and incompatible classifications.

Of the 8,412 compatible SNPs, 6,417 (76.3%) required no additional adjustments, whereas 1,995 (23.7%) did. The most common adjustment, class B, was to set errant Axiom SNP calls between clusters to missing data. The next most common adjustment, class C, was where a subset of true heterozygous individuals was called homozygous in the Axiom data. This classification included instances where an entire or only a part of an additional cluster was originally misclassified as homozygous. Following these more common classifications, a number of less common classifications were defined. In class D there were 223 SNPs with only one or two discordant SNP calls. Of these, 70 were found with the cultivar Alice or one of two seedlings, GDxRenetta_67 and FuPi_036, whereas the remaining instances were present in smaller rates among the other individuals. The inaccurate SNP call was often an outlying location on the cluster plot (D-2 of Additional file 4). In instances where the inaccurate SNP call was unable to be determined through marker co-segregation patterns or from one being a clear outlier, the discordant SNP call was set to missing data, rather than remove a SNP for only one or two unresolved discordances. Next, in class E, 108 SNPs showed clustering differences only in *Malus floribunda* 821 and its descendants. Generally, these SNPs had logical segregation in the Infinium data, but did not in the Axiom data. Next, in class F, 37 SNPs required minor adjustments to the Infinium data where they showed a small separate cluster that was apart from the heterozygous cluster and that had to be recalled to the closest homozygous class. These clusters also always involved a group of likely related individuals, often old German or French cultivars. In class G there were 27 SNPs where the heterozygous null individuals (*AN, BN*) did not typically cluster distinctly from the true homozygous individuals in the Axiom data, whereas they did in the Infinium data. The presence of rare null alleles in Axiom data could occasionally be indirectly identified through Mendelian inconsistent and consistent errors from pedigree information, where one individual in a parent-offspring relationship was clustered in one homozygous cluster and the other individual in the opposite homozygous cluster. Class H had 26 SNPs with differential clustering and whose rare alleles were only present in *Malus floribunda* 821 and its descendants. The allele calls in Axiom data needed to be adjusted for these SNPs. Finally, class I had 11 SNPs with null alleles present in Infinium data where the true allele for the null allele could be determined from clustering from Axiom data. In the example case (I in Additional file 4), the null allele in the Infinium data was due to a secondary polymorphism two bp from the probe’s 3’ end being in coupling with some instances of the intended *A* allele but this secondary polymorphism apparently did not negatively impact clustering of Axiom data.

Another issue very rarely observed was differences in clustering between Axiom plates. The 17^th^ plate from Muranty et al. [32] had cluster positions that deviated from those from other plates. This deviation generally caused additional errant positions and lower average “size” values in some cluster plots. This issue very rarely resulted in the presence of additional clusters causing discordant SNP calls (as in H-1 and H-2 in Additional file 4), however this was sometimes the cause for several SNP calls of some of the SNPs needing to be set to missing data to make the SNPs compatible with the Infinium array (Classification B in Table 3). We didn’t observe any instances where this issue resulted in the false merging of otherwise distinct clusters. In case this issue would have been noticed at an earlier stage, we would have called this plate separately.

### Incompatibility between Infinium and Axiom platforms

There were 1,883 SNPs where Axiom data was deemed incompatible with included Infinium data. They were classified into nine different classes (J-R) (Table 3). Examples of each classification can be found in Additional file 5. Classifications per SNP are included in Additional file 1. The most common issue was poor clustering that resulted in an inability to make accurate SNP calls (class J). The next most common issue was overlapping of the heterozygous cluster with one of the homozygous clusters (class K). These are similar to class C of the compatible SNPs (“Heterozygous cluster(s) mistakenly grouped with homozygous cluster”), but the difference was that in incompatible SNPs, the boundaries of the clusters were not clear enough to manually recluster. Therefore, these SNPs could not be accurately called with the Axiom array.

The remaining classifications each comprised less than ten percent of the total number of incompatible SNPs. There were 161 SNPs that had no clear clustering in the Axiom data (mono or dimorphic) but had clear clustering in the Infinium array (Class L). Next, 68 SNPs had unresolvable discordant clustering between both arrays involving three or more individuals (class M). The example SNP provided also had many instances of unresolved Mendelian consistent errors in Axiom data. There were 78 SNPs with one or more extra clusters in both data sets, which resulted in unresolvable clustering between Axiom and Infinium data and caused multiple false Mendelian consistent errors (class N). There were 51 SNPs that were missing one cluster in the 480K array (class O). Next, 25 SNPs with illogical segregation patterns involving more than five unrelated individuals while not showing a remarkable clustering pattern (class P). There were nine SNPs with common null alleles that could be easily identified in the Infinium data but not in Axiom data. Finally, two SNPs were in the Infinium data but not in the Axiom data (Class R).

### The effects of non-target BLAST results on SNP exclusion and incompatibility

In the following, we traced causes for SNP exclusion from Infinium data and incompatibility of Axiom data using information from the GDDH13 WGS. Hence, SNPs have only been evaluated here where probe sequence were retrieved on the expected chromosome according to the iGL linkage map, which failed for 220 (2.1%) of the 10,295 included SNPs and for 549 (7.1%) of the 7,724 excluded SNPs (Additional file 1). Of the retrieved probes, 96.0% and 90.8 % gave a perfect match with the included and excluded SNPs, respectively and 0.24% and 0.64% gave highly dissimilar sequences with Expect (E) values higher than 1.E-12. Of the 8,412 compatible and 1,883 incompatible SNPs, 97.0% and 91.9% showed a full match, respectively, and 0.08% and 0.63% with E-values higher than 1.E-12. Hence, included or compatible SNPs had higher sequence similarity, and excluded or incompatible SNPs had lower similarity as estimated by E-values (Fig. 1).

**Fig. 1.**
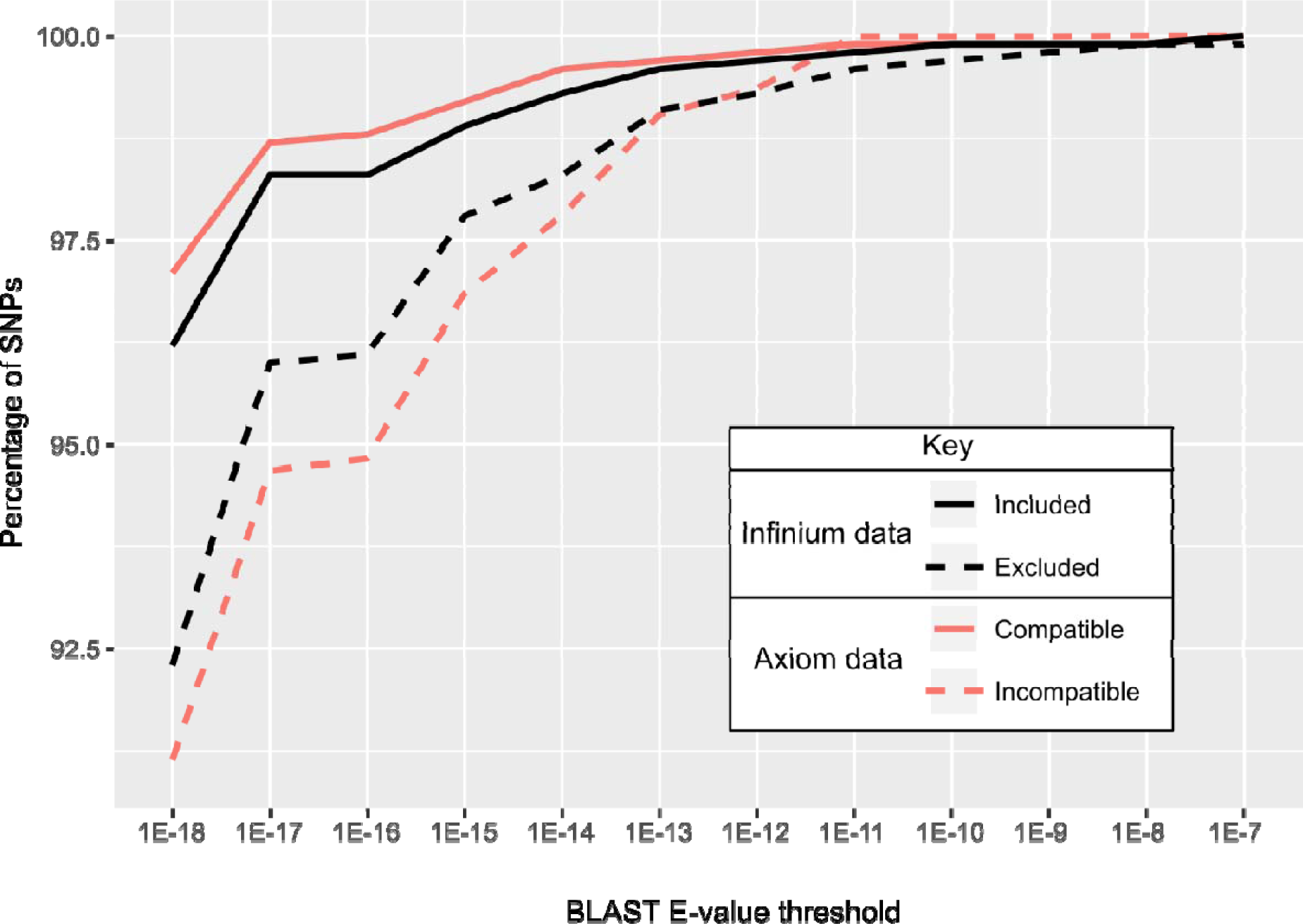
Cumulative distribution for probe E-value for probes vs. the GDDH13 WGS. Only SNPs with a significant BLAST result on the expected chromosome were considered (N=17,250).

Increasingly lower inclusion rates with Infinium data were correlated with increased numbers of BLAST results beyond a single BLAST result (Fig. 2). This was true for each of the three different stringencies on sequence similarities (E-value thresholds) for a successful BLAST result. Inclusion rates decreased from 66% with just a single hit to 6% with more than 10 hits with the least stringent BLAST result. The results for all three E-value thresholds had this same general trend. Compatibility was slightly sensitive to the number of BLAST results, with greater than four BLAST results reducing the compatibility rate (Additional file 6). However, this trend was only observed across a small number of SNPs, as only few SNPs with many BLAST results passed the Infinium data curation step. Examples of clustering likely being impaired by additional probe binding sites can be found in J-1 and J-2 of Additional file 5 and 1-1 and 1-2 of Additional file 7. However, additional highly significant BLAST results were not always associated with problematic clustering (1-3 in Additional file 7).

**Fig. 2.**
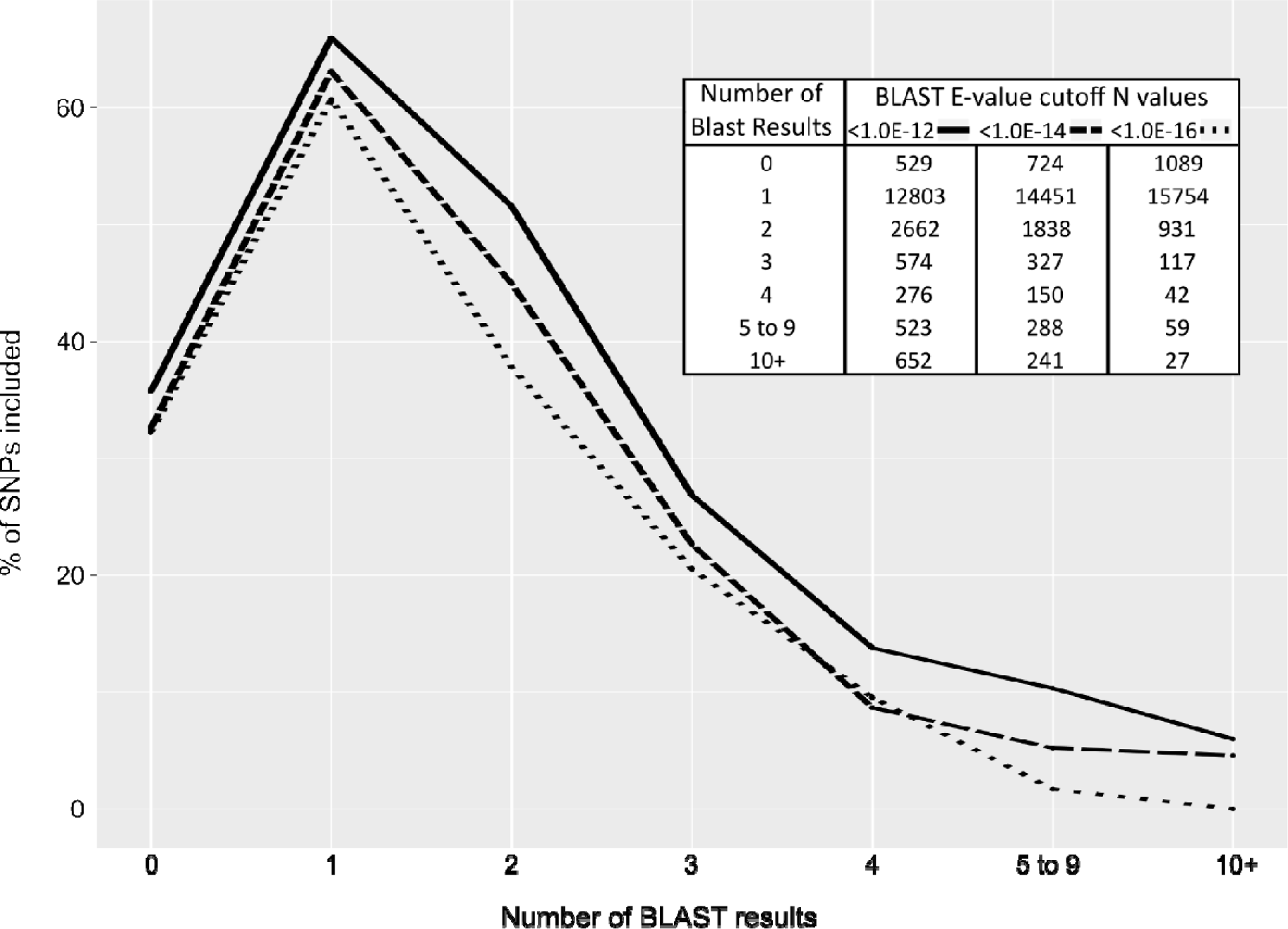
SNP Inclusion percentages vs. the number of BLAST results on the GDDH13 WGS. All 18,019 SNPs of the 20K Infinium array were considered. Three different stringency thresholds for a successful BLAST result were used: 1E-12, 1E-14, and 1E-16. The number of SNPs within each group is listed in the included table. Higher numbers of BLAST results were grouped together because of the diminishing number of SNPs that had higher numbers of BLAST results.

Included SNPs generally had a higher available cluster space than excluded SNPs, regardless of the number of BLAST results per SNP (Fig. 3). Available cluster space also decreased with increasing numbers of BLAST results and decreasing BLAST E-value cutoffs. This decrease was least among included SNPs and strongest with the more stringent threshold with both included and excluded markers. SNPs were still included in the presence of three BLAST hits with the lowest E-value cutoff but never were successful in the presence of four such hits.

**Fig. 3.**
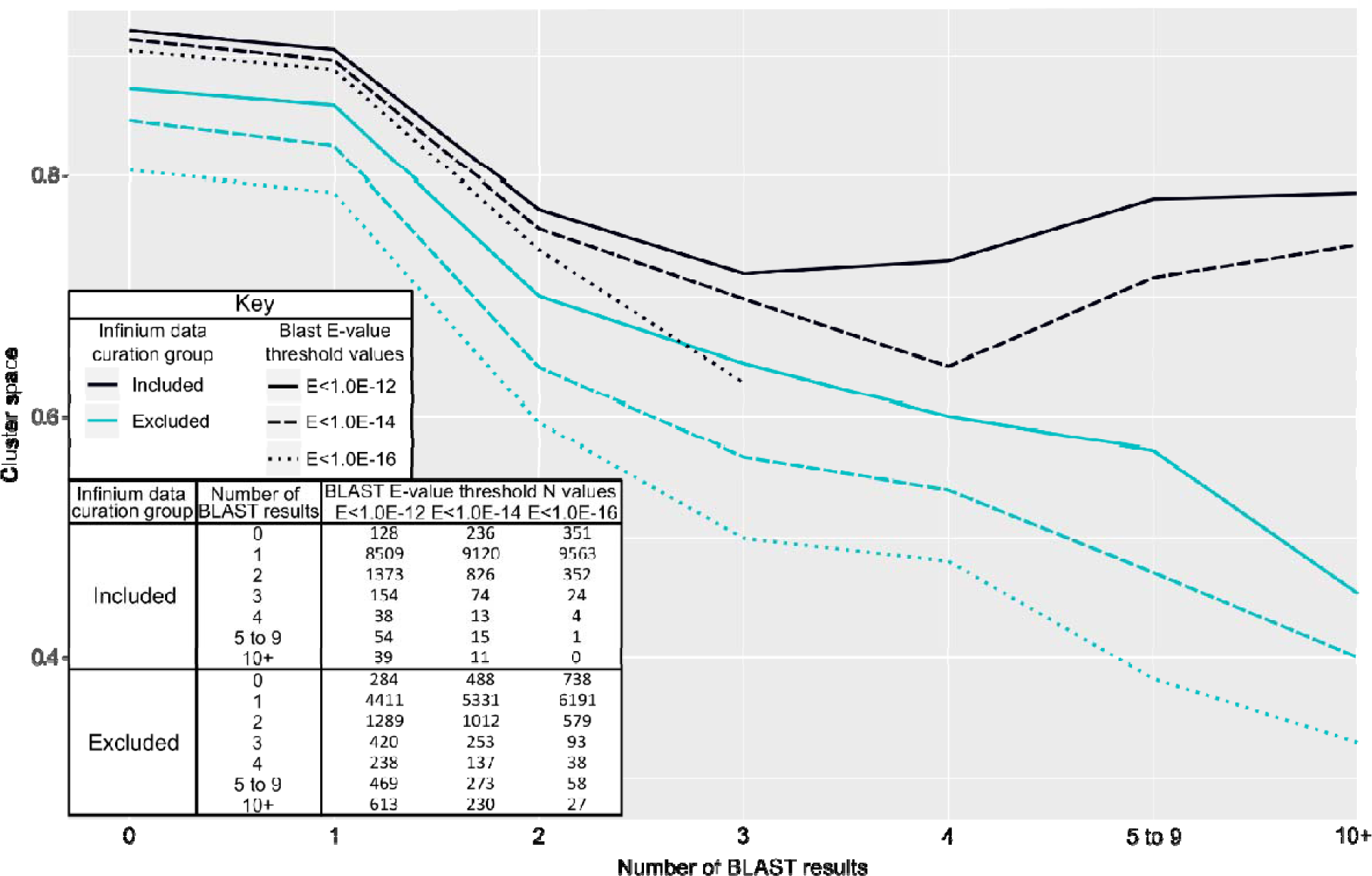
Infinium array cluster space versus the number of BLAST results on the GDDH13 WGS. Three different E-value cutoff values were used: E<1E-12 (solid line), E<1E-14 (dashed line), and E<1E-16 (dotted line). Data points were excluded from the figure if they were comprised of fewer than 10 SNPs.

### The effects of secondary polymorphisms on SNP exclusion and incompatibility

The presence of secondary polymorphism(s) at probe sites negatively impacted SNP inclusion of Infinium data and SNP call compatibility of included SNPs in Axiom data. Increasing numbers of secondary polymorphisms were correlated with reduced SNP inclusion and concordance rates (Additional file 8). We could not effectively compare simultaneously the effects both the number of secondary polymorphisms and their variable positions had on SNP inclusion and compatibility rates, so we instead focused on SNPs that had only a single secondary polymorphism to the intended target genomic sequences. The closer a single secondary polymorphism was to the target SNP, the more likely this SNP was to be excluded during the SNP curation process in Infinium data or to be deemed discordant (Classification C) (Fig. 4). In this analysis, we included secondary polymorphism occurring in at least ten percent of sequenced individuals from germplasm group 5. Probes with secondary polymorphisms within the first three positions from the target SNP were mostly excluded, due to which too few of them remained to effectively examine their compatibility with Axiom data. Because of this, we also examined Axiom cluster plots for these SNPs and they too had the poor clustering observed in the Infinium cluster plots. Among included SNPs, the presence of secondary polymorphisms also frequently resulted in the presence of additional heterozygous cluster(s) that were mistakenly called as homozygous in Axiom data, requiring manual cluster adjustment to achieve compatibility (Table 3 - class C; Fig. 4). This effect gradually diminished with increasing distance between the secondary polymorphisms and the 3’-ends of the probes (Fig. 4).

**Fig. 4.**
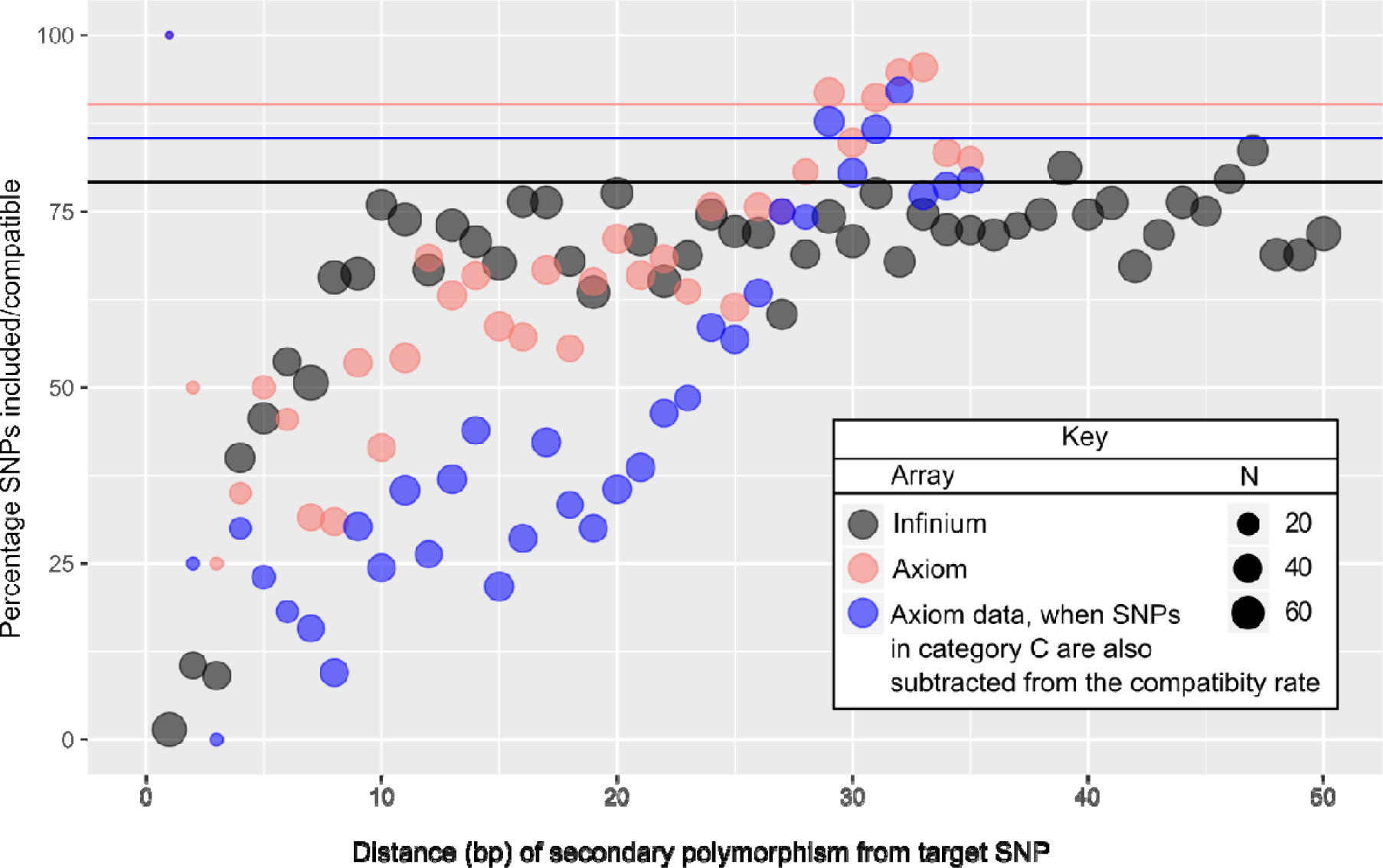
Relationship between the position of a single secondary polymorphism and inclusion and compatibility rate. The inclusion rate of Infinium data is represented by black and the compatibility of these included SNPs with Axiom data with and without class C SNPs (those with additional heterozygous cluster(s) in Axiom cluster plots requiring manual adjustment to make compatible) being classified as compatible are represented by pink and blue, respectively. The horizontal lines represent the inclusion and compatibility rates for SNPs with no identified secondary polymorphisms at their probe site for the three respective data sources that sized 6,632 (black), 6,011 (pink), and 6,011 (blue) SNPs. SNPs included in this analysis had their alternate allele present in at least 10% of the sequenced individuals, had no more than 25% missing data across the sequenced individuals, and had probe sequence with a single BLAST result on the GDDH13 WGS with an E-value <1E-12.

Automated clustering of Infinium data in GenomeStudio was assessed for 187 included SNPs that had a single secondary polymorphism and that required manual cluster adjustment in Axiom data to determine whether these SNPs had also required, or would benefit from, manual cluster adjustment in Infinium data as well. The presence of additional heterozygous clusters was often still observed in Infinium cluster plots (ex. C-3 in Additional file 4), but these additional clusters were almost always correctly called as heterozygous. There were only five cases (2.7%) where the additional heterozygous cluster(s) were set mostly or completely as missing data and only a single case (0.5%) where manual cluster adjustment was also necessary for the Infinium cluster plot to make the SNP data accurate (2-1 and 2-2 in Additional file 7).

Some secondary polymorphisms were identified as likely causes for discordant SNP calls that could be resolved through manual adjustments to Axiom cluster plots (Table 3 – class C; ex. C-1 – C-4 in Additional file 4), or to Infinium data (Table 3 – Class I; ex. I in Additional file 4), or to both platforms’ cluster plots (2-1 and 2-2 in Additional file 7). Others were identified as the likely cause of SNP discordancy (ex. J, K, M, N, O, and Q in Additional file 5), or causing probe failures or poor clustering in both arrays (3 in Additional file 7). There were also occasional instances where a secondary polymorphism possibly caused probe failure in Infinium data but not in Axiom data (ex. 4 in Additional file 7). Finally, some instances were identified where prevalent secondary polymorphisms were close to the target SNPs that lacked problematic clustering (ex. 5-1 and 5-2 in Additional file 7).

The inclusion and concordance of SNPs whose probes had BLAST E-values greater than 1.0E-12 depended on multiple different factors (Additional file 9). In general, for SNPs whose BLAST results were identical in both the GDDH13 and HFTH1 WGSs, the differences between the probe and target sequences for these SNPs were mostly present in the middle or the 5’ end of the probe sequences, though there were five SNPs (12.5% of those evaluated) with apparent mismatches within the first 10 positions from the target SNP. The observed sequence differences between probes and the GDDH13 WGS seem to be almost exclusively true, as both alternative sequences were mostly confirmed by re-sequencing data from individuals in germplasm group 5, or by the HFTH1 WGS [54]. Only occasional SNPs were identified where a likely error in the GDDH13 WGS was the reason for the low E-value observed in the BLAST data we generated. An example on true major sequence differences is probe SNP_FB_0514806 (Infinium name, with the Axiom name AX-115193385), which had a six-nucleotide gap at position 25-31 and a mismatch at position 41 from the target SNP on the GDDH13 WGS, but had a full match on the HFTH1 WGS [54]. This SNP was included and compatible without needing any adjustments to the Axiom cluster plot (Classification A), despite the substantial difference between the probe and target genomic position on the GDDH13 WGS.

## Discussion

Our results demonstrate a high degree of SNP compatibility (81.7%) between the Infinium^®^ 20K SNP array and the Affymetrix Axiom^®^ Apple 480K SNP array across 10,295 SNPs we deemed robust in Infinium data (Table 3). Our initial SNP call concordance rate for duplicate genotypes across the two platforms, 97.1%, was in line with what had been previously reported for these arrays [34]. Our further efforts towards resolving discordances and excluding SNPs with inaccurate SNP calls on either array have now allowed the creation of a combined dataset including SNPs with compatibility clustering and thus will have an exceedingly low SNP call error rate, even for individuals that were only genotyped on one of the two arrays. This highly curated combined dataset will be useful in subsequent studies. Additionally, we were able to identify reasons for both probe failures and discordance through a combined analysis of cluster plot data, SNP co-segregation patterns, and sequence data alongside the GDDH13 WGS [16]. Such an in-depth analysis of array concordance and accuracy has not been previously described in literature. Of the 8,412 SNPs deemed compatible, 2,030 are also present on the Illumina Infinium 8K SNP array (Additional file 1) [2], indicating prospects for data integration with that array as well.

### Genetic map and Infinium data curation

The map revision and SNP data curation steps revealed that many regions of the GDDH13 WGS [16] are still misassembled (Additional file 1), despite its major improvement over the previous ‘Golden Delicious’ genome [55], thereby confirming similar conclusions based on the comparison between the GDDH13 WGS and the iGL genetic linkage map [56]. A thorough evaluation of each of these regions is outside of the scope of this paper but is part of an ongoing study (Van de Weg, personal communication).

A set of 10,295, SNPs passed our rigorous data curation step (Table 1 and Additional file 1). They represent 57% of the initial 18,019 on the Infinium array and exclude 5,613 SNPs that were originally included in the iGL map [15]. The set is larger than the 6,849 SNPs used in QTL discovery studies on multi-parent populations in apple [14,17], thanks to our intense filtering and curation effort. Many of the excluded SNPs showed *AB* sub-clusters and/or null alleles, which made them highly informative for the purpose of linkage map creation using full-sib families [15,57], but which made them less suitable for use in a diversity set of germplasm due to an inability to accurately call null alleles across such germplasm.

### Evaluation of within-platform repeatability

Overall, average within-platform repeatability of genetic duplicates was high, ranging from 99.9984% to 99.9986% for the Infinium array and from 99.6801% to 99.9883% on the Axiom array, depending on the SNP subset used (Table 2). This high repeatability is in line with that observed for other Illumina arrays, such as that for *Zea mays* (99.7076%) [58], *Medicago sativa* (100%) [59], and *Populus nigra* (100%) [60], and for other Axiom arrays, such as that for *Glycine max* (>99%) [61], *Cicer arietinum* (> 99%) [62], and *Juglans regia* (99.2%) [63]. Although in this study the Infinium platform had a higher repeatability than the Axiom platform, this may not be an equal comparison, as the calculated repeatability rate was highly affected by our intense preliminary filtering process performed on the Infinium data.

It should be noted too that a very minor level of discordancy could be due to minute mutation differences between clones held in different collections. However, the chance that a SNP coincides with a mutation between clones is thought to be rare because mutation rates are usually low, and regions of mutation are usually small. Therefore, we assumed discordant SNP calls to have been entirely due to cluster position variance within each platform. Some level of discordancy could be attributed to laboratory or sampling issues. Low quality DNA has been the reason for high levels of discordancy among technical replicates in the Axiom *J. regia* 700K SNP array [63].

### Compatibility of SNP clustering between platforms

The SNPs on the Infinium 20K apple array had all been consciously included in the Axiom array to maximize the potential for cross-platform compatibility, even when Axiom’s selection criteria for array inclusion or calling performance were violated [34]. Consequently, the results of this study should not be used to determine whether the two platforms differ in their overall performance, meaning that the results and interpretations are not meant to be an endorsement or a repudiation of either platform. Despite the differences between the platforms, our results show a high degree of concordance (97.1%) for duplicate genotypes across the 10,295 SNPs deemed to have robust performance on the Infinium array. This rate was in line with the 98% previously observed [34], which evaluated a subset of the duplicate pairs evaluated in this study. This rate is also comparable to the 98.8% for humans [37], the 97.38% observed for sheep [38], and the 98.4% for swine [39].

However, despite this high level of concordance, many SNPs showed significant clustering differences between the platforms, resulting in the need for additional work to maximize compatibility. Of the 10,295 SNPs included from the Infinium array, 19% required adjustments to make them compatible and 18% were deemed incompatible with Axiom data. Researchers using data from both arrays could try to avoid issues by only using the 6,417 SNPs that did not require any additional adjustments, though this would result in either a smaller number of SNPs available or a higher level of missing data present in the combined dataset, which we wished to avoid. However, our manual curation of 1,995 cluster plots was very time consuming. The most common manual curation performed was to set some percentage of individuals to missing data in the Axiom data due to their errant positions between clusters in cluster plots (B and D in Table 3 and Additional file 4). This issue occurred in 52% of the 1,995 SNPs requiring manual curation to make them compatible and could possibly have been greatly reduced or largely eliminated by using more stringent threshold settings in the Axiom Analysis Suite [64]. Though this might have also resulted in additional missing data, it would have been a time saving step that would have possibly still retained the majority of accurate SNP calls for the relevant SNPs. The other SNPs requiring manual curation required more advanced analyses to identify and may not be as amenable to automated adjustment. They generally required either manual reassignment of sub-clusters (39%; C and F in Table 3 and Additional file 4) or the addressing of issues related to null alleles (2%; G and I in Table 3 and Additional file 4). The latter largely required pedigree information to identify the needed adjustment and correct and it was time consuming to do so. Hence, one needs to decide if it is worth one’s time to make this effort. However, in the case of the current two arrays, problematic SNPs have now been identified and solutions have been validated using a very diverse set of germplasm (Additional file 1), thereby providing a reference for future data integration efforts involving these arrays.

### Effects of non-target BLAST results on inclusion and compatibility

In our study, the effect of the overall sequence (dis)similarity between probes and intended genomic sequences on SNP inclusion and compatibility was examined. Hereto, all the SNPs’ BLAST results that met the various E-value thresholds were compared together regardless of where sequence differences occurred. This was a conscious decision, as the nature of the differences between intended and non-intended targets was often complex and involved multiple mismatches, making it difficult to quantitatively classify the differences in affinity at a more detailed level.

As much as 94.1% of the probe sequences had a full match to the GDDH13 WGS. This high proportion is not surprising, considering the full genetic relatedness between GDDH13 and ‘Golden Delicious’ [16], the source for the GDv1 and GDv2 WGSs that were used to design the Infinium and Axiom probes, respectively [3,34]. The remaining 5.9% of probes may come from incompleteness of the GDDH13 WGS, from local sequence errors in the GDv2 or GDDH13 WGSs, or from sequence differences between the two ‘Golden Delicious’ homologs. The former two reasons lead to misleading high E-values, whereas the latter results in true secondary polymorphisms and indels in part of the germplasm that may raise issues in clustering. Hence, it is not surprising that full matches were more prevalent in included and in compatible SNPs than in excluded and in-compatible SNPs (Fig. 1). Other reasons for exclusion and incompatibility was correlated with the number of BLAST results. Many SNPs had two BLAST results, usually with the non-intended target having a higher E-value and often appearing on the homoeologous chromosomes from the recent whole genome duplication in *Malus* [55]. Other SNPs had more than two significant BLAST results.

Probes with multiple BLAST results were more likely to be excluded (Fig. 2) because of a loss in available cluster space (Fig. 3). The binding of a probe to one or more of these non-intended targets results in additional signal, typically for only one of the two possible marker alleles, whichever is present at the non-intended site(s). This causes the heterozygous and one of the homozygous clusters to shift towards the allele with extra representation (as in cases 1-1 and 1-2 of Additional file 7). Occasionally both homozygous clusters may shift towards each other, reducing the effective cluster space from both sites of the cluster plot. This happens in the presence of multiple effective hits that produce additional signal for each of the two possible marker alleles. These observations are in line with Hyten et al. [48], which previously reported a reduction in cluster space due to paralogous binding sites. These results highlight the importance of a high-quality reference genome and adequate SNP filtering for array design.

Though SNPs whose probes had more BLAST results were more likely to be removed during the SNP data curation steps and to be deemed discordant, many were both included and deemed compatible in Infinium and Axiom data, respectively (e.g. 1-3 of Additional file 7). This could be due to the presence of just one segregating locus that did not suffer from additional complicating issues like secondary polymorphisms of major effect, due to which clusters remained well separated despite a reduced effective cluster space, the non-intended BLAST result(s) having no or weak binding affinity to the probes, or part of the BLAST hits being false due to remaining assembly errors in the GDDH13 WGS (see below).

### Effects of secondary polymorphisms on inclusion and compatibility

Secondary polymorphism make a probe differ from its target genomic sequence: the closer a secondary polymorphism was located to the 3’ end of the probe, the more likely the probe was excluded during the Infinium data curation step, deemed incompatible with Axiom data, or require manual adjustment to Axiom cluster data to make compatible (Fig. 4). Probes with secondary polymorphisms on the first seven positions from the 3’ end had the lowest inclusion rate on Infinium data, and those excluded often had null alleles present. The increased presence of null alleles in probes with secondary polymorphisms within the 3’ portion of probes and their negative effect on concordance and accuracy has also been previously reported with Illumina human SNP arrays [46,47].

A higher impact of secondary polymorphisms on clustering was observed for Axiom data than for Infinium data (Fig. 4). This is likely due to the difference in probe size. The longer probe size in the Infinium array could result in a stabilizing effect by the 5’end for binding when the secondary polymorphism(s) exist in the middle or 3’ end of the probe. This stabilizing effect apparently allowed for successful SNP calling even when multiple SNPs or long gaps existed in the middle of the probe sequences, though in some cases this resulted in the need for some manual clustering (Additional file 9). With the smaller 35-mer Axiom probe, this stabilizing effect may be greatly reduced to approximately the final 7 nucleotides. The probable increase of stability with increasing probe size might have been the reason why the initial 30-mer oligonucleotides of the Axiom platform have been extended to the current 35-mers [51]. These results and insights could be used to guide SNP filtering for future array construction, including for decisions on probe size.

SNPs were identified with prevalent secondary polymorphisms close to the target SNP that nevertheless lacked problematic clustering (ex. 5-1 and 5-2 in Additional file 7) and there were also instances where the exact cause of poor clustering could not be identified (ex. L-2 in Additional file 5). Sequence data and the GDDH13 WGS are not perfect, and it could be that we simply did not have the available information to detect the likely cause for poor clustering in some cases. Likely, there are also other factors unaccounted for in this study, such as the exact nucleotides that are mismatching or GC content of probes.

### Array construction and Reference WGSs

Secondary polymorphism and paralogous sequences are known to negatively impact SNP array marker performance [46,47], the first causes problems with probe hybridization which might affect the probe efficiency while the second might introduce false SNPs obtained from the consensus of sequences coming from each of the two slightly different regions when erroneously merged together. Therefore, measures are undertaken to limit their occurrence in the design of a new SNP array. Following the initial SNP discovery process, candidate SNPs are checked i) for the lack of additional polymorphisms in their flanking sequence and ii) filters on the read depth at the SNP site are used as a proxy for identifying the risk of erroneous calls due to the fusion of reads from paralogous regions. The effectiveness of the additional polymorphisms filtering step is a function of the size and genetic diversity represented by the discovery panel, which consisted of 13 and 63 individuals in for the 20K Infinium and 480K Axiom array, respectively. Hence, compatibility issue rates observed for the Infinium array SNPs observed in this work are likely not representative for the other 460K SNPs included on the Axiom array since the filtering pipeline of the latter had more chances to identify additional polymorphisms in the probe due to the larger discovery panel. Another factor in play is the quality of the reference WGS used in the alignment of sequencing data of the discovery panel members. To avoid the presence of paralogous sequences, several filters can be applied, such as a check on the read depth at the SNP site and/or a kmer analysis of the probes of selected candidate SNP markers. For example, it is possible to align the entire probe (or multiple subsequences of the probe i.e. kmers) against the reference genome and make sure that these sequences do not appear multiple times across the genome. The efficiency of this approach is a function of the size of the queried segments, the allowed number of mismatches, the sensitivity of the probe for mismatches given the genotyping platform, and the quality of the reference WGS. In the design of the 20K array, the full probes (50-mers) were aligned to the reference genome allowing for two mismatches, while a kmer analysis was performed on all the 24-mers from the probes to make sure they did not appear multiple times in the genome [3]. Our results indicate this approach to be effective for the Axiom genotyping platform, but not for the Infinium where probes with more than two polymorphisms at their first 24 bps platform may still result in relevant marker signal (see examples in Additional file 9). Also, filtering against secondary polymorphisms takes out a major source for null alleles. In contrast, identification of indels through an SNP discovery panel is quite error prone when the re-sequencing data are of short read length and at moderate read depth.

Finally, the quality of the reference WGS used in the design of an array matters. The WGS used in the design of the 20K Infinium array was of a complex nature. It consisted of a pseudomolecule for each chromosome in apple plus a series of additional scaffolds, which mostly were highly similar homologous and homoeologous sequences. These scaffolds could not be distinguished for their chromosomal origin, due to which they could not be collapsed into the primary sequence. However, the recent availability of chromosome scale assemblies like the GDDH13 or that of the HFTH1 WGS [54], gives better opportunities to perform the aforementioned filters more effectively.

### The GDDH13 Whole Genome Sequence

While the GDDH13 WGS is a great improvement over the previous ‘Golden Delicious’ WGSs, several separate observations in our study point to the existence of probable assembly errors in the GDDH13 WGS. First, some probes had more than one perfect BLAST result, while the SNP array cluster plots showed signal for just one locus (1-3 in Additional file 7). Second, we observed mismatches between the GDDH13 physical positions and the genetic positions that were not due to mapping errors (see Additional file 1, column “phys_blast_GDDH13v1.1”). Finally, we identified specific Infinium probes that did not give a BLAST result on the GDDH13 WGS but resulted in regular heterozygous signal in ‘Golden Delicious’, the cultivar that served as source for the GDDH13 individual due to incompleteness in the GDDH13 WGS (Additional file 9).

## Conclusions

We identified 8,412 SNPs that could be used to construct an accurate integrated Infinium and Axiom apple SNP array dataset. Problematic clustering encountered in our study was primarily due to secondary polymorphisms, alternate probe binding locations interfering with probes’ target sequences, and differences between probes and target genomic sequences. The proximity of the sequence difference(s) to the target SNP between probes and their target and non-target sequences was identified as a major factor in SNP inclusion and compatibility. The specifics regarding these results could be used to help guide further SNP filtering for array construction.

## Methods

### Available array data

Illumina Infinium 20K SNP array data came from previous [11,15,20,27] and ongoing studies. Affymetrix Axiom 480K SNP array data came from Bianco et al. [34] completed by Muranty et al. [32]. SNP intensity data were made available on request. SNP calling procedures progressively developed during the successive stages of this work as described below. The two arrays will be referred to as the Infinium and Axiom array, respectively.

### Germplasm

Each accession used for SNP curation, the analysis of duplicates, and the analysis of SNP call concordance and accuracy in this study is listed in Additional file 3. This group of accessions was filtered for being diploid and for having high SNP call quality using methods outlined in Vanderzande et al. [29]. Each cultivar was assigned a *Malus* UNiQue genotype code (MUNQ) value as described in Muranty et al. [32].

Five general groups of germplasm were distinguished, which partly overlapped. The first group was comprised of 1,566 Infinium genotyped cultivars/accessions, some of which were included more than once as different accessions from different institutions. The second group was comprised of 1,466 individuals genotyped on the Axiom array. The third group was comprised of 30 full-sib families totalling 2,422 seedlings genotyped on the Infinium array and evaluated in various previous genetic linkage mapping and QTL discovery studies [11,15,21,22,27] (these families are listed in Additional file 10). The fourth group consisted of known pedigree ancestors of the above families (group 3) and individuals of groups 1 that were known to have direct genetic relationships. Finally, a fifth group comprised of individuals whose sequence data was used in the Axiom array SNP discovery panel [34] was used for the identification of secondary polymorphisms. Groups 1 and 2 were used to identify genetic duplicates between the two groups; these duplicates were used to evaluate concordance. Groups 3 and 4 were used for the genetic map revision, curation of Infinium SNP data, and filtering of SNPs. Groups 2, 3, and 4 were used for accuracy evaluations.

### Identification of genetic duplicates analysed on both arrays

Germplasm groups 1 and 2 (Infinium and Axiom genotyped cultivars, respectively) were used to identify genetic duplicates. SNP calls for the first group were generated using default settings in GenomeStudio^®^ v2.03 (Illumina Inc. San Diego, CA, USA). SNP calls for the second group were obtained from Muranty et al. [32]. Individuals were deemed genetic duplicates when they shared more than 97% of SNP calls from 7,206 selected SNPs that were deemed “robust” in Muranty et al. and, on the Infinium platform, not excluded from Vanderzande et al. [29], free of apparent null alleles in full-sib families from germplasm group 3, and that were included in the iGL map with single locus segregation [15]. This set of SNPs was only used for this purpose; a more thoroughly evaluated set of SNPs was used for subsequent steps. The cutoff value of 97% was established by comparing the SNP calls for the wild genotype *Malus floribunda* 821 from both platforms and rounding down to the closest whole percentage point, as *Malus floribunda* 821 was most prone to atypical clustering with each of the arrays and showed a relatively high level of discordant SNP calls compared to other expected genetic duplicates, due to its wild origin compared to the domesticated pool for which both arrays were initially built.

### Genetic map revision, curation of Infinium SNP data, and filtering of SNPs

We used a revised version of the iGL map [15] to ensure adequate evaluation of the accuracy of SNP data at the later stages of this research. The iGL map resulted from the genetic mapping of virtual haploblock markers in which segregation information of 1 to 12 tightly linked SNPs was aggregated, while the SNPs within a haploblock were ordered according to the Golden Delicious WGS version 2 [3]. We revised the SNP sorting order and cM positions for SNPs in this map using the process described in Vanderzande et al. [29] using physical coordinates obtained by blasting [65] the 50bp long probe sequences from all SNPs on the Infinium array [3] against the Golden Delicious doubled haploid v1.1 WGS (GDDH13) [16]. BLASTn parameters used are as follows: Expected Value Threshold 0.001, Word Size 11, Match/Mismatch Scores 1 and -2, Gap Costs; existence 5 and extension 2. Parameters were chosen to generate a dataset neutral to bias in detection of orthologs and paralogs [66]. To ensure the accuracy of the physical positions, results from the more extensive, 121 long probe source sequences [3] were used in a few instances where no results were found using only the 50bp probe sequences. For SNPs with multiple matches, only matches with Expect (E) values <1.0E-12, approximately equivalent to less than 4 mismatches in a 50 bp sequence, that were not in conflict with coordinates of genetically flanking SNPs were considered. When multiple matches with physical positions between flanking SNPs had E-values <1.0E-12, the match with the lowest E-value was used. SNPs were rearranged to match these physical coordinates when not resulting in spurious double recombinations in seedlings used to construct the 8K [20] and 20K iGL maps [15]. We allowed re-ordering of SNPs up to 2 cM away from their original position, rather than just within the genetically and physically more confined haploblocks. These new positions were accepted when they did not lead to false Mendelian consistent errors.

New cM position estimates were made for SNPs where rearrangements resulted in conflicts with the original cM positions. This was accomplished by using a linear algebra approach using the relative physical distances to the nearest flanking SNPs with concordant positions. Following this step, SNPs that were originally not included in the 20K iGL map were placed into the genetic map, estimating their genetic position from their identified physical position, using the same algebraic method as previously described. For rearranged SNPs that required new cM positions that had no blast hits, a cM position was assigned that was equidistant between the genetically flanking SNPs. Occasional SNPs that required rearrangement and new cM positions to address spurious double recombinations (due to incorrect SNP order) had blast hits that were inconsistent with flanking SNPs. If one of these adjacent SNPs was inconsistent with the SNP that needed a new cM position and its other flanking SNP, another adjacent SNP that had a consistent blast hit was used for cM position imputation using linear algebra. If this was not possible, a cM position that was equidistant between the adjacent SNPs was assigned instead. The ordering of SNPs for the first ∼ 6Mb of LG1, was based on unpublished data made available by Eric van de Weg.

After the construction of this preliminary updated map, all Infinium genotyped individuals that had direct parent-child relationships (listed in Additional file 3) were called for all SNPs in the updated map using the default clustering settings in GenomeStudio. Mendelian consistent and inconsistent errors were identified using GenomeStudio and FlexQTL™ [67] as outlined in Vanderzande et al. [29]. The resulting output on Mendelian errors from FlexQTL™ was used to curate the data as described in Vanderzande et al. [29] and to classify SNPs as either included or excluded. SNPs classified as included were used in later portions of our study and did not have unresolved Mendelian inconsistent errors and were not involved in numerous spurious double recombination events, or false Mendelian consistent errors. SNPs that did not meet these criteria were visually inspected to determine whether they could be addressed to be able to fit the criteria for being included or whether they should instead be excluded. Excluded SNPs generally had poor clustering or problems due to null alleles. Poor clustering was defined as a lack of clarity in differentiating between *AA, AB*, and *BB* clusters. In some cases, clustering for these SNPs could be resolved via manually adjusting cluster positions or the occasional omission of errant SNP calls. When resolution of these clustering problems was not possible, the SNP was excluded from analysis. SNPs with null alleles had individuals with cluster positions of lower intensity, or in other words, a lower “Norm R” value in GenomeStudio cluster plots compared to true homozygous clusters. These were identified via the individuals with those lower intensity cluster positions also having a parent or an offspring with a cluster position of lower intensity located below the opposite homozygous cluster. In such a parent-offspring relationship, GenomeStudio assigns one of the individuals in this relationship a SNP call of “*AA*” and the other individual a SNP call of “*BB*”, when in reality the two individuals share a third allele, termed “Null” (*N* in additional files). These null alleles may be due to either indels or additional polymorphisms on probe sequences resulting in calling problems obscuring the true alleles at the intended locus [6,9,67]. In segregating families, the presence of such alleles could be unequivocally assessed based on a-typical segregation ratios. For cultivars with known, genotyped parentages, null alleles may also generate Mendelian consistent and Mendelian inconsistent segregation errors. For cultivars for which parents are unknown or not genotyped and also lack genotyped offspring, calling was solely based on the shape of genotype clusters in GenomeStudio cluster plots. Here, in many cases, heterozygous null genotypes could not be clearly differentiated from truly homozygous genotypes. If the null allele for such SNPs was rare, the SNP was still classified as included. This distinction was made because we wished to avoid excluding SNPs if they were only problematic in a very small number of individuals, particularly those involving genetically unique or obscure individuals. SNPs were also removed if they were found to be monomorphic.

### Identification and evaluation of discordant SNPs in duplicated individuals

Only SNPs that passed the previously described curation steps on the Infinium array data were considered for evaluating discordant SNPs in duplicate individuals. SNP calls for Axiom genotyped individuals were obtained using Axiom Analysis Suite software using the default diploid settings. Discordant SNP calls were identified for all pairs of individuals that were genotyped on both arrays. They were further evaluated to determine the nature of the discordancy and whether manual cluster adjustments could be made to improve concordance. In cases of ambiguity regarding which of the discordant calls was accurate, pedigree information was used to evaluate inheritance or co-segregation using FlexQTL™. Confirmed null alleles in Infinium data were not tallied as discordant SNP calls with Axiom data, as they had been hand called during the Infinium data curation step. If manual cluster adjustment of one or both platforms could result in the resolution of the discordance, the SNP was included in the following steps. If not, the SNP was deemed discordant.

### Identification of higher order SNP data quality issues affecting concordance and accuracy

Following the previous investigation of discordance between duplicated genotypes, discordance and accuracy were investigated in a single, mixed Infinium-Axiom dataset using all germplasm included in this study. Individuals were filtered to include only those having published parent-offspring relationships (Additional file 3). This dataset was imported into FlexQTL™ to identify Mendelian inconsistent and consistent errors which were then systematically evaluated as described in Vanderzande et al. [29]. Through this evaluation SNPs were identified that had irresolvably inaccurate calls. SNPs with inaccurate Infinium data were reclassified as excluded. Following this reclassification, SNPs with inaccurate Axiom data were deemed incompatible and classified as described in the section “Classification of compatible and incompatible SNP clustering”.

### Evaluation of within-platform repeatability

Repeatability of SNP calls was evaluated for each platform using individuals that were genotyped twice or more on both platforms from separate DNA extractions (biological replicates) and using different subsets of SNPs. These subsets were 1) the SNPs that passed the Infinium data curation steps in this study, 2) SNPs considered concordant and accurate between both arrays, 3) SNPs in the Axiom array previously deemed robust [34], 4) SNPs from subset 3 that were filtered for absence of more than one pedigree relationship in Muranty et al. [32], 5) SNPs in the Axiom array that were classified as “Poly High Resolution” or “No Minor Homozygous” by Axiom Analysis Suite software, and 6) SNPs from subset 5 with SNPs removed that had Mendelian errors in two or more parent-offspring relationships, discordant SNP calls in two or more duplicate pairs, or were heterozygous in doubled haploid accessions from Muranty et al. [32]. The few available technical replicates were not considered.

### Classification of compatible and incompatible SNP clustering

During the above described Identification and evaluation of discordant SNP in duplicated individuals, SNPs deemed compatible, which we define here as having concordant and accurate SNP calls between both platforms, were classified into nine groups according to the type and prevalence of the applied manual cluster adjustments to achieve compatibility. These groups are as follows:

A. Adequate cluster differentiation, no adjustments of allele calls required.
B. Diffuse clustering in Axiom data which required setting the area(s) between cluster groups in Axiom data to missing as three or more individuals were identified as discordant and with ambiguous positions in Axiom data.
C. Two or more heterozygous sub-clusters present in the Axiom data, where one or more of these clusters was fully or partially incorrectly grouped with a homozygous cluster and where the additional cluster(s) was able to be unambiguously recalled.
D. Only one or two discordant allele calls that were made missing.
E. Discordant calls for *Malus floribunda* 821 and sometimes some of its descendants that had differential clustering between the arrays, resulting in the SNP call for these individuals being ignored in Axiom data.
F. Presence of a small heterozygous sub cluster in Infinium data located between the main heterozygous and a homozygous cluster where the allele in the sub-cluster with reduced signal has become complete invisible in the Axiom data. Could be made accurate by adjustment to cluster positions in Infinium data for a small number of individuals and was thought to be due to (disturbing) signal from an additional locus which is not present in or influencing Axiom data.
G. Low frequency (<0.5%) of null alleles in *AN* or *BN* heterozygous genotypes in the Infinium data that could not be as easily identified in the Axiom data. Null alleles were inferred from Infinium cluster plots or from segregation data in Axiom data when possible, otherwise some individuals were left with unidentified null alleles.
H. SNP that was only polymorphic in *Malus floribunda* 821 and possibly some of its descendants which had discordant SNP calls. Infinium data was retained for these individuals as Axiom data was deemed inaccurate through the evaluation of segregation data. Occasional need to readjust cluster definitions in Axiom data for non-*Malus floribunda* 821 related individuals.
I. Null alleles in the Illumina data that could easily identified through *AN* or *BN* or *NN* clusters, but not in the Axiom data, where the true allele for the probe’s target SNP was identified using Axiom and sequence data; entire Infinium (sub)clusters were recalled. SNPs deemed incompatible were classified into nine groups and organized by how prevalent each observed issue was. In our study we defined incompatibility as SNPs with Axiom SNP calls that were either irresolvably discordant with curated Infinium data or SNPs with Axiom SNP calls that were deemed inaccurate. No systematic effort was made to identify the cause of each instance of discordancy. These groups are as follows:
J. Three clusters are clearly present in Infinium data, but have poor cluster differentiation at first sight, with illogical segregation in Axiom data.
K. One or more heterozygous clusters partially overlap with homozygous cluster in Axiom data.
L. All clusters overlap, without any cluster differentiation in Axiom data.
M. Inconsistent and irresolvable clustering between platforms.
N. Presence of extra cluster(s) with irregular positions that caused inconsistent clustering or illogical segregation.
O. Missing either one homozygous cluster or the heterozygous cluster in Axiom data, while the Infinium data showed all three genotype clusters.
P. No remarkable clustering but had unresolvable Mendelian inconsistent and/or consistent errors observed in >5 unrelated individuals in Axiom data.
Q. Had null alleles present in more than five unrelated individuals that could be accurately called in Illumina data but could not be accurately called in Axiom data.
R. SNP not present in Axiom array

### Identification of reasons for SNP exclusion and incompatibility

We explored whether non-intended genomic sites that also hybridize to probes could be causing SNPs to be excluded during the initial Infinium data curation steps. All 18,019 SNPs on the Infinium array were grouped by the number of BLAST results versus the GDDH13 WGS [16] that each had, using the 50bp probe sequence from the Infinium array [3]. Next, for each group the percentage of excluded SNPs was determined. The number of BLAST results was tallied at three different E-value thresholds: <1.0E-12, <1.0E-14, and <1.0E-16. A maximum E-value cutoff of <1.0E-12 was chosen because it included intended match for over 99.3% of SNPs from the genetic map revision portion of this work.

Next, we explored whether the presence of additional non-intended genomic sites reduced cluster space available for SNP calling in Infinium cluster plots. Hereto, the effective cluster space was compared between SNPs grouped by inclusion or exclusion and by the number of BLAST results at the different E-value thresholds. This was done because a reduction in cluster space could result in poor resolution between heterozygous and homozygous clusters, which could reduce SNP call accuracy. Available cluster space was calculated for each SNP by the difference between 5% and 95% quantiles of observed Theta values. This metric was used instead of the difference between the minimum and maximum Theta values to account for occasional outliers.

We also assessed how secondary polymorphisms at the probe site were influencing SNP inclusion rates of Infinium data and SNP call compatibility rates of included SNPs in Axiom data. To accomplish this, we first identified secondary polymorphisms among 53 individuals from the 480K Axiom array SNP discovery panel [34] (Group 5 in Additional file 3) by aligning their sequence data to the GDDH13 WGS [16] using the Burrows-Wheeler Alignment tool [68] in conjunction with SAMtools [69] and identified all variants within the physical coordinates for the 50bp long Infinium probes, as identified through the previously described BLAST results. Secondary polymorphisms were considered if they were present in at least ten percent of the sequenced individuals. We used the resulting data to demonstrate how secondary polymorphisms influenced SNP calling quality in two ways. First, for each relevant (in)compatibility classification, we identified one or more representative SNPs. Next, their sequence-based polymorphism data was examined alongside cluster plot data to determine whether the observed clustering problems resulting in inaccurate and thus incompatible SNP calls were likely caused by secondary polymorphisms, as to provide explicit example cases. Second, information on the positions of secondary polymorphisms was correlated to the inclusion/exclusion rate in Infinium data and by compatibility of Axiom data with included Infinium SNPs. Additionally, to determine the effects of major differences between a probe and an intended target sequence, we also evaluated all cases of included SNPs that had probes with E-values greater than 1.0E-12. To minimise the risk of examining artefacts due to high E-values being caused by local sequence errors in the GDDH13 WGS [16], we also considered BLAST results from the HFTH1 WGS [54] for this step, which became available during this study.

## Abbreviations

SNP: Single nucleotide polymorphism
iGL map: Integrated genetic linkage map
cM: centiMorgan
WGS: Whole genome sequence
E-value: Expect values; the number of expected hits of similar quality (score) that could be found just by chance

## Acknowledgements

We would like to thank Diego Micheletti for providing aligned resequencing data and Simone Larger for performing genotyping. We also thank the following institutions for providing germplasm used in this study: Seed Savers Exchange in Decorah, Iowa, USA, Ökowerk in Emden, Germany, and NIAB EMR in Kent, UK.

## Availability of data and materials

The Axiom data used in this study was made available through Muranty et al. [32]. Curated Infinium data for duplicates and pedigree ancestors will be available at the Genome Database for *Rosaceae*. The highly curated integrated apple SNP array dataset will be shared through following publications on downstream analyses.

## Ethics declarations

### Ethics approval and consent to participate

Not applicable.

### Consent for publication

Not applicable

## Additional files

Additional file 1:

Updated iGL map for Illumina Infinium 20K SNP array.

Additional file 2:

Changes in position of SNP markers of 9685 SNPs included in this study that were also included in the iGL map [15].

Additional file 3:

Germplasm included in study. Additional file 3.1 includes only genetically unique individuals and their associated “analysis names”, pedigrees, and MUNQ values and whose SNP data was used for data curation and/or evaluating concordancy and accuracy. Some individuals genotyped on the Infinium array and whose SNP data was only used for identification of genetic duplicates in Axiom data are not listed in this table and instead are included in Additional file 3.2 contains a listing by accession name and ID and includes all individuals, including duplicates. Additional file 3.3 contains a listing of the institutions who provided germplasm listed in Additional file 3.2.

Additional file 4:

Cluster plot examples for classifications of compatible SNPs from Table 3

Additional file 5:

Cluster plot examples for classifications of incompatible SNPs from Table

Additional file 6:

Compatibility rate of Axiom data across the 10,295 SNPs that were included from the Illumina curation step grouped by the number of BLAST results per SNP. Three different BLAST E-value thresholds were considered.

Additional file 7:

Additional cluster plot examples

Additional file 8:

Number of secondary polymorphisms verses inclusion of SNPs in Infinium data and compatibility of Infinium data with Axiom data.

Additional file 9:

Information for SNPs whose probes have BLAST result E-values greater than 1.0E-12 that were still included in the Infinium data curation step and possible explanations as to why the apparent differences between their probes and target genomic positions were not sufficient to result in exclusion of the SNP and why these differences may or may not have negatively influenced the compatibility of Axiom clustering. Differences between the probe sequences and either genome sequence are listed in columns I and J as mismatches (M) and/or gaps (G).

Additional file 10:

Families used for the improvement of the 20K integrated genetic linkage map.

