## Additional file 4 for "Integration of Infinium and Axiom SNP array data in the outcrossing species *Malus* × *domestica* and causes for seemingly incompatible calls"

### Slide 1
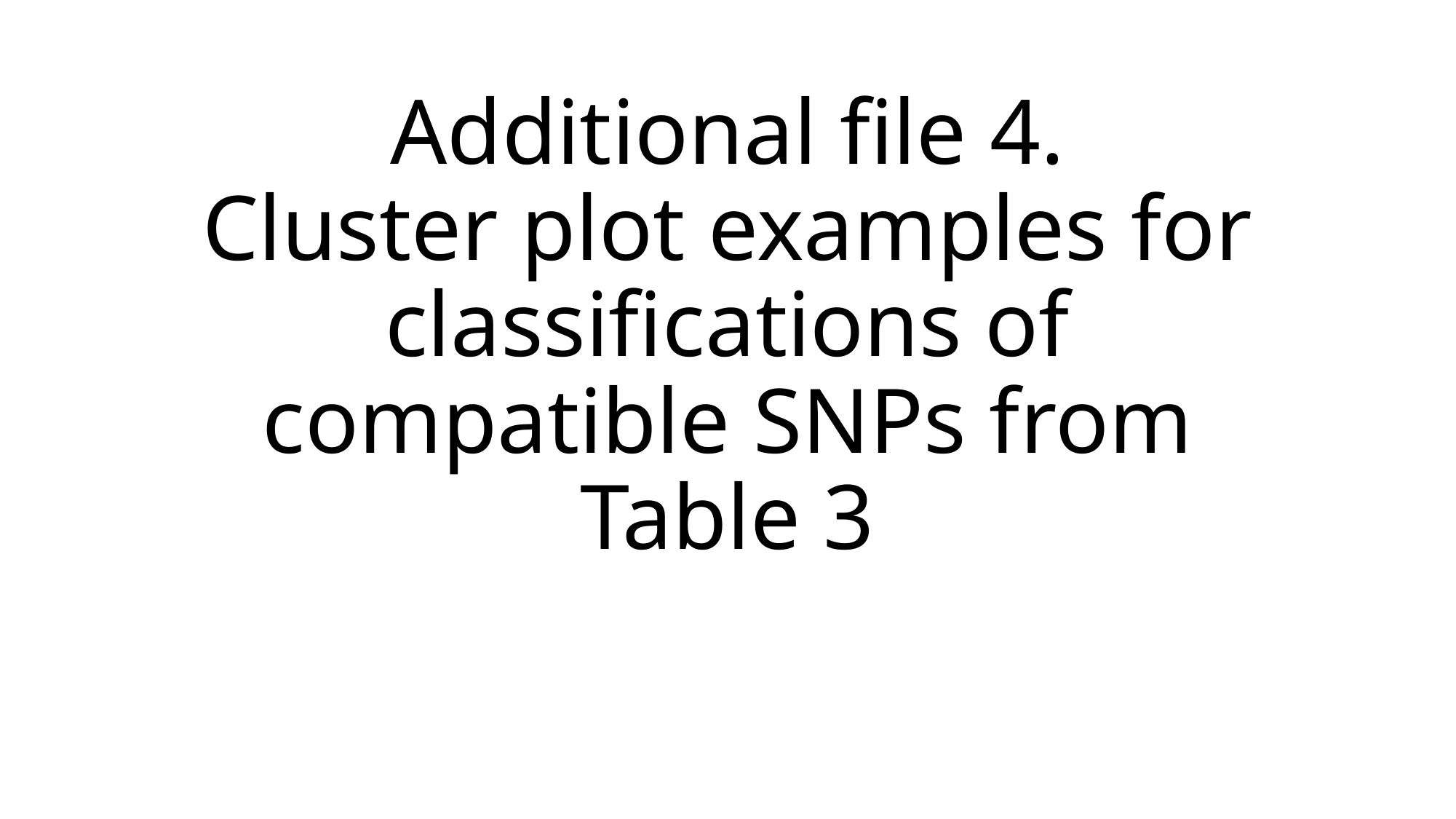

#### Additional file 4. Cluster plot examples for classifications of compatible SNPs from Table 3

### Slide 2
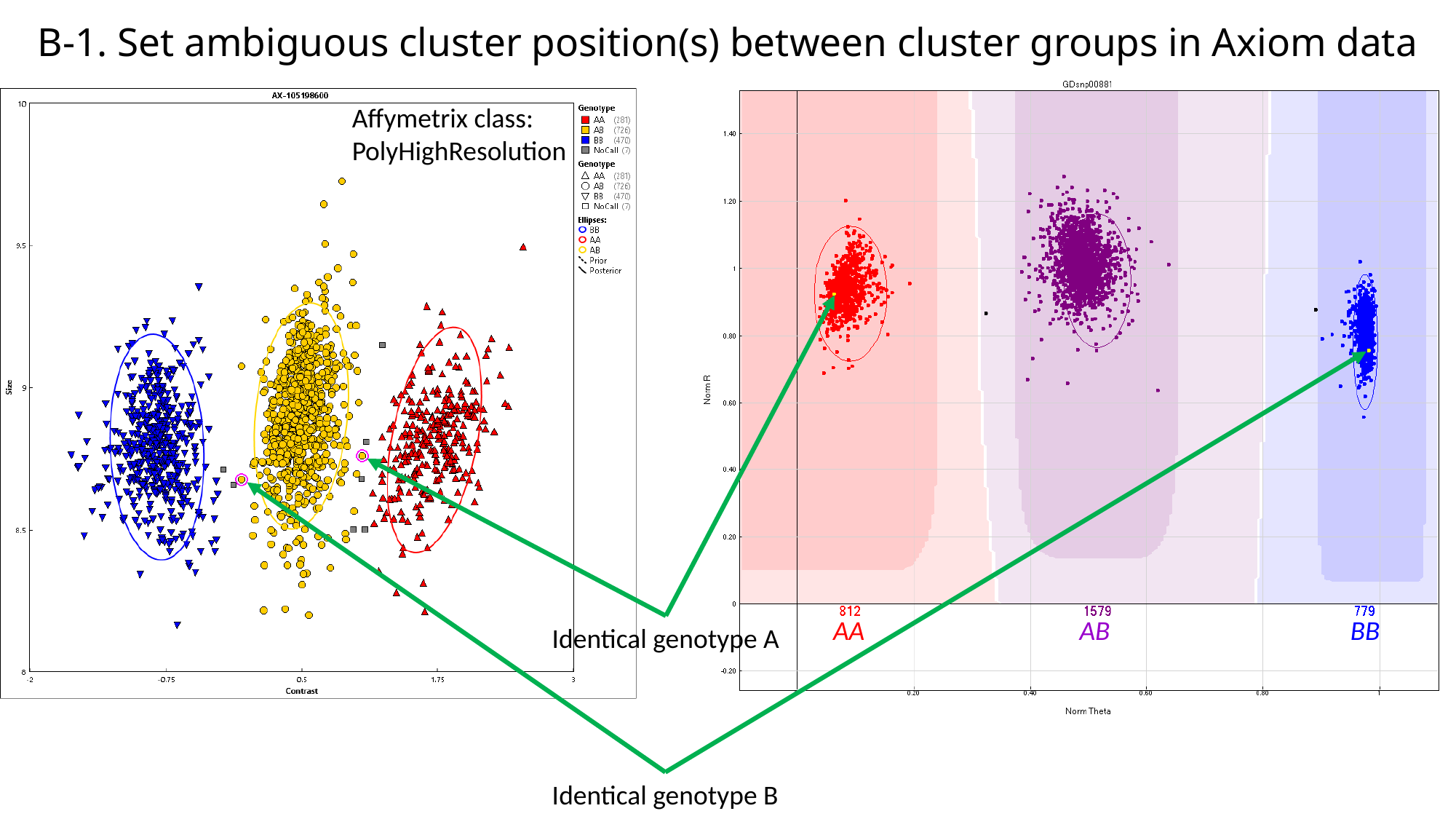

#### B-1. Set ambiguous cluster position(s) between cluster groups in Axiom data
Affymetrix class:
PolyHighResolution
AA
AB
BB
Identical genotype A
Identical genotype B

### Slide 3
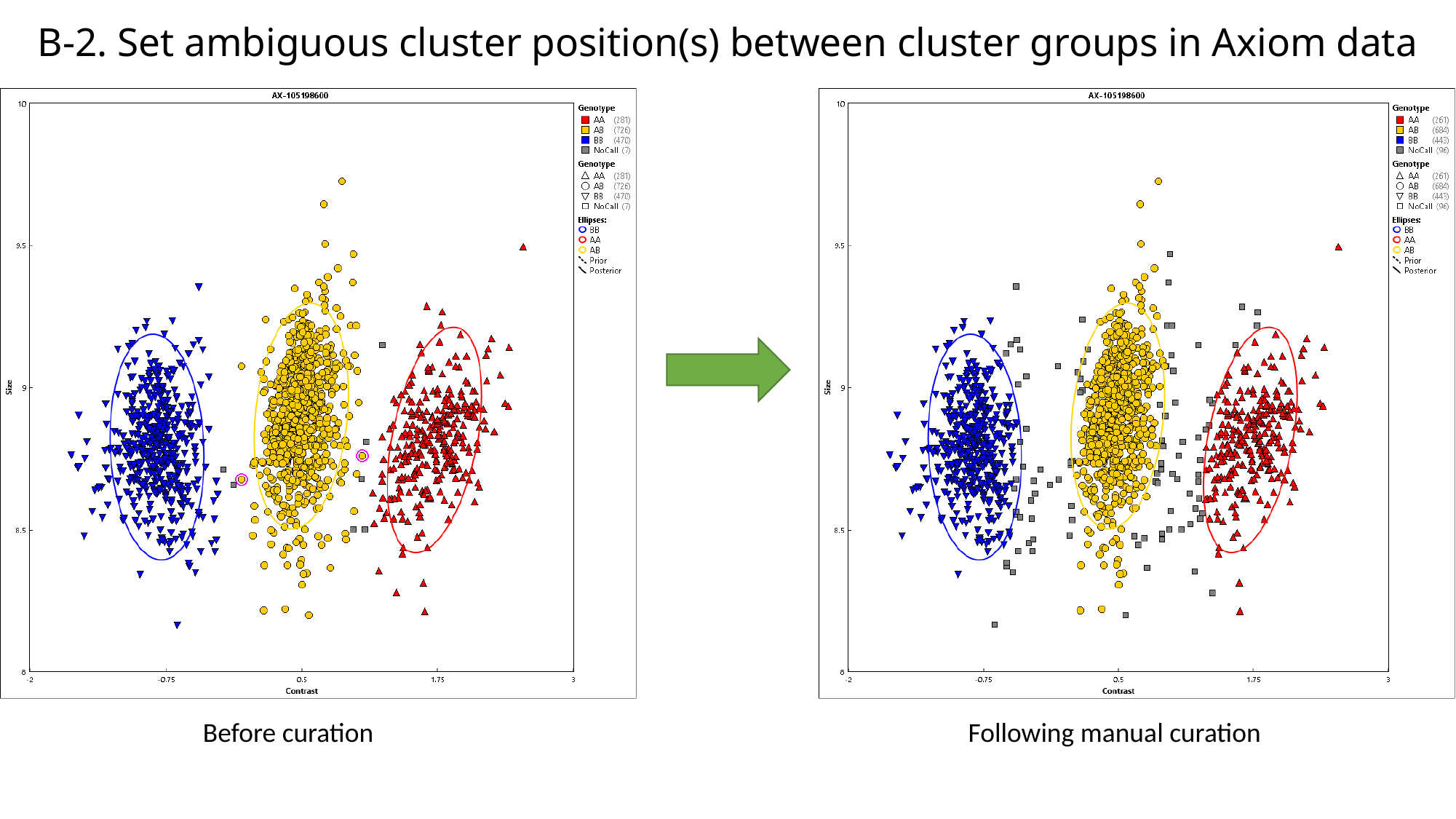

#### B-2. Set ambiguous cluster position(s) between cluster groups in Axiom data
Before curation
Following manual curation

### Slide 4
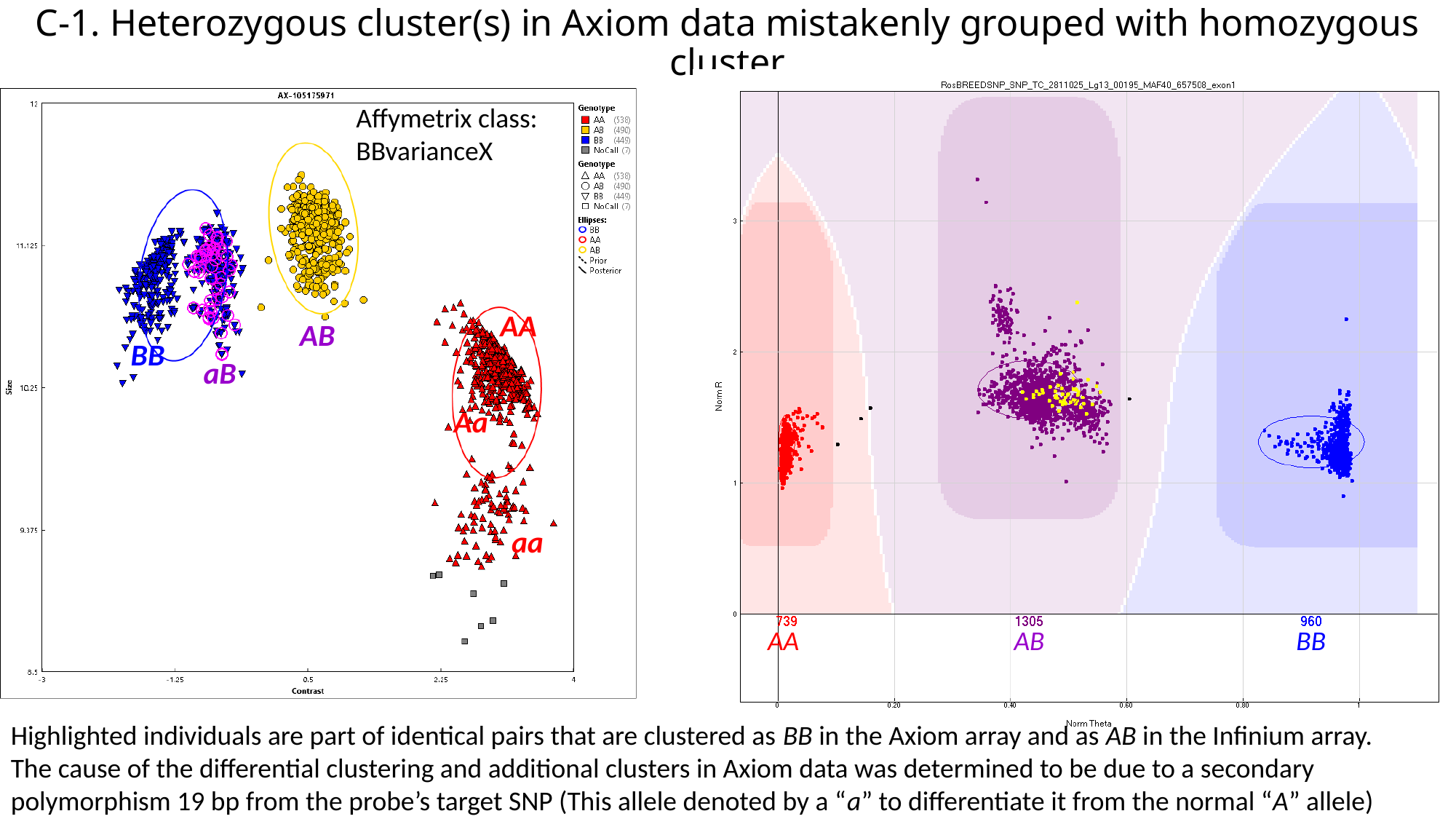

#### C-1. Heterozygous cluster(s) in Axiom data mistakenly grouped with homozygous cluster
Affymetrix class:
BBvarianceX
AA
AB
BB
aB
Aa
aa
AA
AB
BB
Highlighted individuals are part of identical pairs that are clustered as BB in the Axiom array and as AB in the Infinium array. The cause of the differential clustering and additional clusters in Axiom data was determined to be due to a secondary polymorphism 19 bp from the probe’s target SNP (This allele denoted by a “a” to differentiate it from the normal “A” allele)

### Slide 5
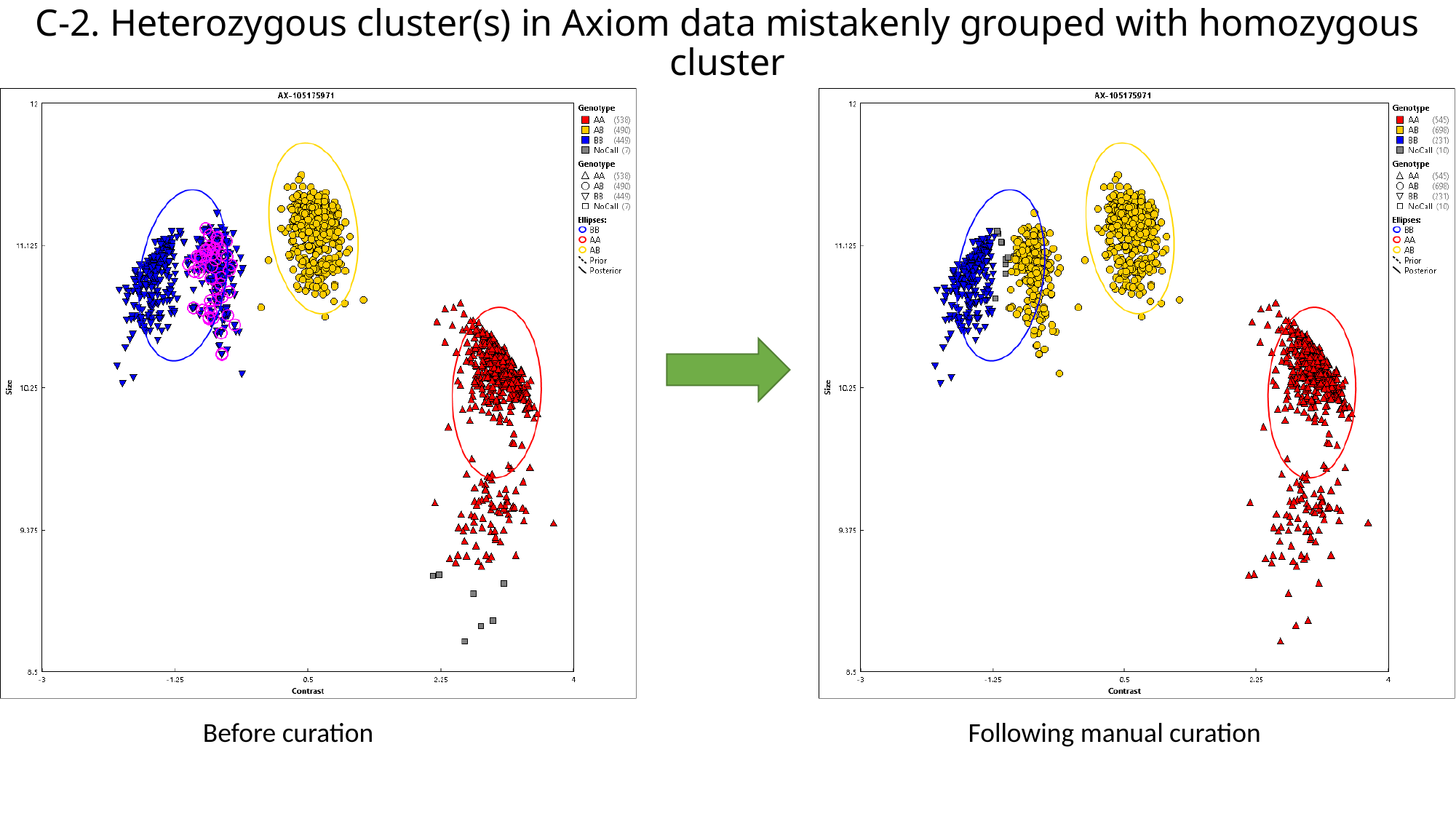

#### C-2. Heterozygous cluster(s) in Axiom data mistakenly grouped with homozygous cluster
Before curation
Following manual curation

### Slide 6
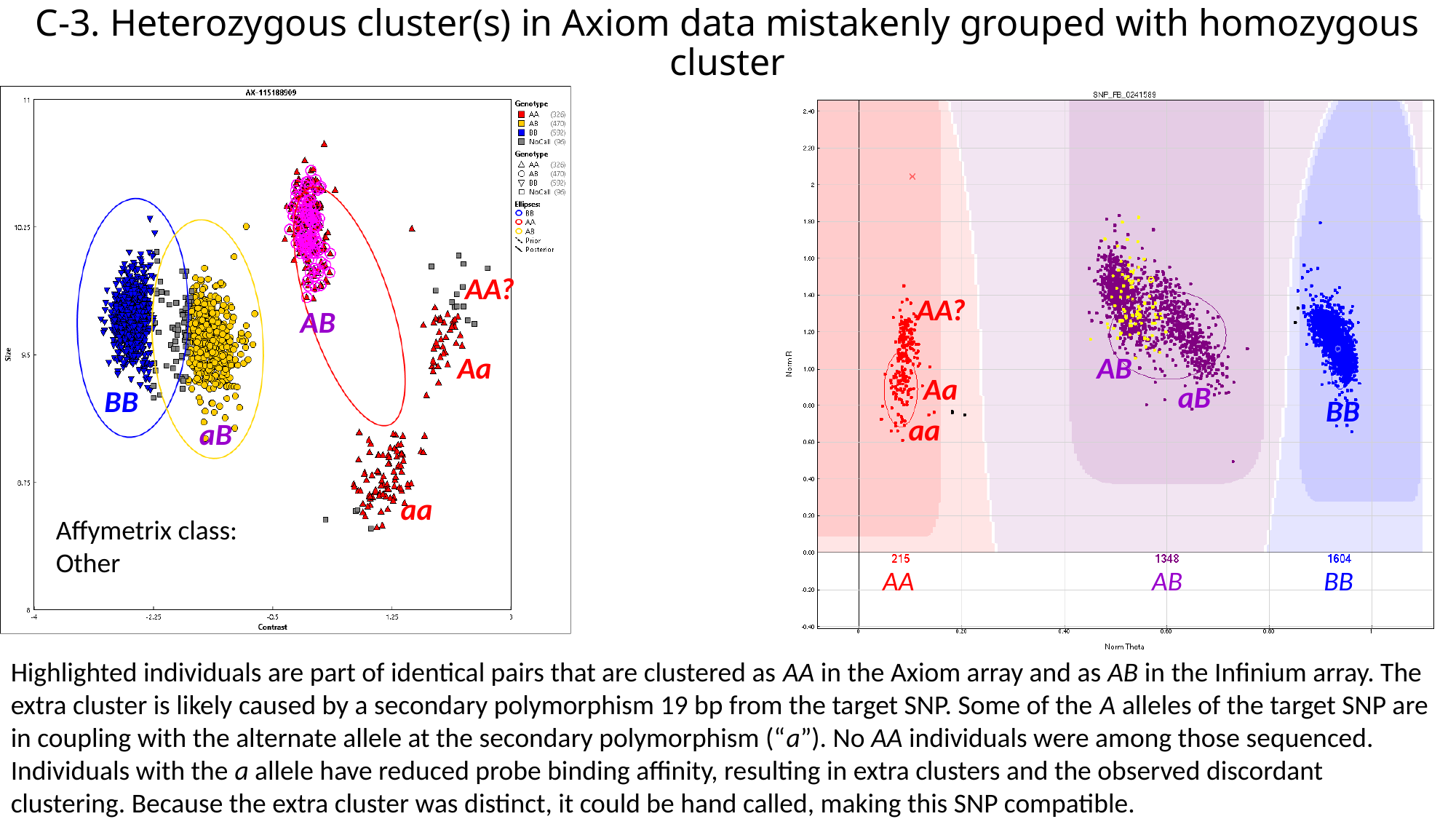

#### C-3. Heterozygous cluster(s) in Axiom data mistakenly grouped with homozygous cluster
AA?
AA?
AB
Aa
AB
Aa
aB
BB
BB
aa
aB
aa
Affymetrix class:
Other
AA
AB
BB
Highlighted individuals are part of identical pairs that are clustered as AA in the Axiom array and as AB in the Infinium array. The extra cluster is likely caused by a secondary polymorphism 19 bp from the target SNP. Some of the A alleles of the target SNP are in coupling with the alternate allele at the secondary polymorphism (“a”). No AA individuals were among those sequenced. Individuals with the a allele have reduced probe binding affinity, resulting in extra clusters and the observed discordant clustering. Because the extra cluster was distinct, it could be hand called, making this SNP compatible.

### Slide 7
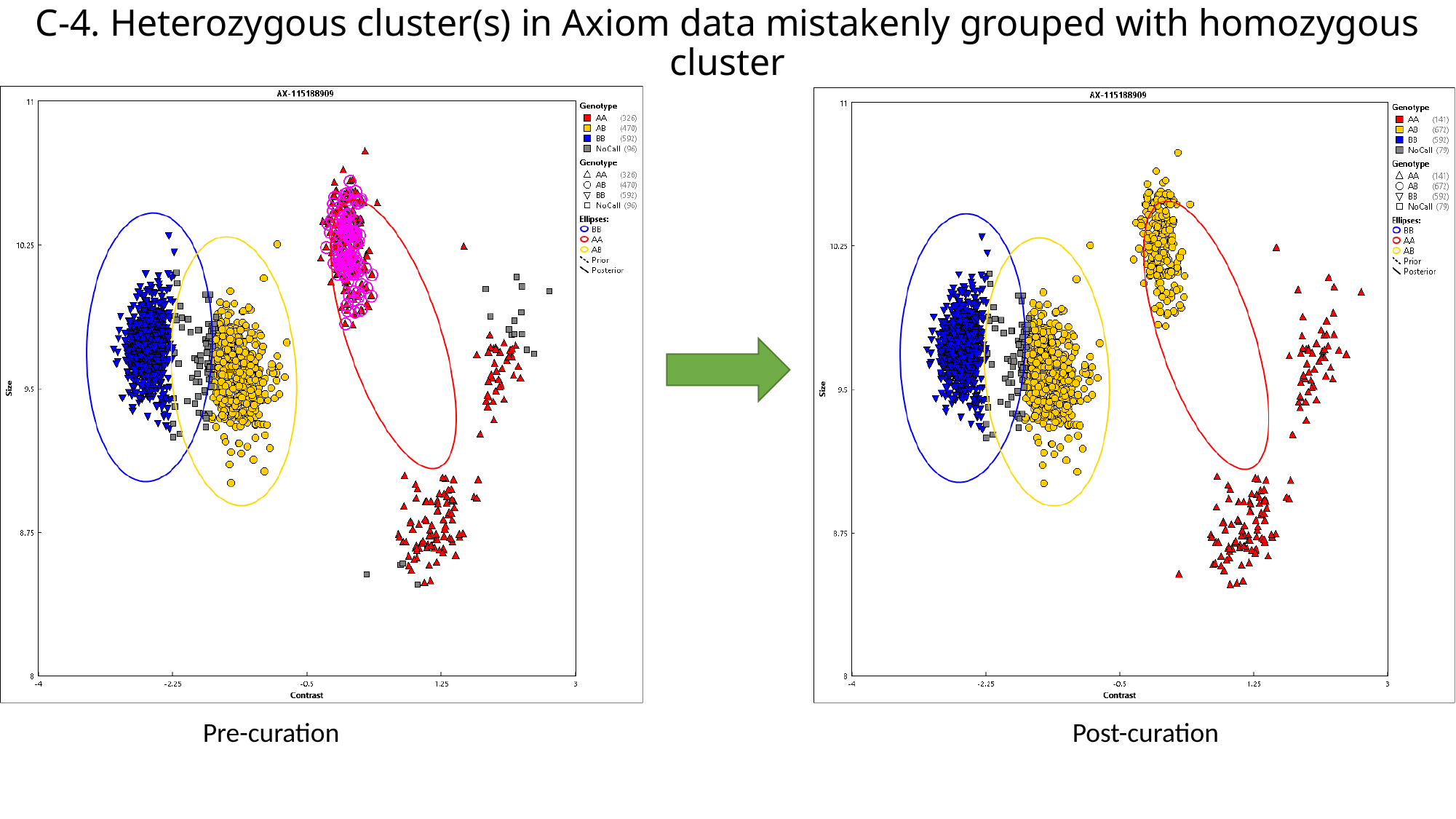

#### C-4. Heterozygous cluster(s) in Axiom data mistakenly grouped with homozygous cluster
Pre-curation
Post-curation

### Slide 8
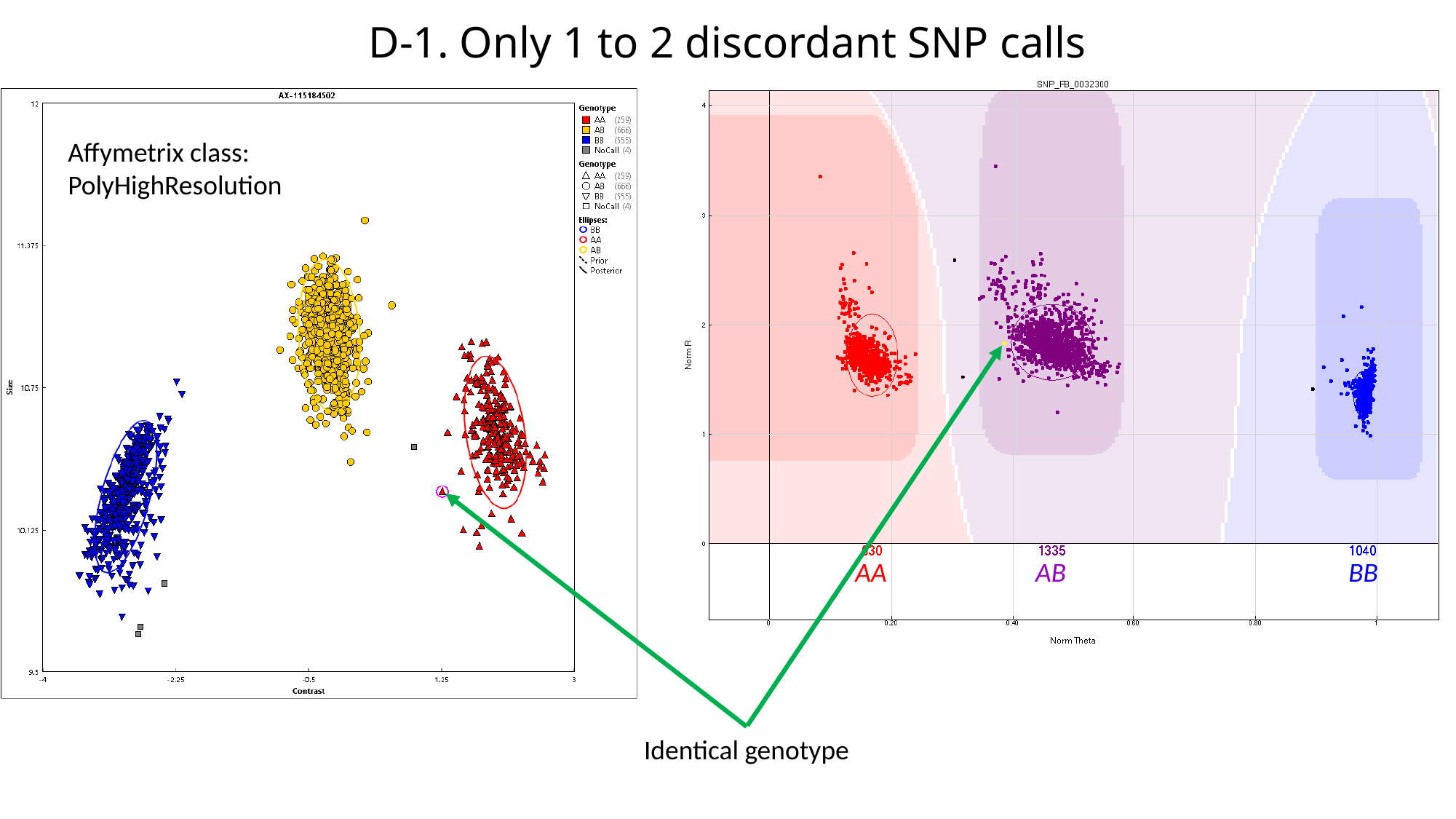

#### D-1. Only 1 to 2 discordant SNP calls
Affymetrix class:
PolyHighResolution
AA
AB
BB
Identical genotype

### Slide 9
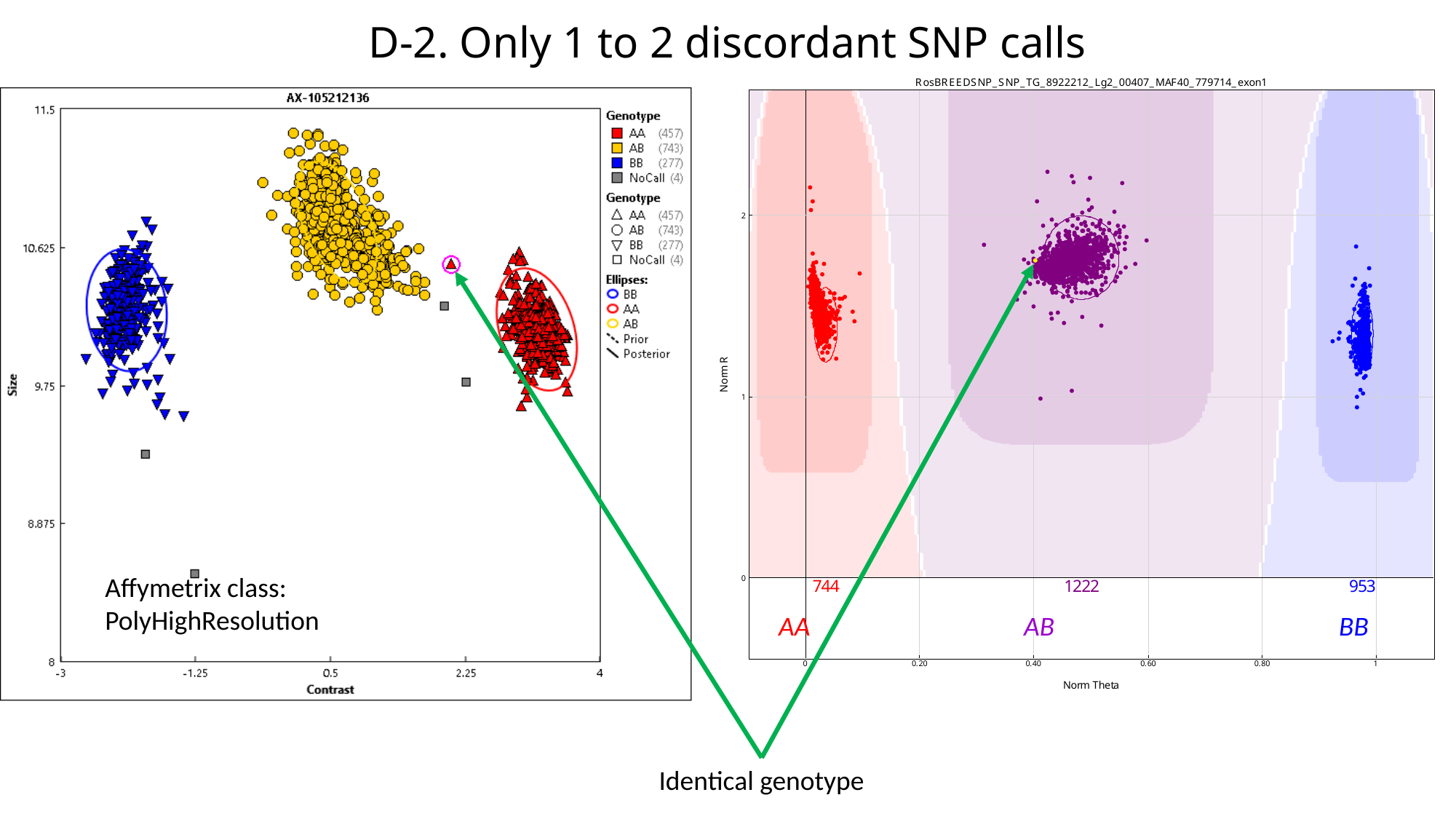

#### D-2. Only 1 to 2 discordant SNP calls
Affymetrix class:
PolyHighResolution
AA
AB
BB
Identical genotype

### Slide 10
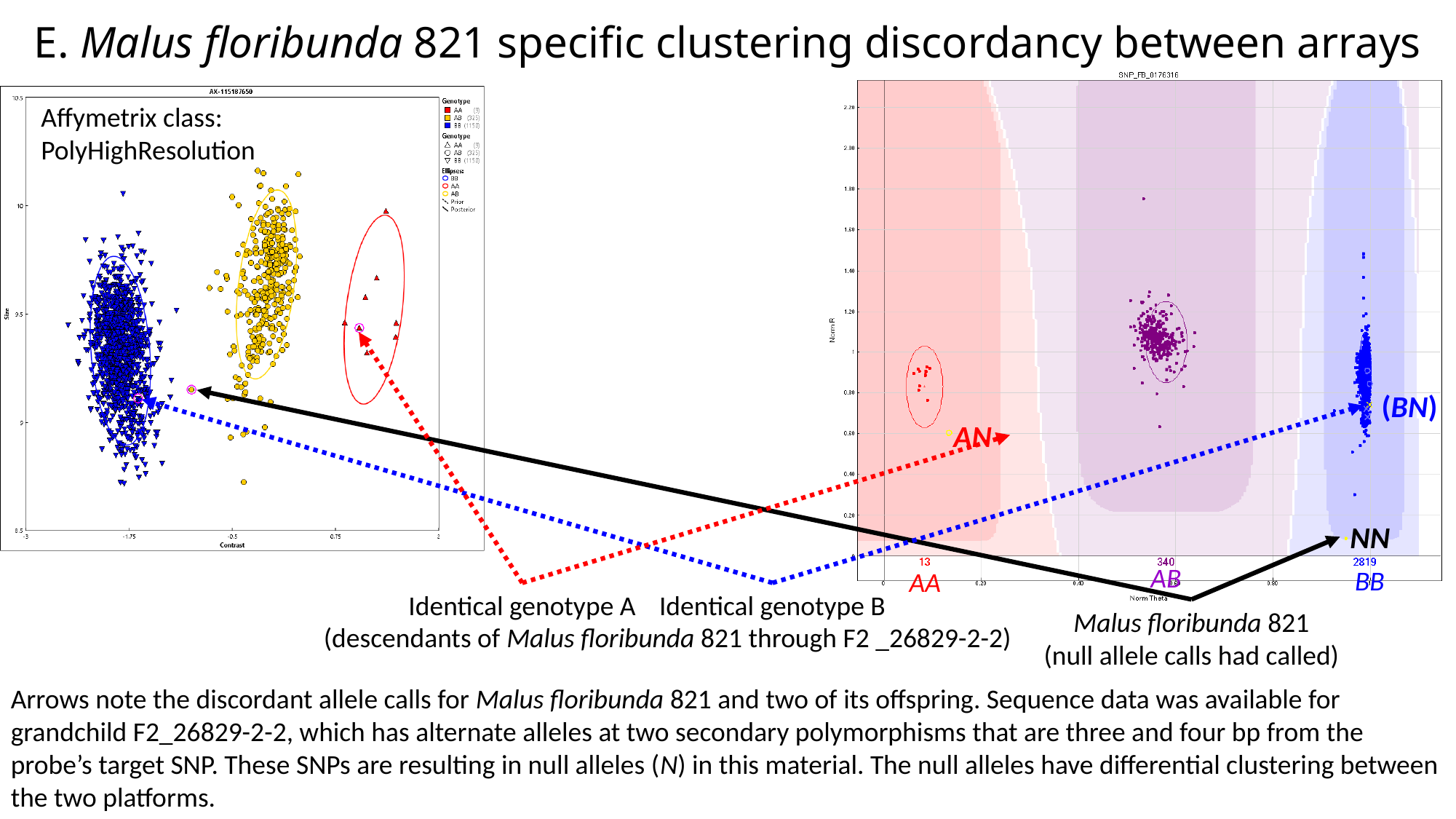

#### E. Malus floribunda 821 specific clustering discordancy between arrays
Affymetrix class:
PolyHighResolution
(BN)
AN
NN
AB
BB
AA
Identical genotype A
Identical genotype B
Malus floribunda 821
(null allele calls had called)
(descendants of Malus floribunda 821 through F2 _26829-2-2)
Arrows note the discordant allele calls for Malus floribunda 821 and two of its offspring. Sequence data was available for grandchild F2_26829-2-2, which has alternate alleles at two secondary polymorphisms that are three and four bp from the probe’s target SNP. These SNPs are resulting in null alleles (N) in this material. The null alleles have differential clustering between the two platforms.

### Slide 11
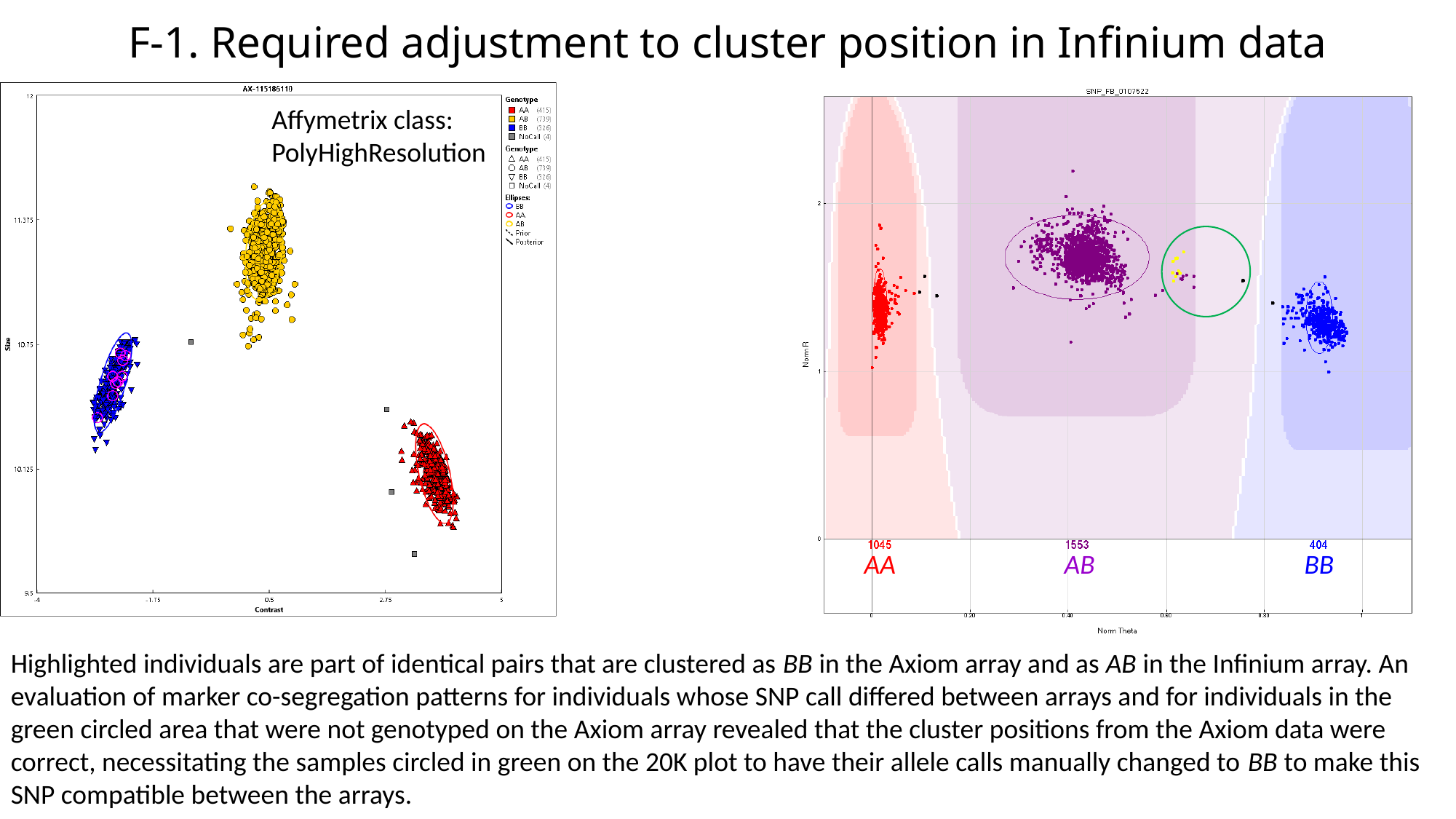

#### F-1. Required adjustment to cluster position in Infinium data
Affymetrix class:
PolyHighResolution
AA
AB
BB
Highlighted individuals are part of identical pairs that are clustered as BB in the Axiom array and as AB in the Infinium array. An evaluation of marker co-segregation patterns for individuals whose SNP call differed between arrays and for individuals in the green circled area that were not genotyped on the Axiom array revealed that the cluster positions from the Axiom data were correct, necessitating the samples circled in green on the 20K plot to have their allele calls manually changed to BB to make this SNP compatible between the arrays.

### Slide 12
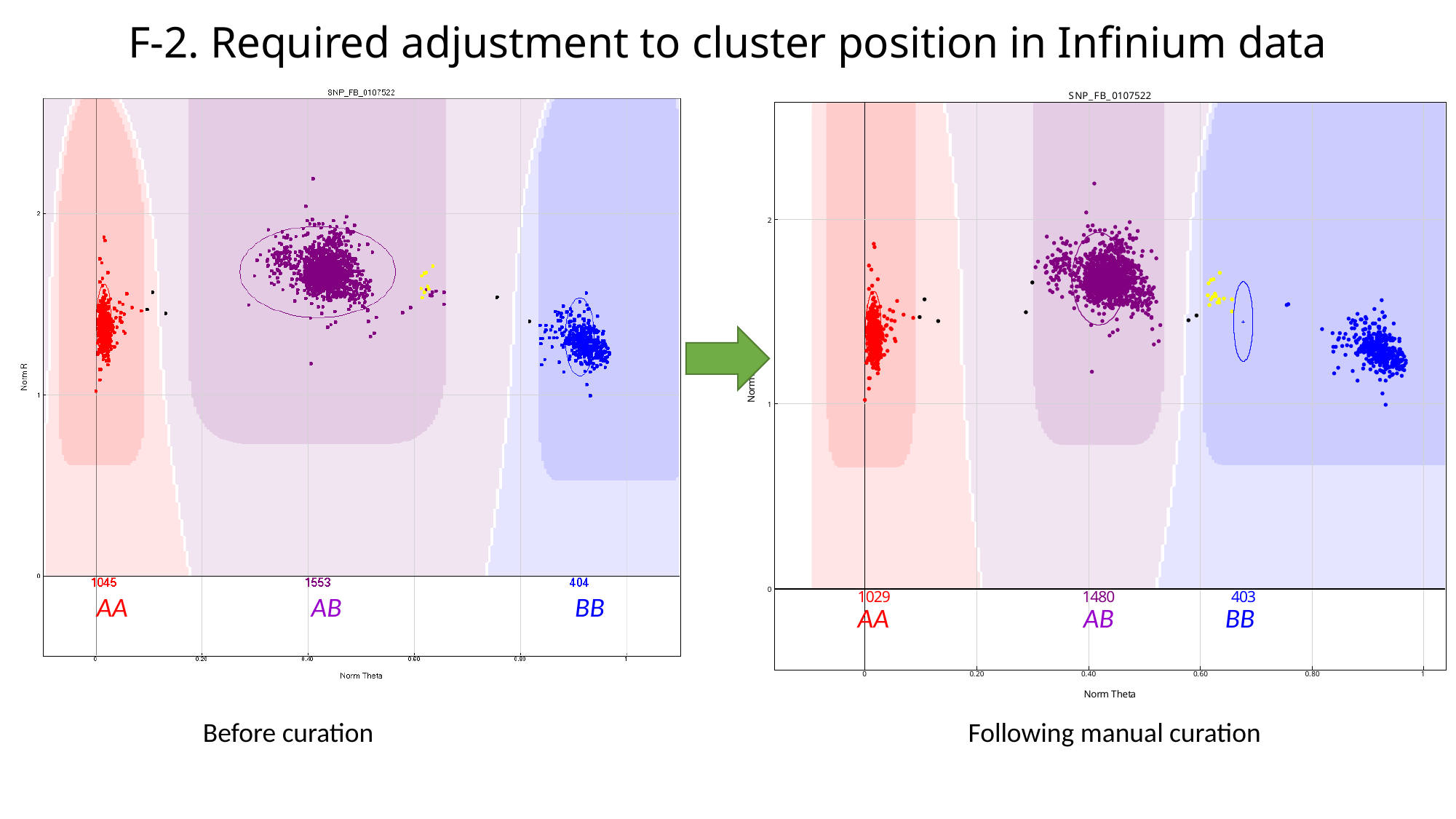

#### F-2. Required adjustment to cluster position in Infinium data
AA
AB
BB
AA
AB
BB
Before curation
Following manual curation

### Slide 13
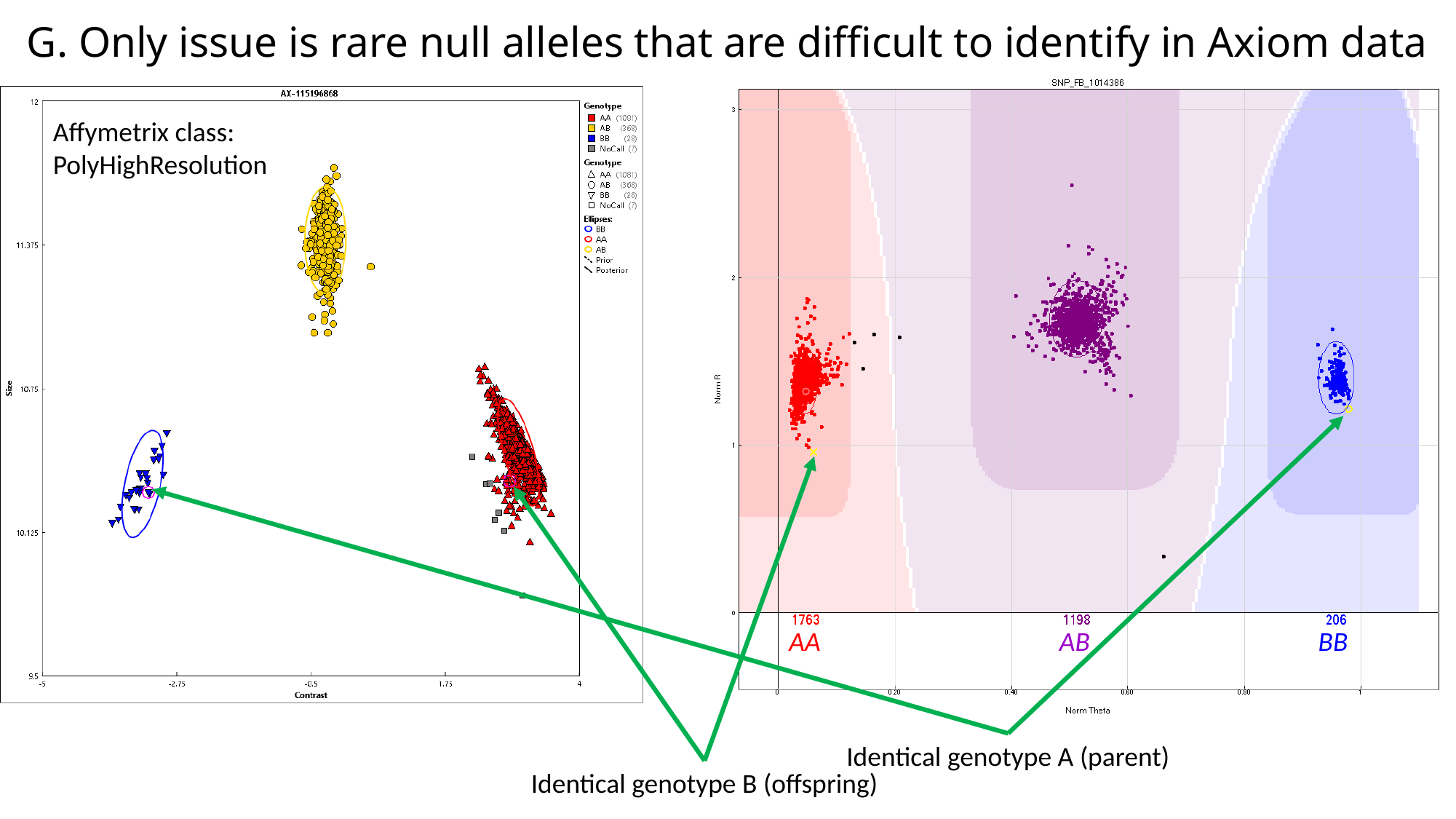

#### G. Only issue is rare null alleles that are difficult to identify in Axiom data
Affymetrix class:
PolyHighResolution
AA
AB
BB
Identical genotype A (parent)
Identical genotype B (offspring)

### Slide 14
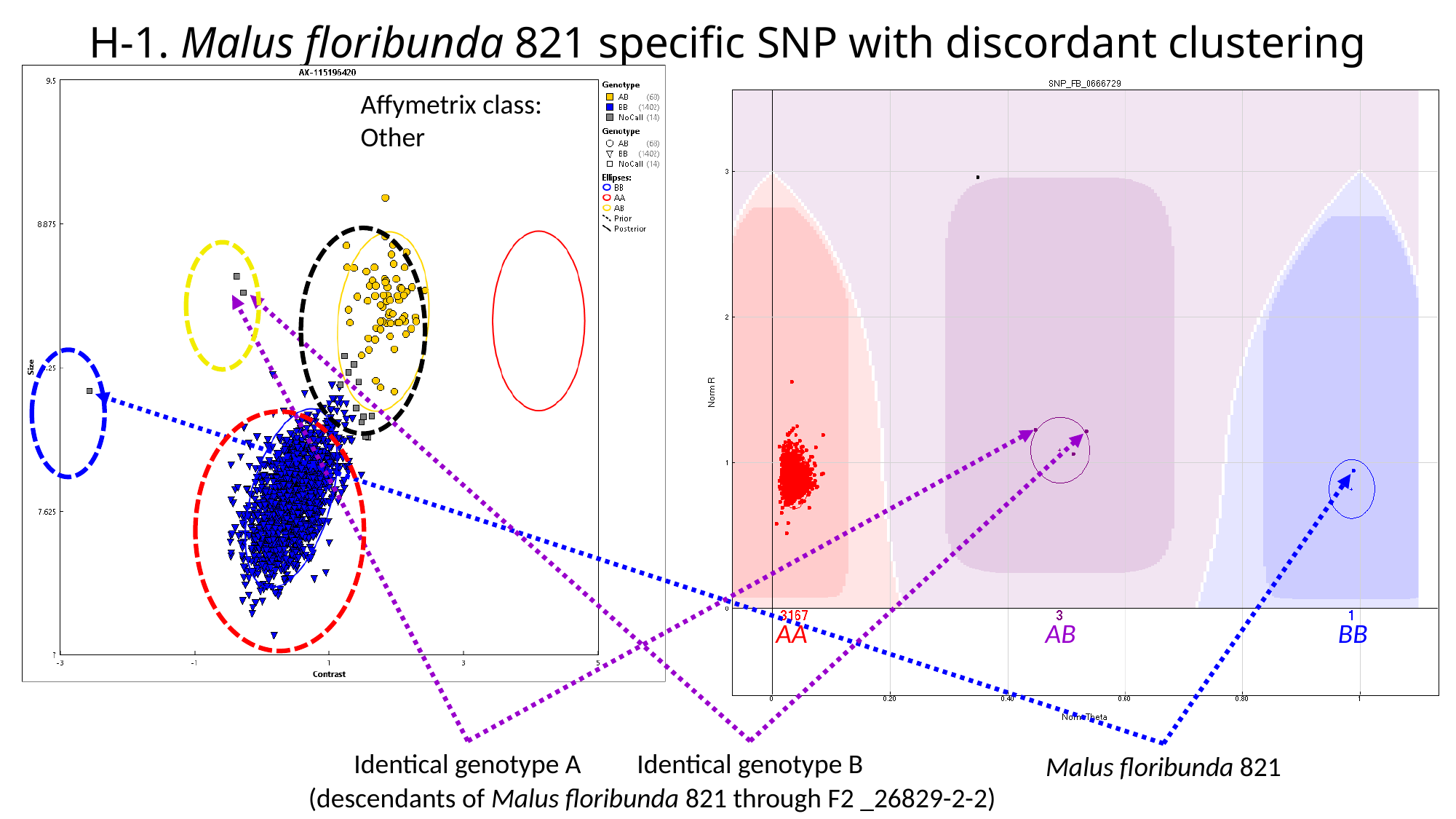

#### H-1. Malus floribunda 821 specific SNP with discordant clustering
Affymetrix class:
Other
I could also include an example of
assumed null alleles from 480K relationships
AA
AB
BB
Identical genotype A
Identical genotype B
Malus floribunda 821
(descendants of Malus floribunda 821 through F2 _26829-2-2)

### Slide 15
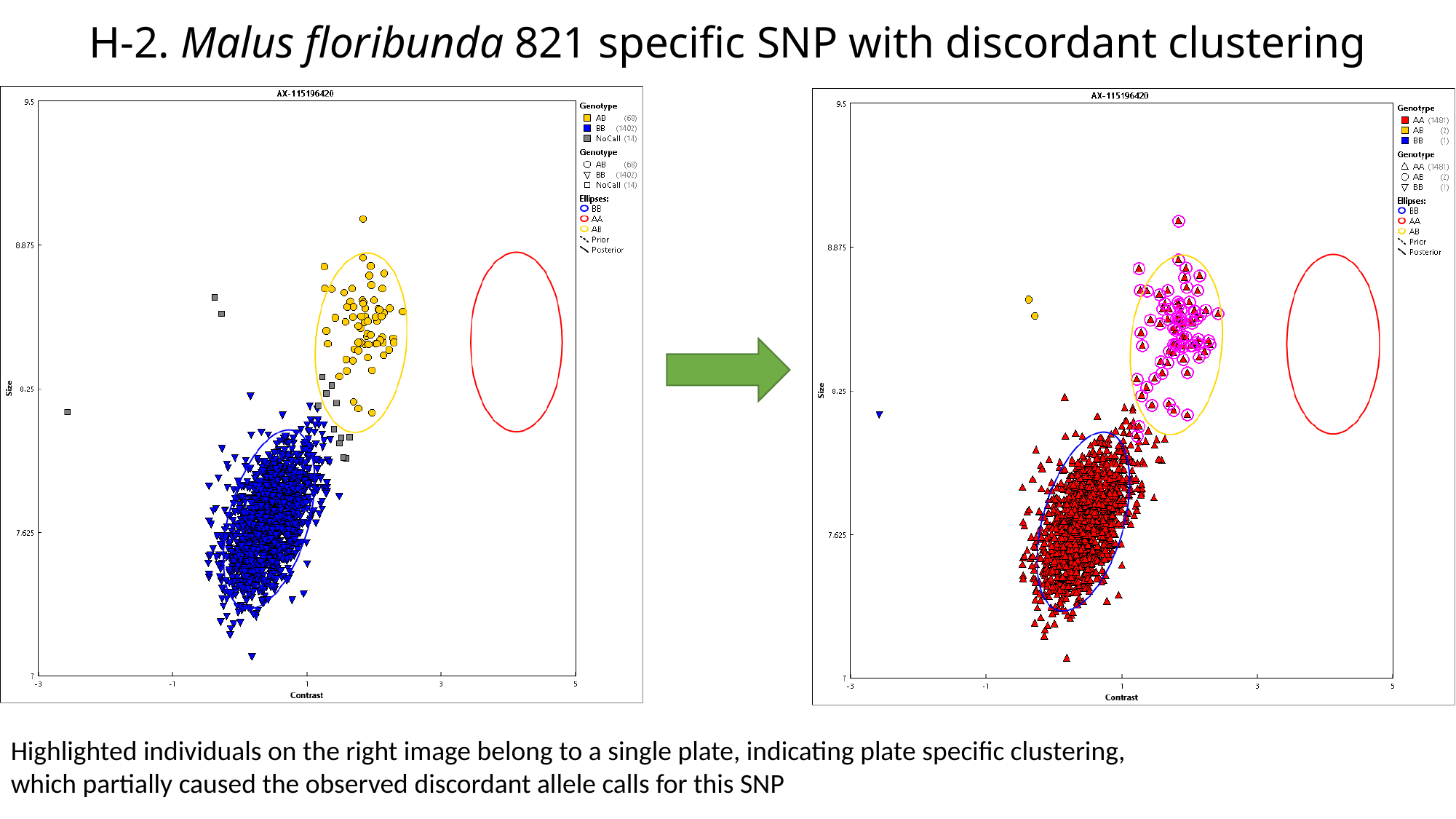

#### H-2. Malus floribunda 821 specific SNP with discordant clustering
Highlighted individuals on the right image belong to a single plate, indicating plate specific clustering,
which partially caused the observed discordant allele calls for this SNP

### Slide 16
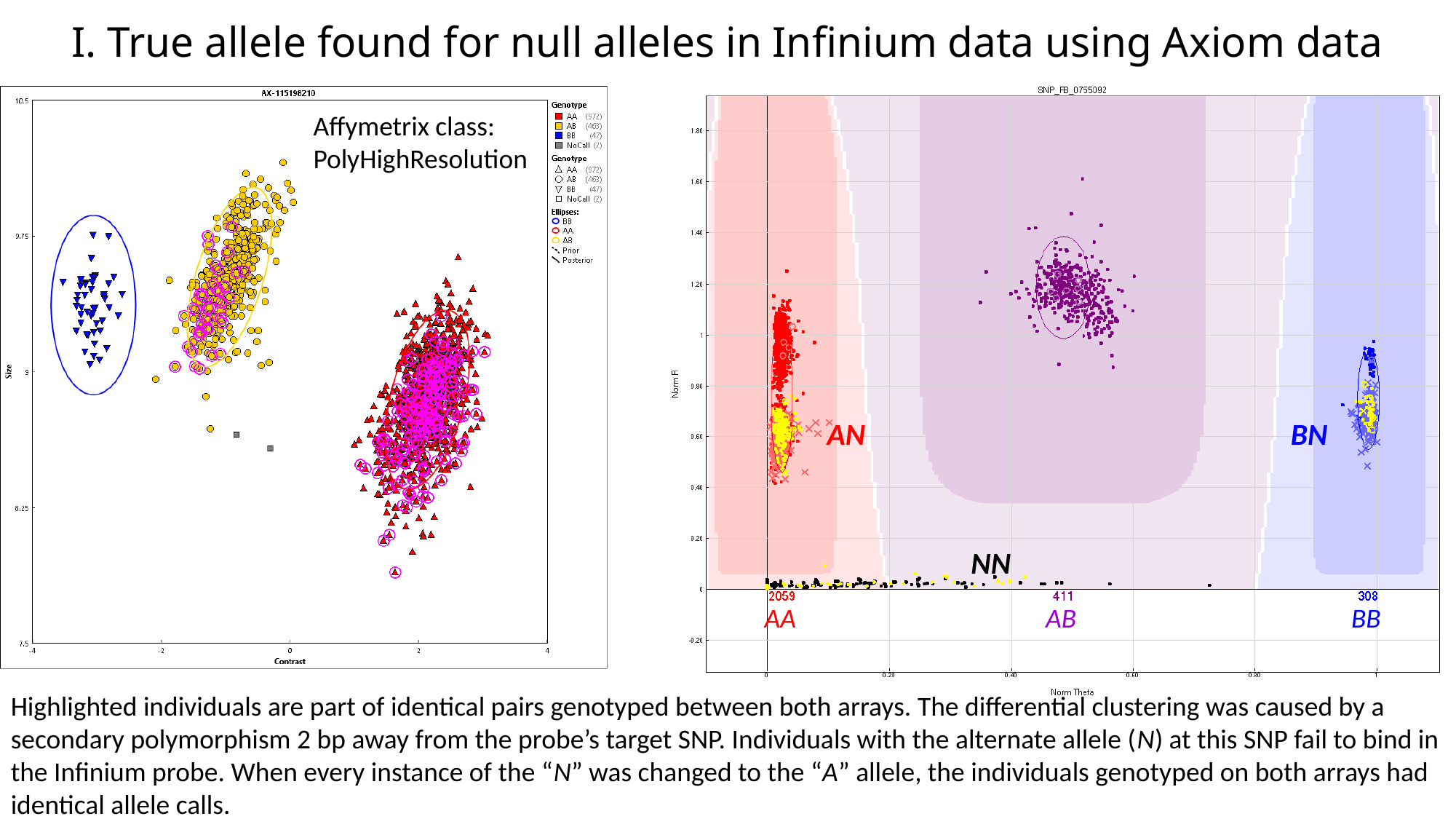

#### I. True allele found for null alleles in Infinium data using Axiom data
Affymetrix class:
PolyHighResolution
AN
BN
NN
AA
AB
BB
Highlighted individuals are part of identical pairs genotyped between both arrays. The differential clustering was caused by a secondary polymorphism 2 bp away from the probe’s target SNP. Individuals with the alternate allele (N) at this SNP fail to bind in the Infinium probe. When every instance of the “N” was changed to the “A” allele, the individuals genotyped on both arrays had identical allele calls.
