## Additional file 5 for "Integration of Infinium and Axiom SNP array data in the outcrossing species *Malus* × *domestica* and causes for seemingly incompatible calls"

### Slide 1
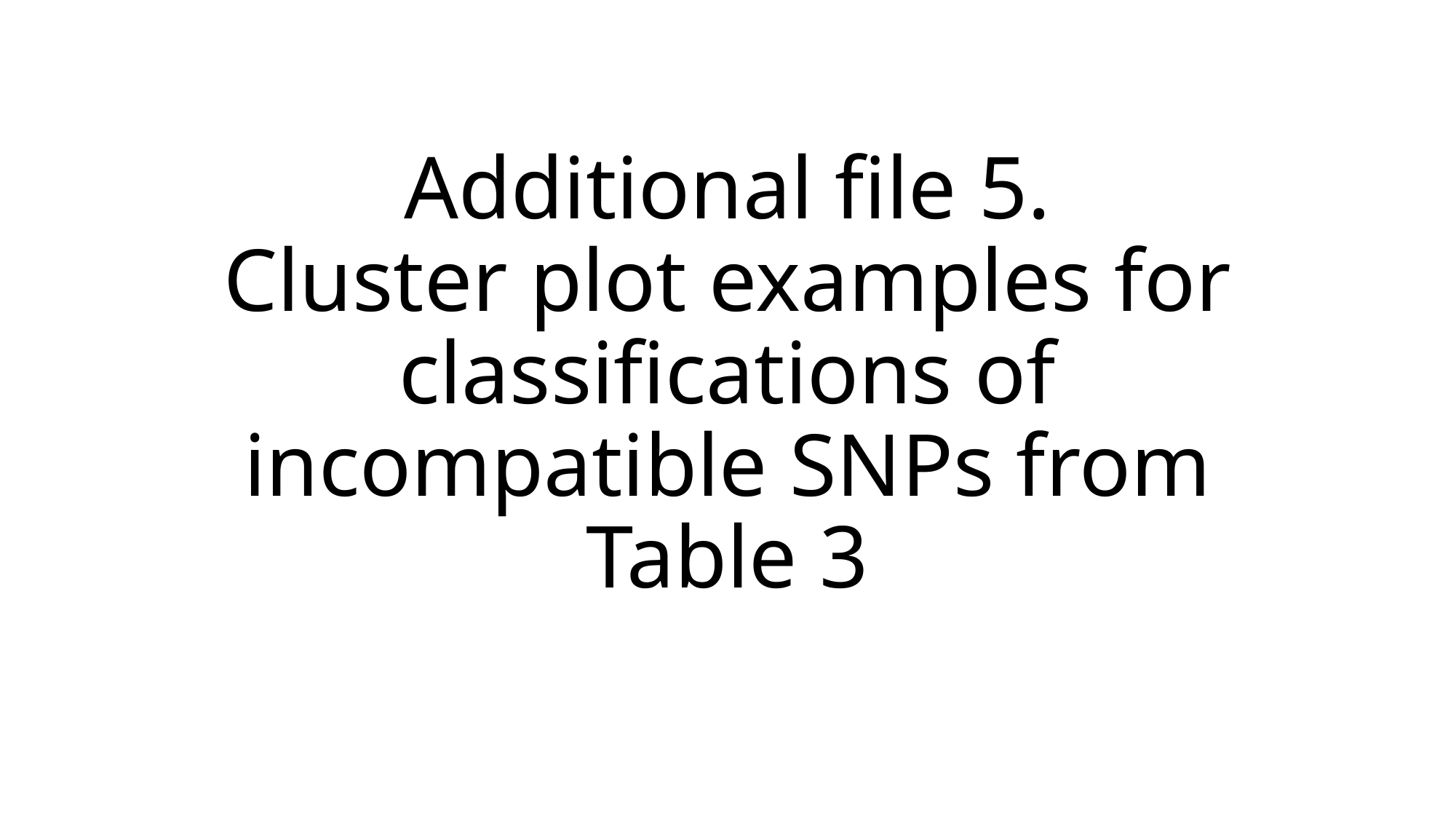

#### Additional file 5. Cluster plot examples for classifications of incompatible SNPs from Table 3

### Slide 2
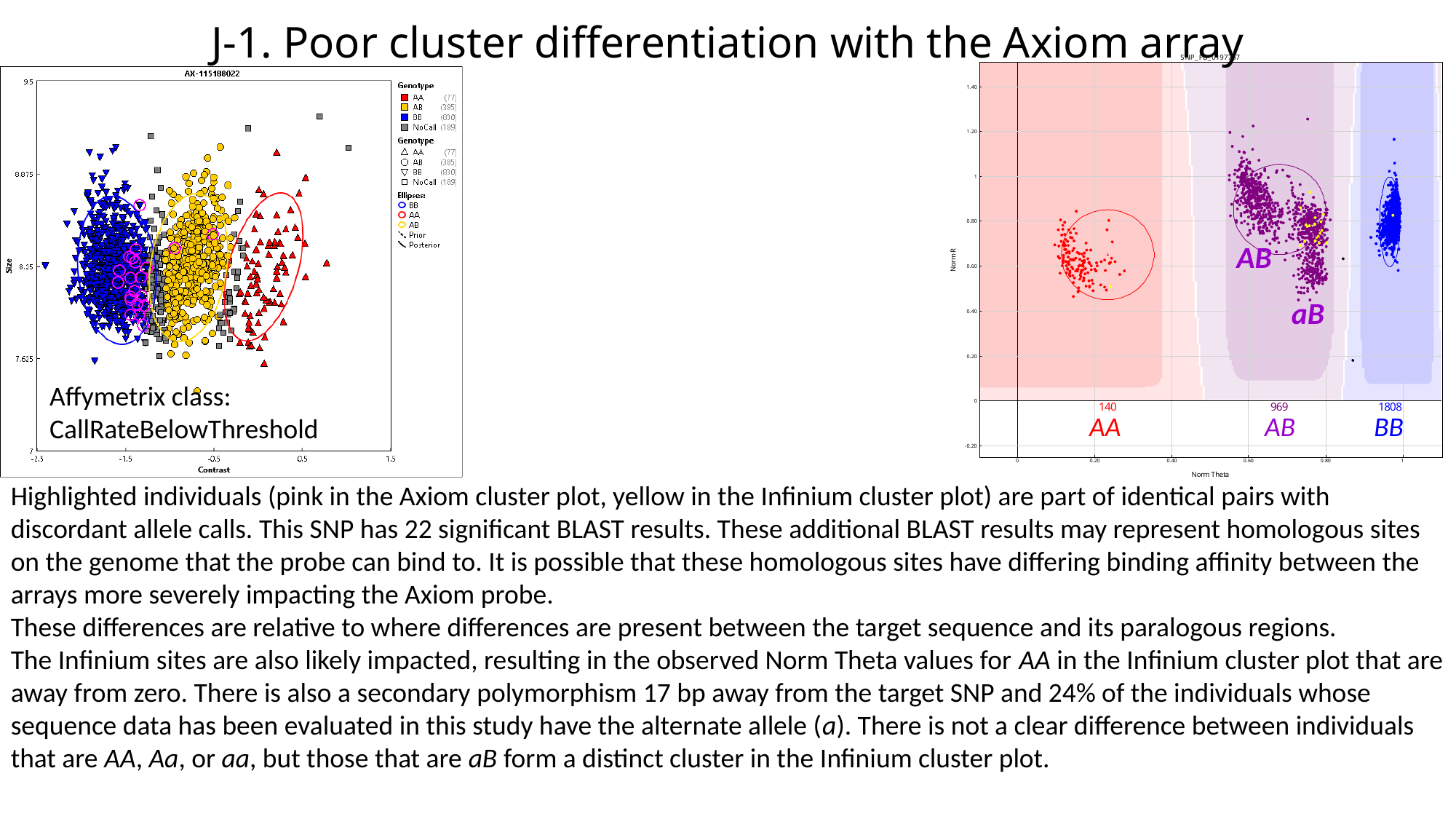

J-1. Poor cluster differentiation with the Axiom array
AB
aB
Affymetrix class:
CallRateBelowThreshold
AA
AB
BB
Highlighted individuals (pink in the Axiom cluster plot, yellow in the Infinium cluster plot) are part of identical pairs with discordant allele calls. This SNP has 22 significant BLAST results. These additional BLAST results may represent homologous sites on the genome that the probe can bind to. It is possible that these homologous sites have differing binding affinity between the arrays more severely impacting the Axiom probe.
These differences are relative to where differences are present between the target sequence and its paralogous regions.
The Infinium sites are also likely impacted, resulting in the observed Norm Theta values for AA in the Infinium cluster plot that are away from zero. There is also a secondary polymorphism 17 bp away from the target SNP and 24% of the individuals whose sequence data has been evaluated in this study have the alternate allele (a). There is not a clear difference between individuals that are AA, Aa, or aa, but those that are aB form a distinct cluster in the Infinium cluster plot.

### Slide 3
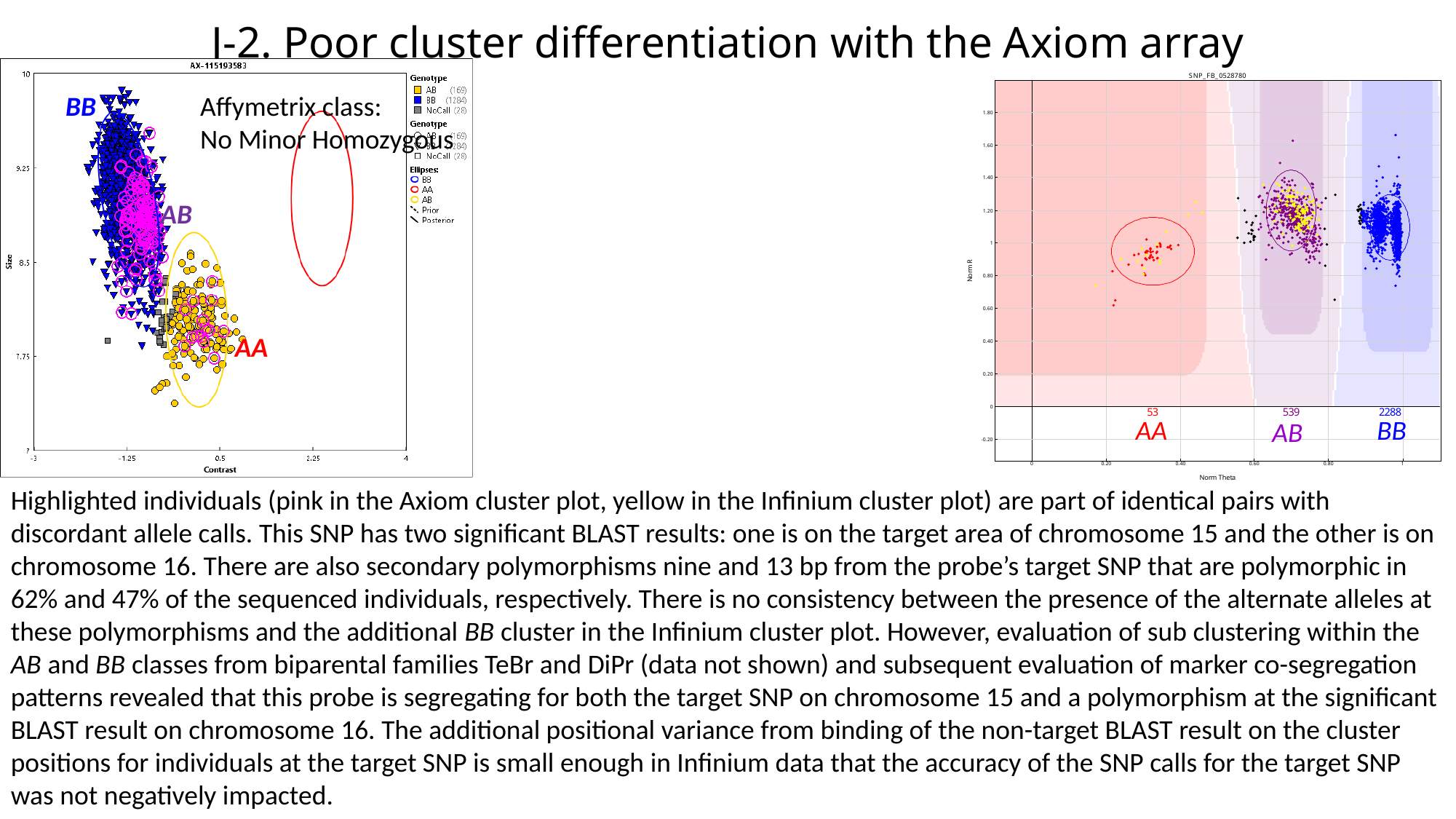

J-2. Poor cluster differentiation with the Axiom array
BB
Affymetrix class:
No Minor Homozygous
AB
AA
AA
BB
AB
Highlighted individuals (pink in the Axiom cluster plot, yellow in the Infinium cluster plot) are part of identical pairs with discordant allele calls. This SNP has two significant BLAST results: one is on the target area of chromosome 15 and the other is on chromosome 16. There are also secondary polymorphisms nine and 13 bp from the probe’s target SNP that are polymorphic in 62% and 47% of the sequenced individuals, respectively. There is no consistency between the presence of the alternate alleles at these polymorphisms and the additional BB cluster in the Infinium cluster plot. However, evaluation of sub clustering within the AB and BB classes from biparental families TeBr and DiPr (data not shown) and subsequent evaluation of marker co-segregation patterns revealed that this probe is segregating for both the target SNP on chromosome 15 and a polymorphism at the significant BLAST result on chromosome 16. The additional positional variance from binding of the non-target BLAST result on the cluster positions for individuals at the target SNP is small enough in Infinium data that the accuracy of the SNP calls for the target SNP was not negatively impacted.

### Slide 4
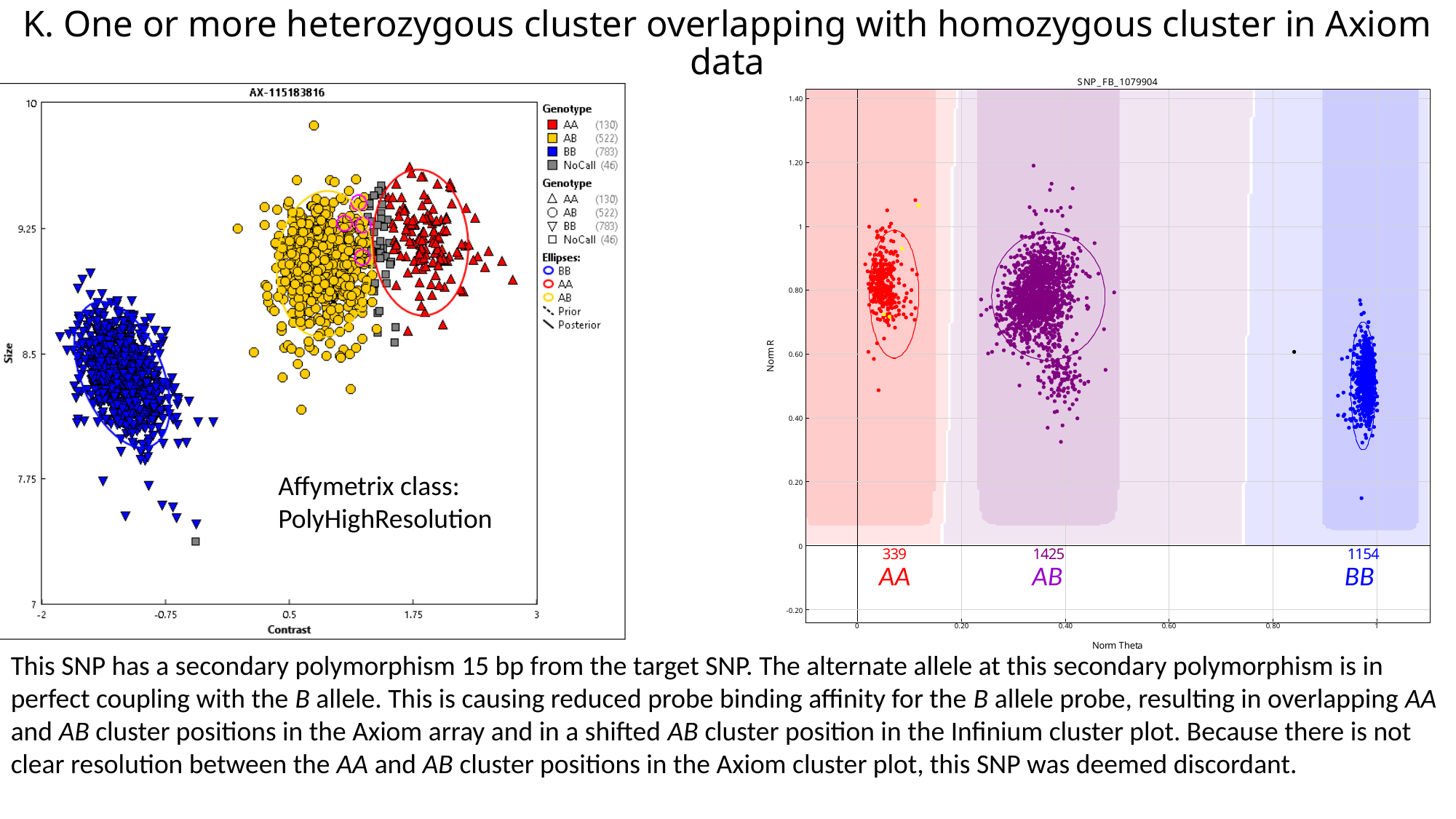

K. One or more heterozygous cluster overlapping with homozygous cluster in Axiom data
Affymetrix class:
PolyHighResolution
AA
AB
BB
This SNP has a secondary polymorphism 15 bp from the target SNP. The alternate allele at this secondary polymorphism is in perfect coupling with the B allele. This is causing reduced probe binding affinity for the B allele probe, resulting in overlapping AA and AB cluster positions in the Axiom array and in a shifted AB cluster position in the Infinium cluster plot. Because there is not clear resolution between the AA and AB cluster positions in the Axiom cluster plot, this SNP was deemed discordant.

### Slide 5
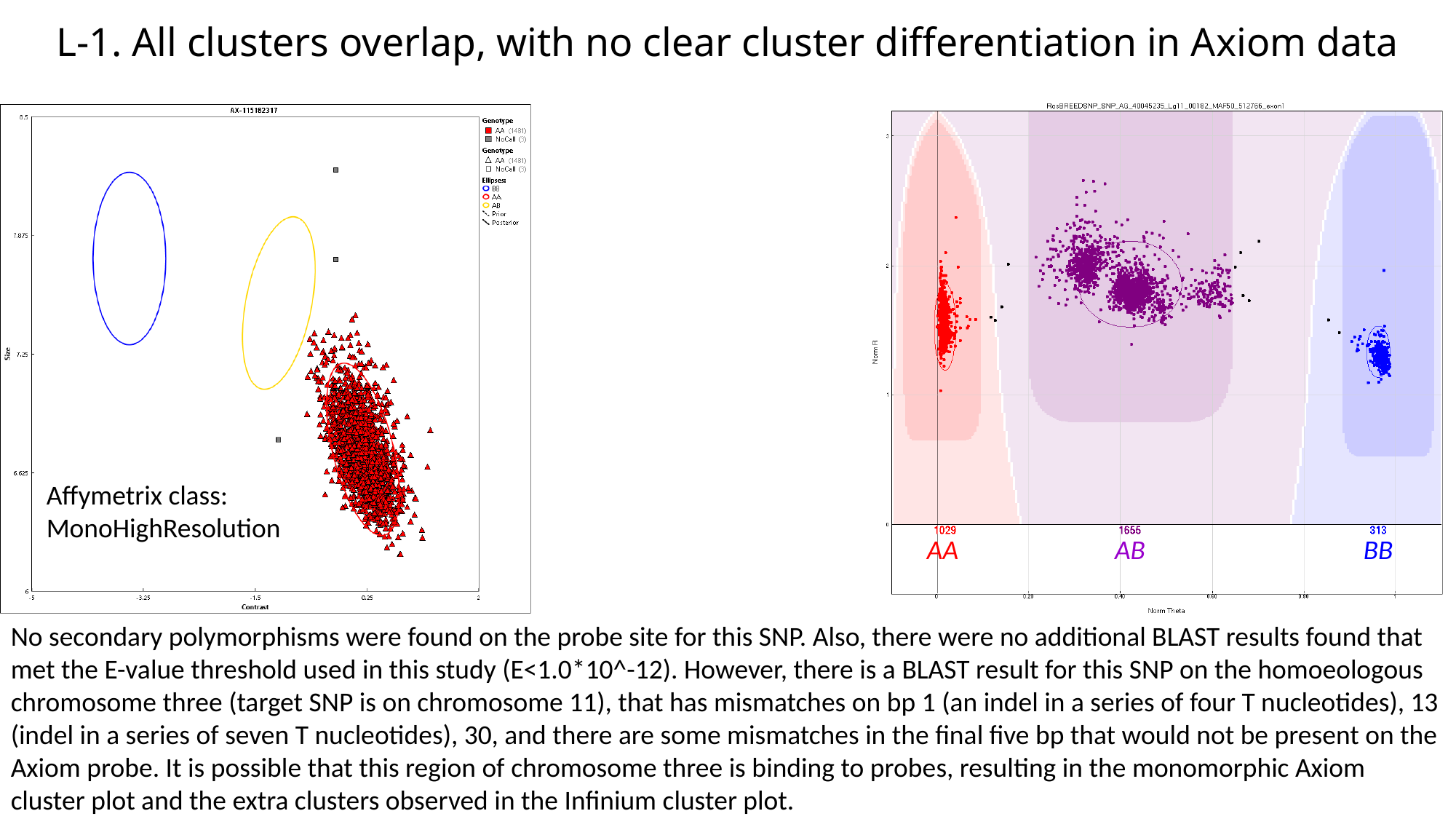

L-1. All clusters overlap, with no clear cluster differentiation in Axiom data
Affymetrix class:
MonoHighResolution
AA
AB
BB
No secondary polymorphisms were found on the probe site for this SNP. Also, there were no additional BLAST results found that met the E-value threshold used in this study (E<1.0*10^-12). However, there is a BLAST result for this SNP on the homoeologous chromosome three (target SNP is on chromosome 11), that has mismatches on bp 1 (an indel in a series of four T nucleotides), 13 (indel in a series of seven T nucleotides), 30, and there are some mismatches in the final five bp that would not be present on the Axiom probe. It is possible that this region of chromosome three is binding to probes, resulting in the monomorphic Axiom cluster plot and the extra clusters observed in the Infinium cluster plot.

### Slide 6
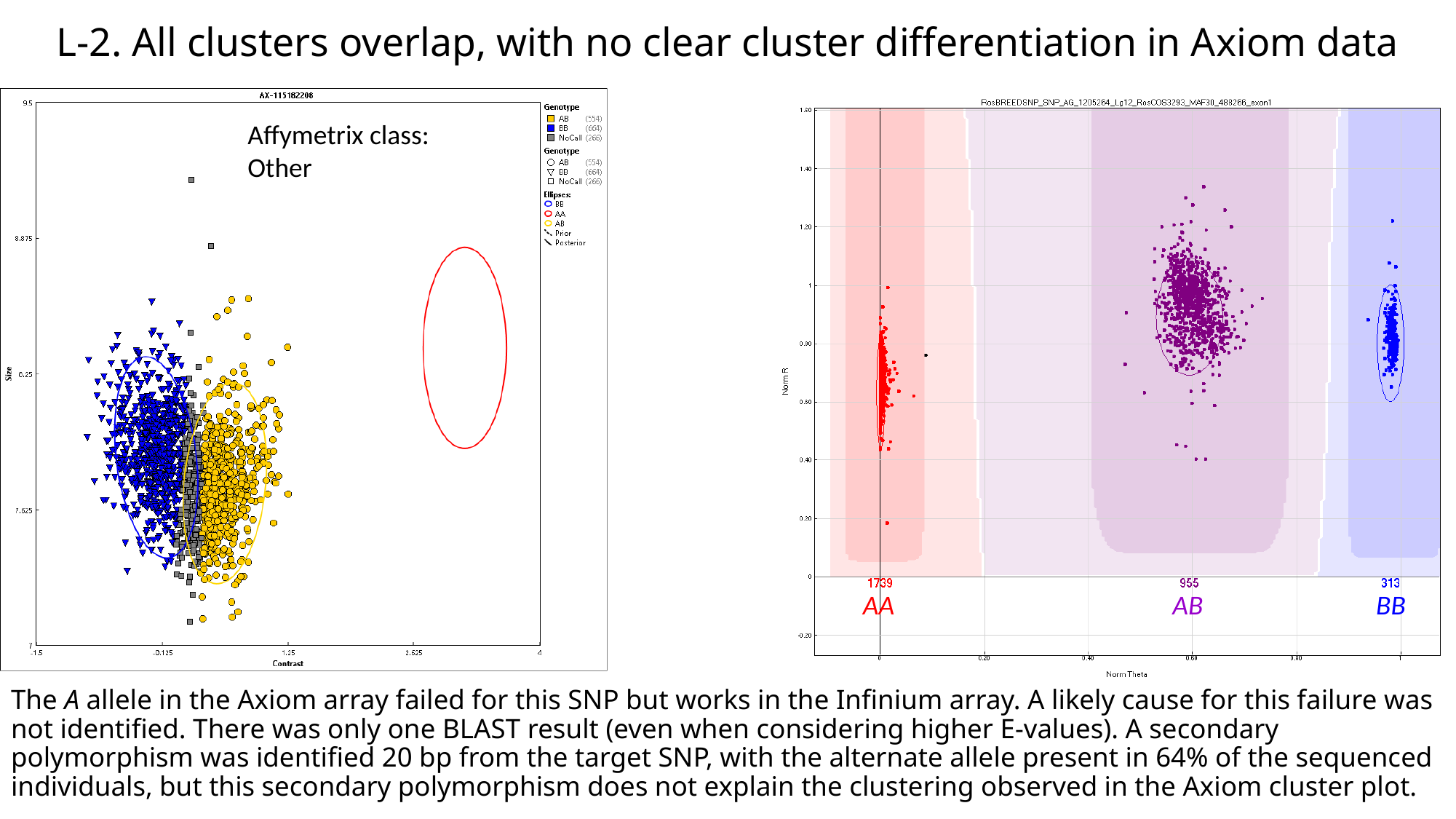

L-2. All clusters overlap, with no clear cluster differentiation in Axiom data
Affymetrix class:
Other
AA
AB
BB
The A allele in the Axiom array failed for this SNP but works in the Infinium array. A likely cause for this failure was not identified. There was only one BLAST result (even when considering higher E-values). A secondary polymorphism was identified 20 bp from the target SNP, with the alternate allele present in 64% of the sequenced individuals, but this secondary polymorphism does not explain the clustering observed in the Axiom cluster plot.

### Slide 7
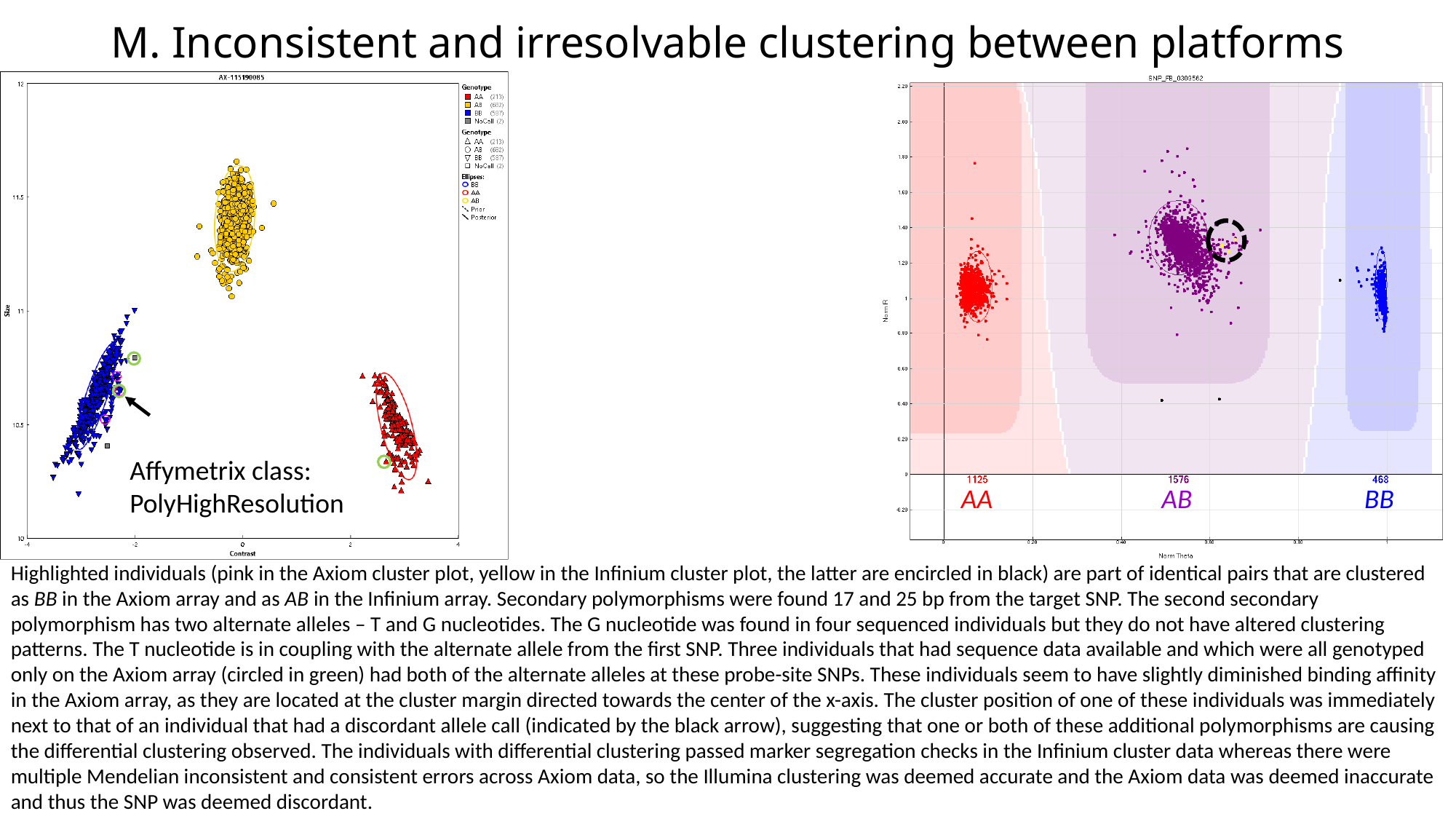

M. Inconsistent and irresolvable clustering between platforms
Affymetrix class:
PolyHighResolution
AA
AB
BB
Highlighted individuals (pink in the Axiom cluster plot, yellow in the Infinium cluster plot, the latter are encircled in black) are part of identical pairs that are clustered as BB in the Axiom array and as AB in the Infinium array. Secondary polymorphisms were found 17 and 25 bp from the target SNP. The second secondary polymorphism has two alternate alleles – T and G nucleotides. The G nucleotide was found in four sequenced individuals but they do not have altered clustering patterns. The T nucleotide is in coupling with the alternate allele from the first SNP. Three individuals that had sequence data available and which were all genotyped only on the Axiom array (circled in green) had both of the alternate alleles at these probe-site SNPs. These individuals seem to have slightly diminished binding affinity in the Axiom array, as they are located at the cluster margin directed towards the center of the x-axis. The cluster position of one of these individuals was immediately next to that of an individual that had a discordant allele call (indicated by the black arrow), suggesting that one or both of these additional polymorphisms are causing the differential clustering observed. The individuals with differential clustering passed marker segregation checks in the Infinium cluster data whereas there were multiple Mendelian inconsistent and consistent errors across Axiom data, so the Illumina clustering was deemed accurate and the Axiom data was deemed inaccurate and thus the SNP was deemed discordant.

### Slide 8
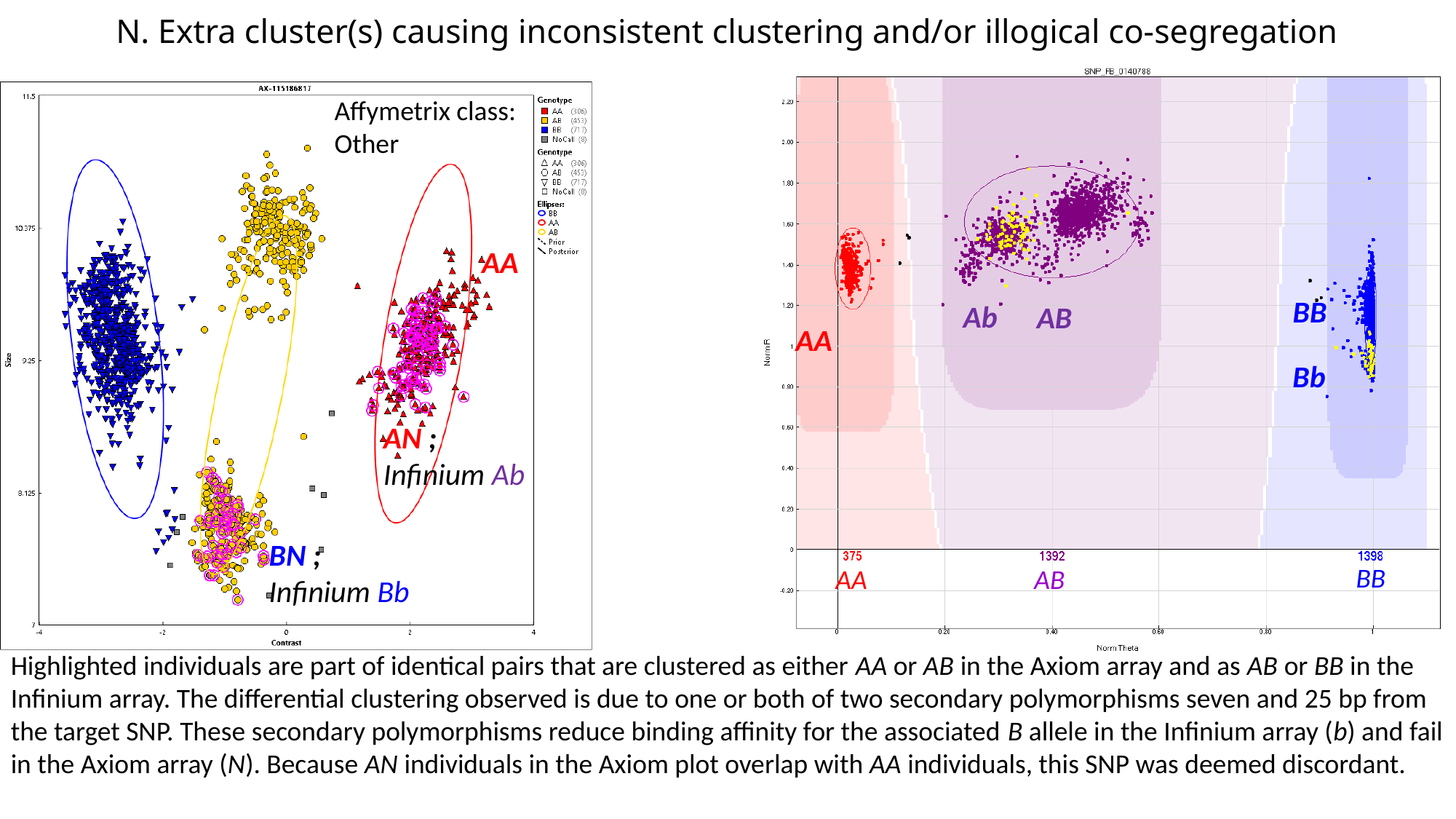

#### N. Extra cluster(s) causing inconsistent clustering and/or illogical co-segregation
Affymetrix class:
Other
AA
BB
Ab
AB
AA
Bb
AN ;
Infinium Ab
BN ;
Infinium Bb
BB
AA
AB
Highlighted individuals are part of identical pairs that are clustered as either AA or AB in the Axiom array and as AB or BB in the Infinium array. The differential clustering observed is due to one or both of two secondary polymorphisms seven and 25 bp from the target SNP. These secondary polymorphisms reduce binding affinity for the associated B allele in the Infinium array (b) and fail in the Axiom array (N). Because AN individuals in the Axiom plot overlap with AA individuals, this SNP was deemed discordant.

### Slide 9
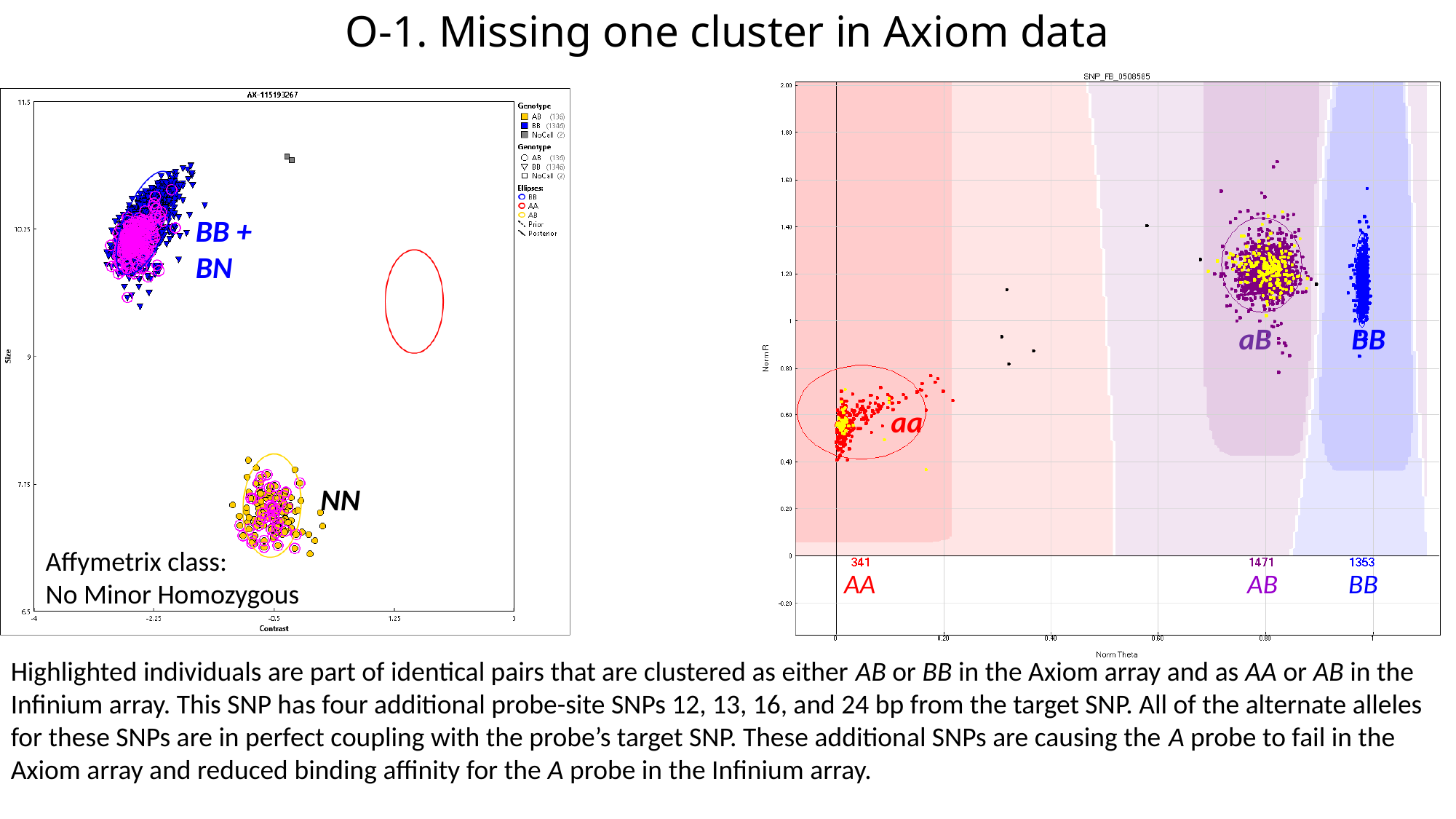

#### O-1. Missing one cluster in Axiom data
BB + BN
aB
BB
aa
NN
Affymetrix class:
No Minor Homozygous
AA
AB
BB
Highlighted individuals are part of identical pairs that are clustered as either AB or BB in the Axiom array and as AA or AB in the Infinium array. This SNP has four additional probe-site SNPs 12, 13, 16, and 24 bp from the target SNP. All of the alternate alleles for these SNPs are in perfect coupling with the probe’s target SNP. These additional SNPs are causing the A probe to fail in the Axiom array and reduced binding affinity for the A probe in the Infinium array.

### Slide 10
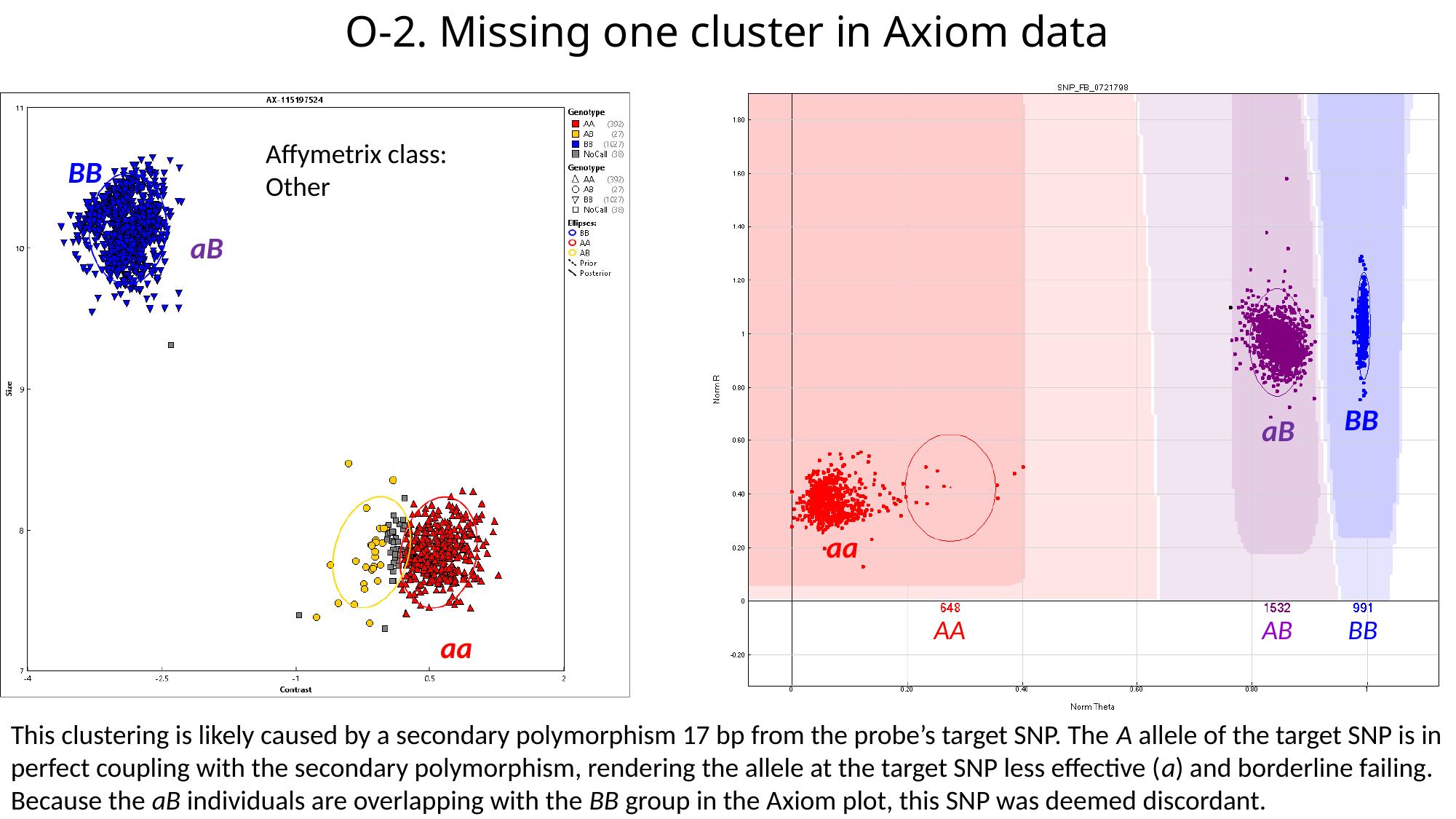

#### O-2. Missing one cluster in Axiom data
Affymetrix class:
Other
BB
aB
BB
aB
aa
AA
AB
BB
aa
This clustering is likely caused by a secondary polymorphism 17 bp from the probe’s target SNP. The A allele of the target SNP is in perfect coupling with the secondary polymorphism, rendering the allele at the target SNP less effective (a) and borderline failing. Because the aB individuals are overlapping with the BB group in the Axiom plot, this SNP was deemed discordant.

### Slide 11
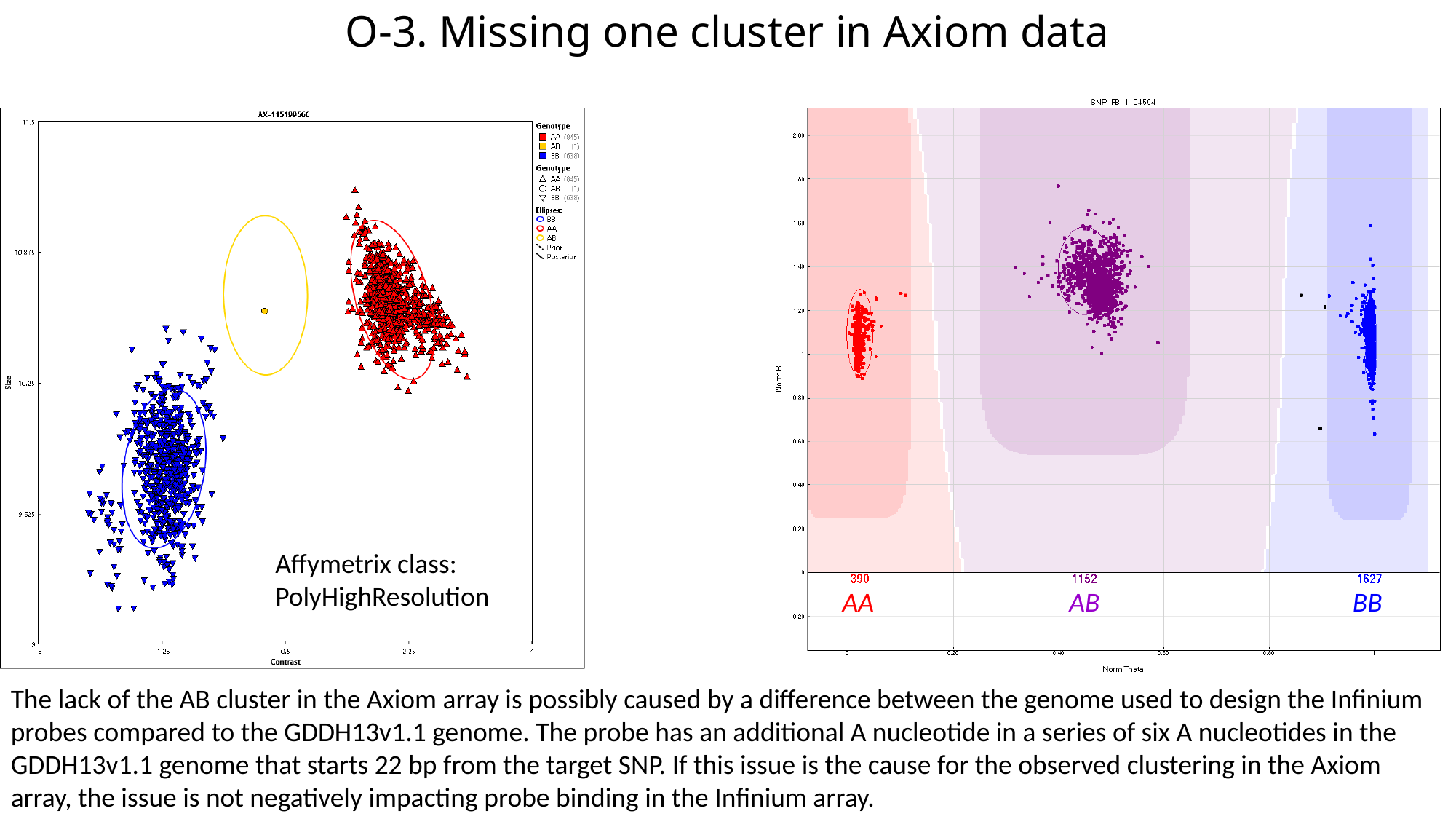

#### O-3. Missing one cluster in Axiom data
Affymetrix class:
PolyHighResolution
AA
AB
BB
The lack of the AB cluster in the Axiom array is possibly caused by a difference between the genome used to design the Infinium probes compared to the GDDH13v1.1 genome. The probe has an additional A nucleotide in a series of six A nucleotides in the GDDH13v1.1 genome that starts 22 bp from the target SNP. If this issue is the cause for the observed clustering in the Axiom array, the issue is not negatively impacting probe binding in the Infinium array.

### Slide 12
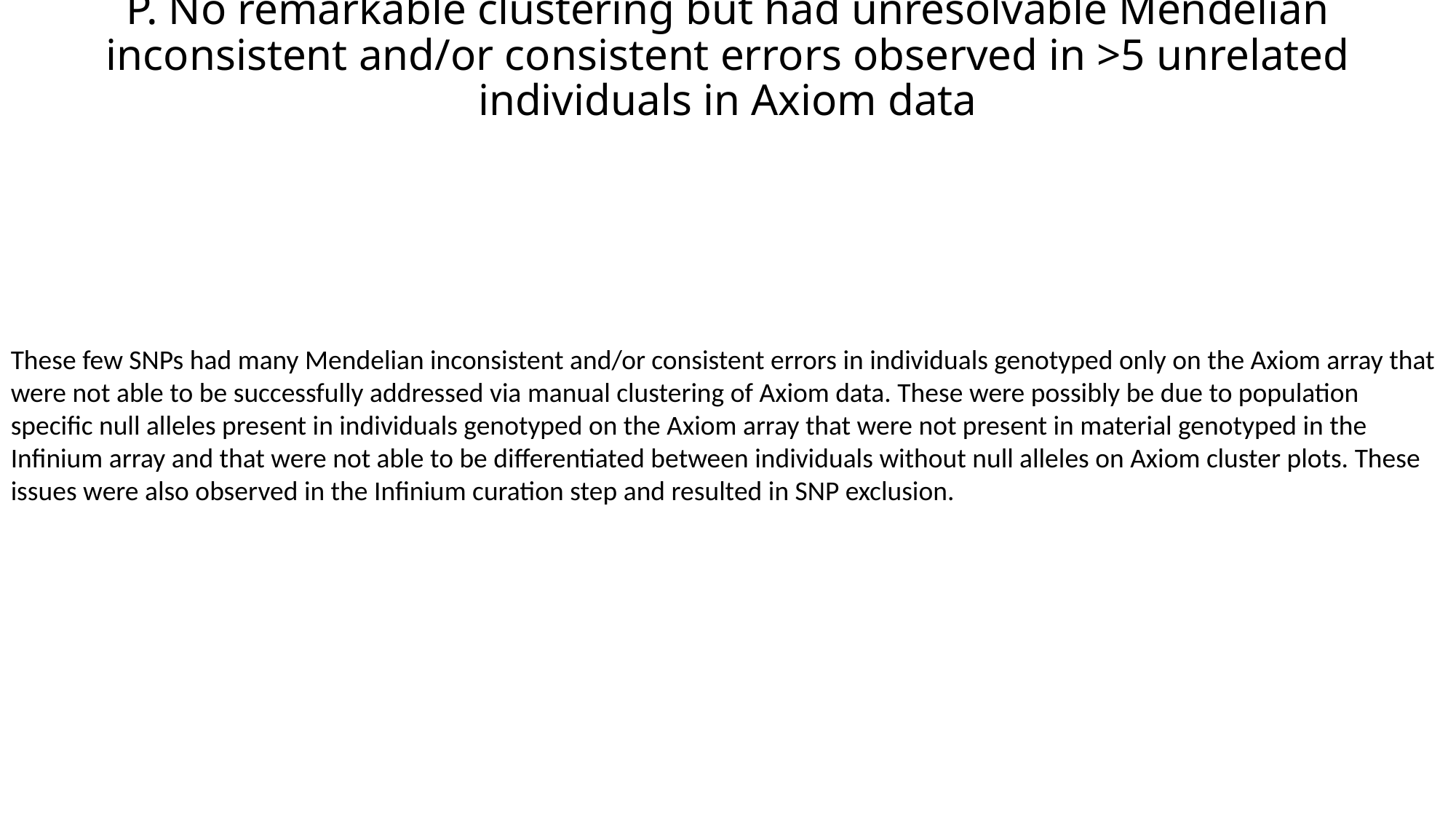

#### P. No remarkable clustering but had unresolvable Mendelian inconsistent and/or consistent errors observed in >5 unrelated individuals in Axiom data
These few SNPs had many Mendelian inconsistent and/or consistent errors in individuals genotyped only on the Axiom array that were not able to be successfully addressed via manual clustering of Axiom data. These were possibly be due to population specific null alleles present in individuals genotyped on the Axiom array that were not present in material genotyped in the Infinium array and that were not able to be differentiated between individuals without null alleles on Axiom cluster plots. These issues were also observed in the Infinium curation step and resulted in SNP exclusion.

### Slide 13
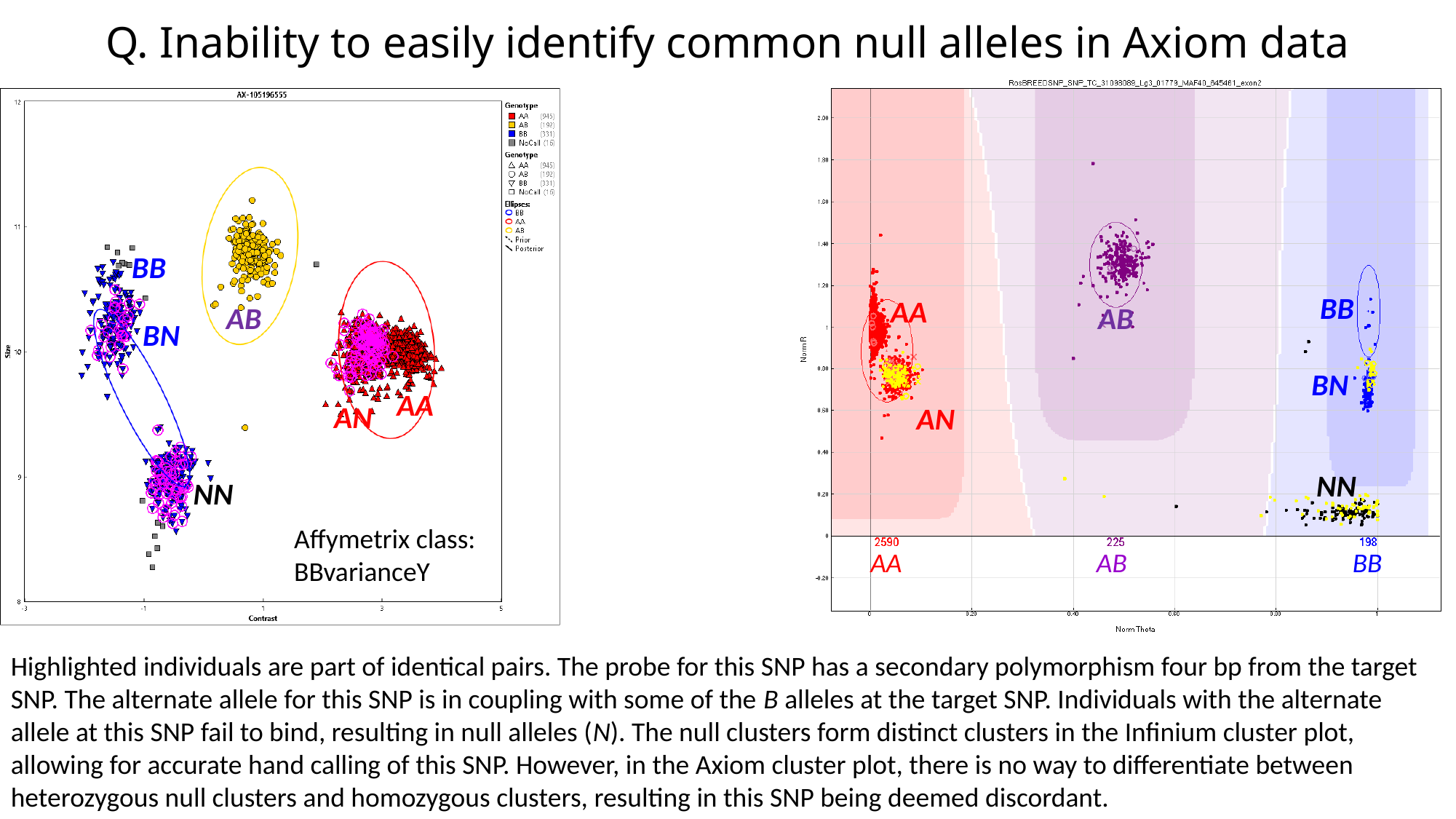

#### Q. Inability to easily identify common null alleles in Axiom data
BB
BB
AA
AB
AB
BN
BN
AA
AN
AN
NN
NN
Affymetrix class:
BBvarianceY
AA
AB
BB
Highlighted individuals are part of identical pairs. The probe for this SNP has a secondary polymorphism four bp from the target SNP. The alternate allele for this SNP is in coupling with some of the B alleles at the target SNP. Individuals with the alternate allele at this SNP fail to bind, resulting in null alleles (N). The null clusters form distinct clusters in the Infinium cluster plot, allowing for accurate hand calling of this SNP. However, in the Axiom cluster plot, there is no way to differentiate between heterozygous null clusters and homozygous clusters, resulting in this SNP being deemed discordant.
