## Additional file 7 for "Integration of Infinium and Axiom SNP array data in the outcrossing species *Malus* × *domestica* and causes for seemingly incompatible calls"

### Slide 1
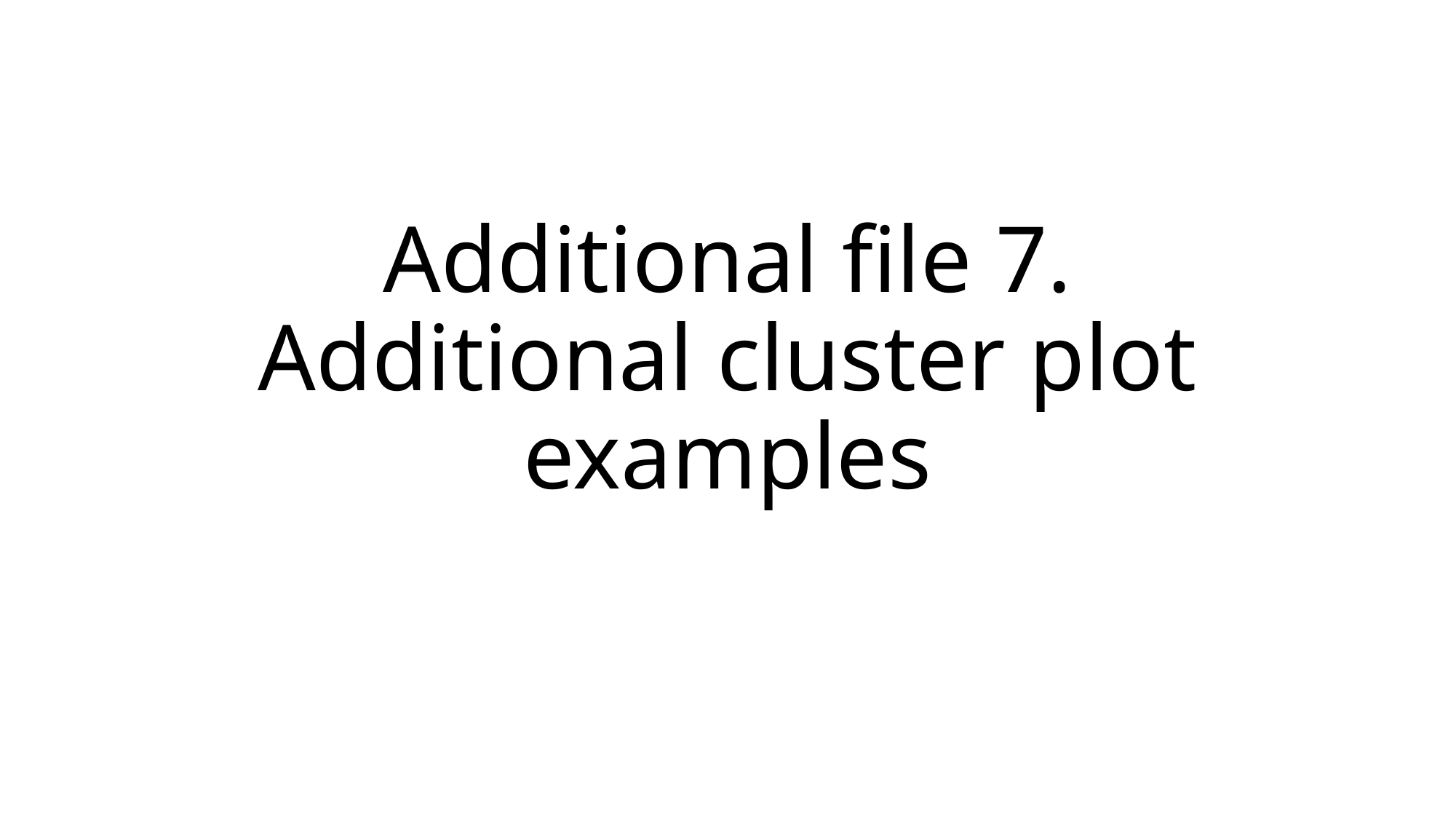

#### Additional file 7. Additional cluster plot examples

### Slide 2
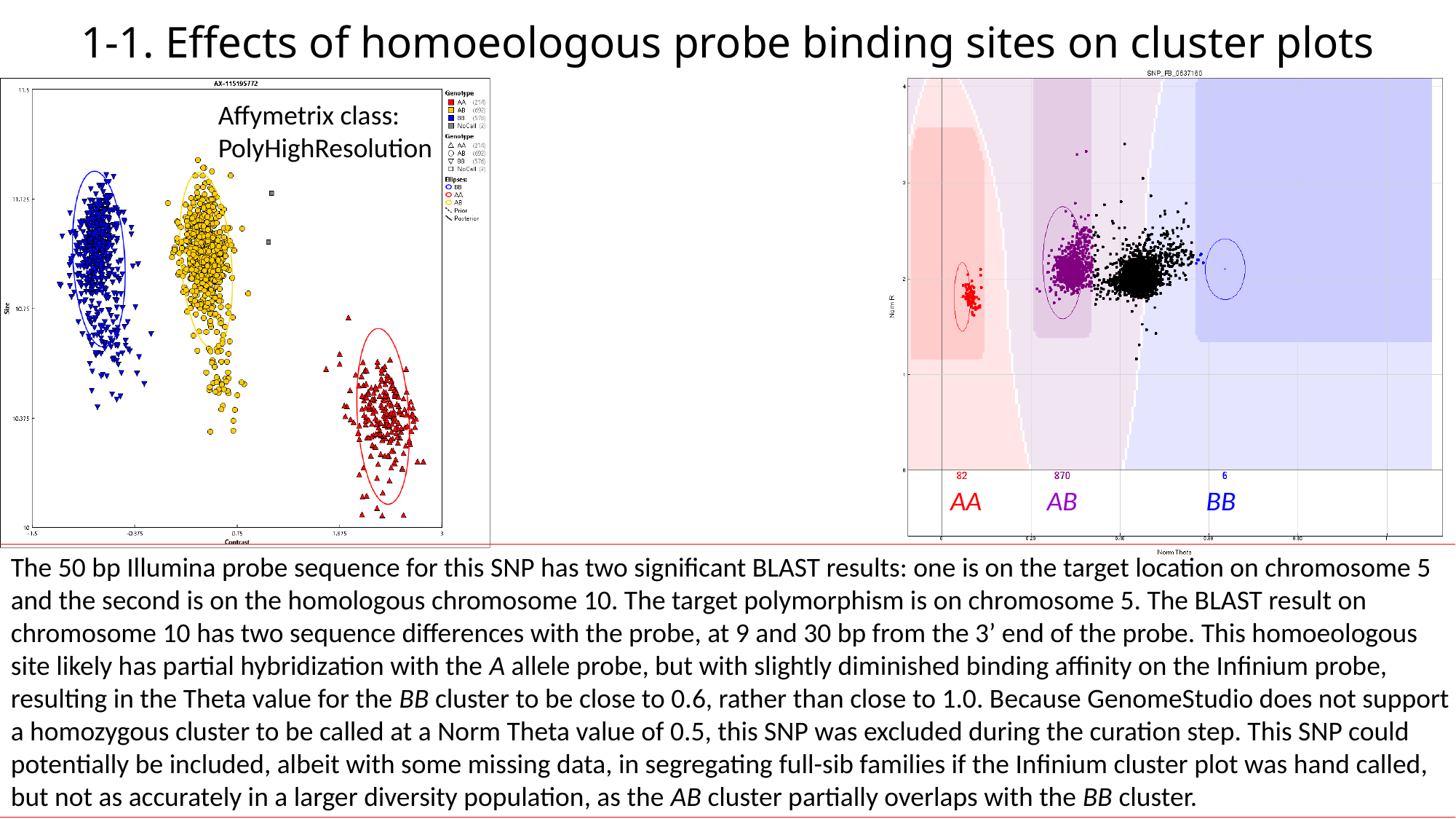

#### 1-1. Effects of homoeologous probe binding sites on cluster plots
Affymetrix class:
PolyHighResolution
AA
AB
BB
The 50 bp Illumina probe sequence for this SNP has two significant BLAST results: one is on the target location on chromosome 5 and the second is on the homologous chromosome 10. The target polymorphism is on chromosome 5. The BLAST result on chromosome 10 has two sequence differences with the probe, at 9 and 30 bp from the 3’ end of the probe. This homoeologous site likely has partial hybridization with the A allele probe, but with slightly diminished binding affinity on the Infinium probe, resulting in the Theta value for the BB cluster to be close to 0.6, rather than close to 1.0. Because GenomeStudio does not support a homozygous cluster to be called at a Norm Theta value of 0.5, this SNP was excluded during the curation step. This SNP could potentially be included, albeit with some missing data, in segregating full-sib families if the Infinium cluster plot was hand called, but not as accurately in a larger diversity population, as the AB cluster partially overlaps with the BB cluster.

### Slide 3
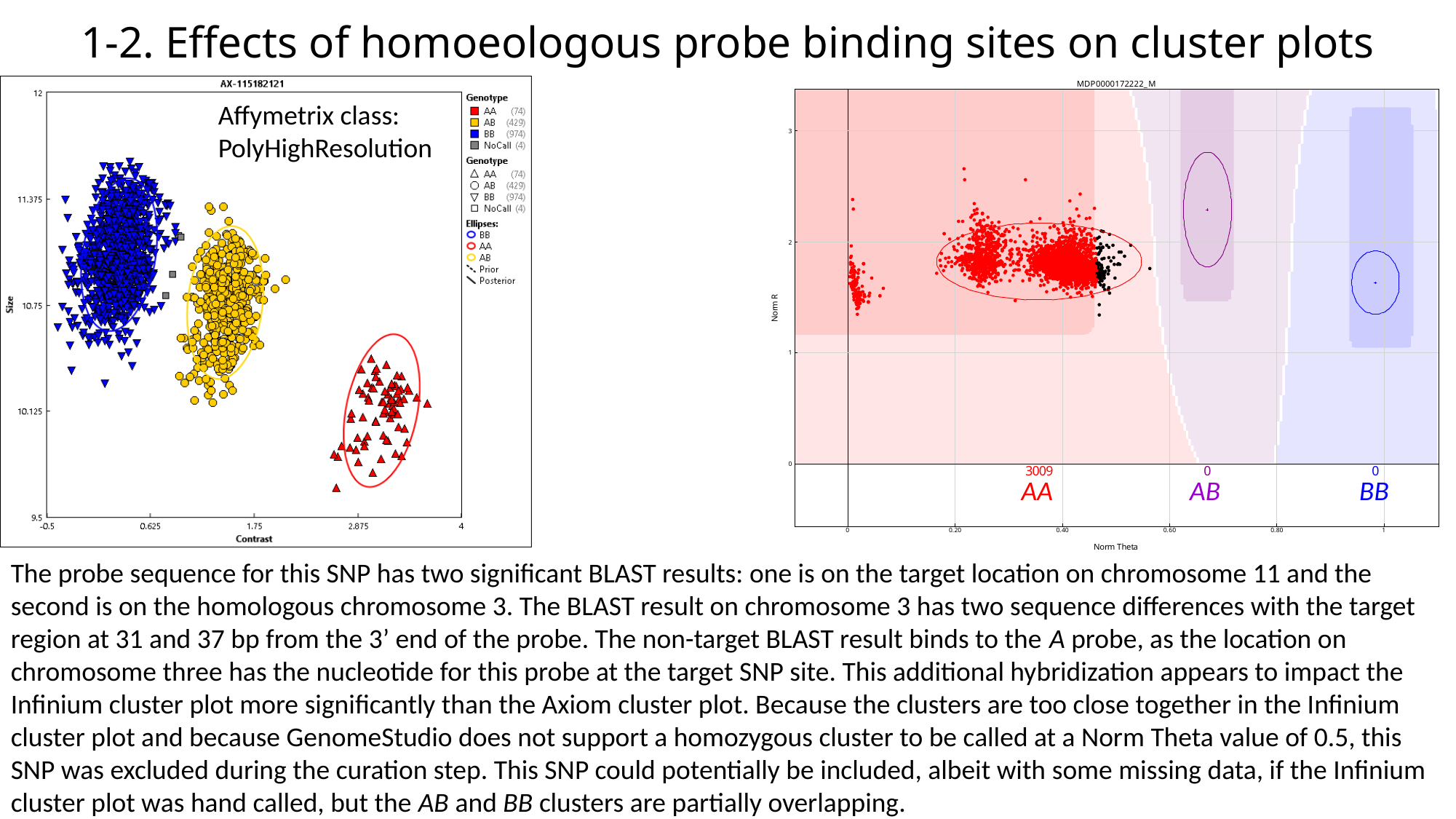

#### 1-2. Effects of homoeologous probe binding sites on cluster plots
Affymetrix class:
PolyHighResolution
AA
AB
BB
The probe sequence for this SNP has two significant BLAST results: one is on the target location on chromosome 11 and the second is on the homologous chromosome 3. The BLAST result on chromosome 3 has two sequence differences with the target region at 31 and 37 bp from the 3’ end of the probe. The non-target BLAST result binds to the A probe, as the location on chromosome three has the nucleotide for this probe at the target SNP site. This additional hybridization appears to impact the Infinium cluster plot more significantly than the Axiom cluster plot. Because the clusters are too close together in the Infinium cluster plot and because GenomeStudio does not support a homozygous cluster to be called at a Norm Theta value of 0.5, this SNP was excluded during the curation step. This SNP could potentially be included, albeit with some missing data, if the Infinium cluster plot was hand called, but the AB and BB clusters are partially overlapping.

### Slide 4
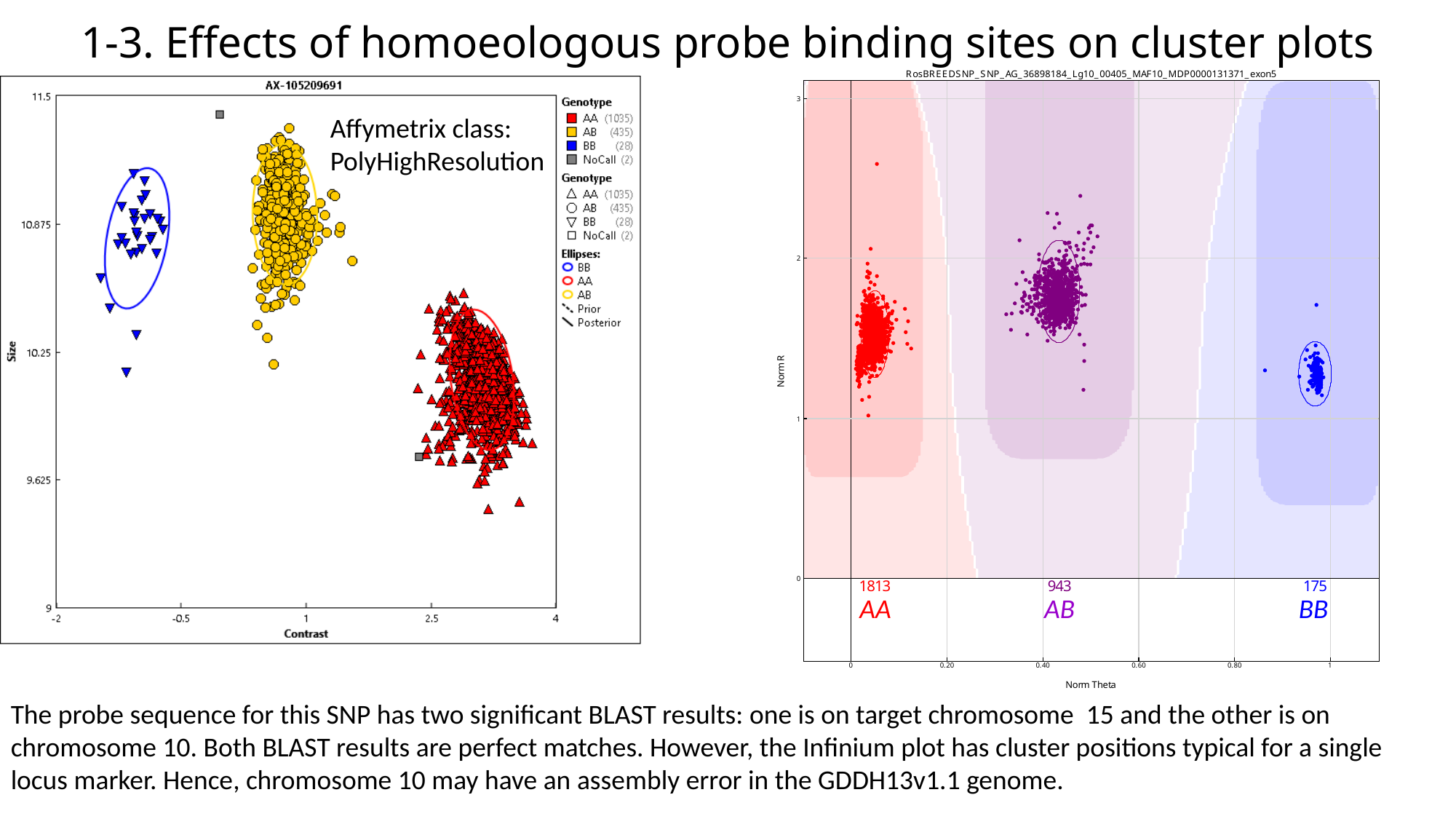

#### 1-3. Effects of homoeologous probe binding sites on cluster plots
Affymetrix class:
PolyHighResolution
AA
AB
BB
The probe sequence for this SNP has two significant BLAST results: one is on target chromosome 15 and the other is on chromosome 10. Both BLAST results are perfect matches. However, the Infinium plot has cluster positions typical for a single locus marker. Hence, chromosome 10 may have an assembly error in the GDDH13v1.1 genome.

### Slide 5
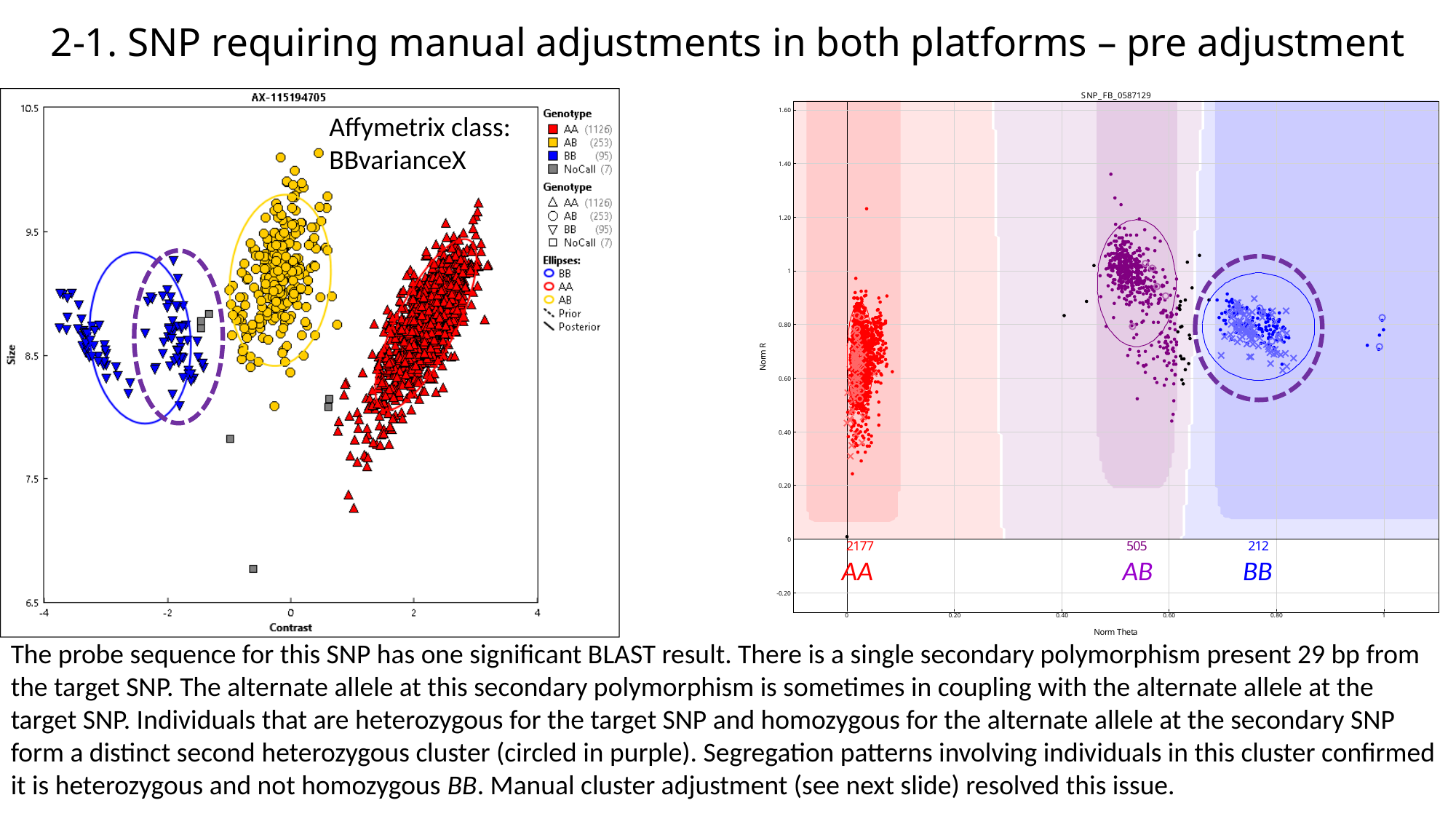

#### 2-1. SNP requiring manual adjustments in both platforms – pre adjustment
Affymetrix class:
BBvarianceX
AA
AB
BB
The probe sequence for this SNP has one significant BLAST result. There is a single secondary polymorphism present 29 bp from the target SNP. The alternate allele at this secondary polymorphism is sometimes in coupling with the alternate allele at the target SNP. Individuals that are heterozygous for the target SNP and homozygous for the alternate allele at the secondary SNP form a distinct second heterozygous cluster (circled in purple). Segregation patterns involving individuals in this cluster confirmed it is heterozygous and not homozygous BB. Manual cluster adjustment (see next slide) resolved this issue.

### Slide 6
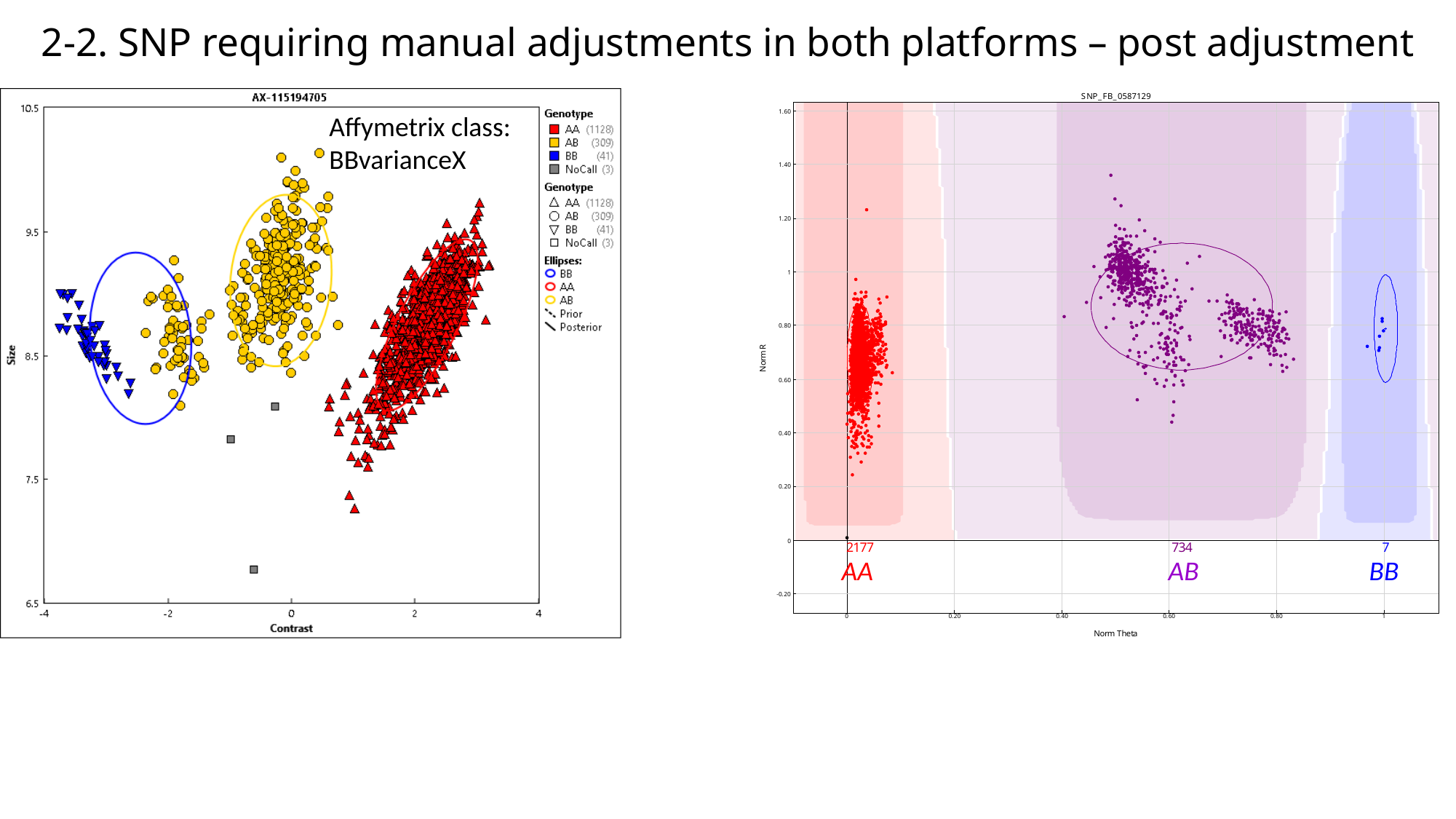

#### 2-2. SNP requiring manual adjustments in both platforms – post adjustment
Affymetrix class:
BBvarianceX
AA
AB
BB

### Slide 7
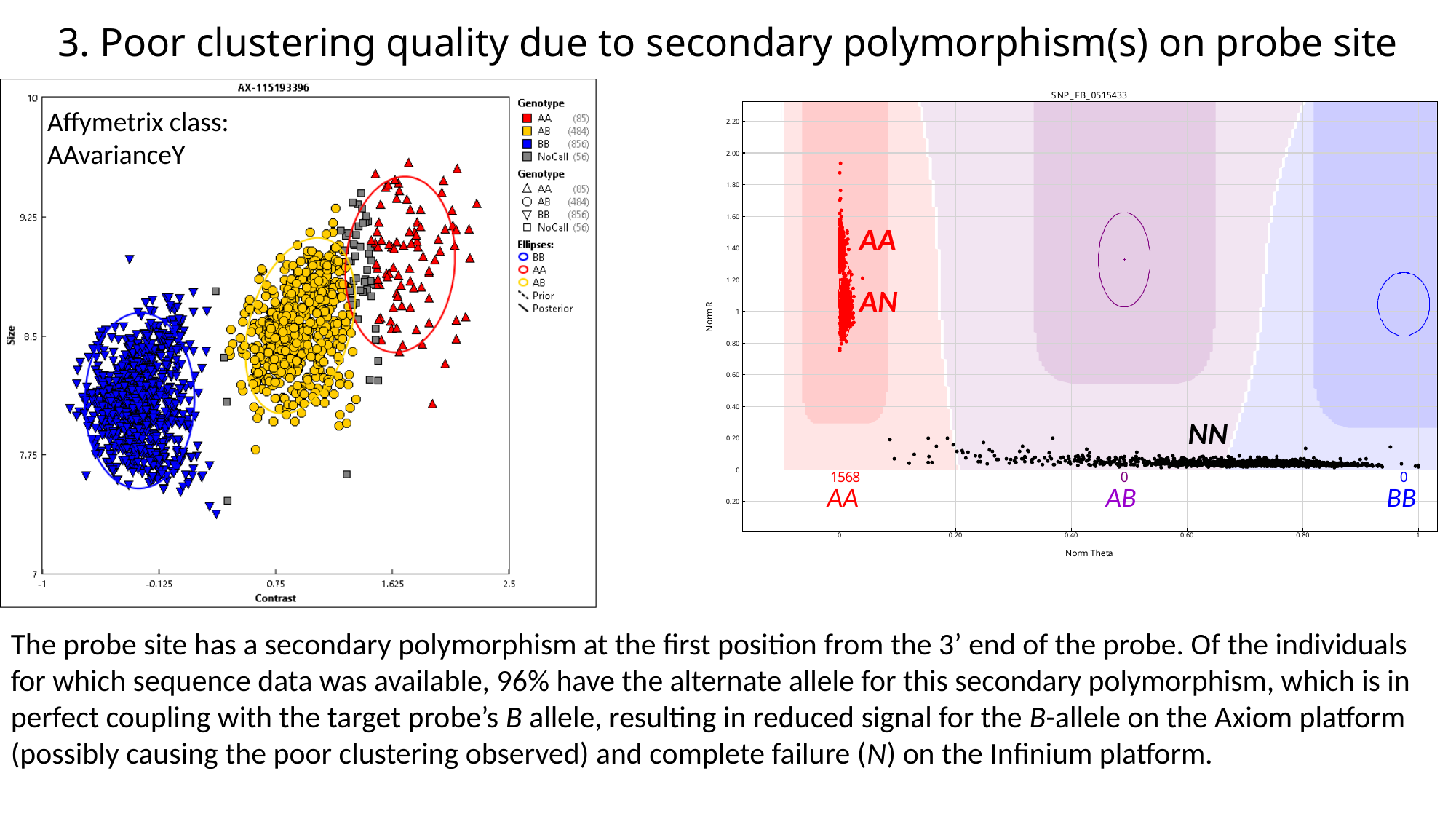

#### 3. Poor clustering quality due to secondary polymorphism(s) on probe site
Affymetrix class:
AAvarianceY
AA
AN
NN
AA
AB
BB
The probe site has a secondary polymorphism at the first position from the 3’ end of the probe. Of the individuals for which sequence data was available, 96% have the alternate allele for this secondary polymorphism, which is in perfect coupling with the target probe’s B allele, resulting in reduced signal for the B-allele on the Axiom platform (possibly causing the poor clustering observed) and complete failure (N) on the Infinium platform.

### Slide 8
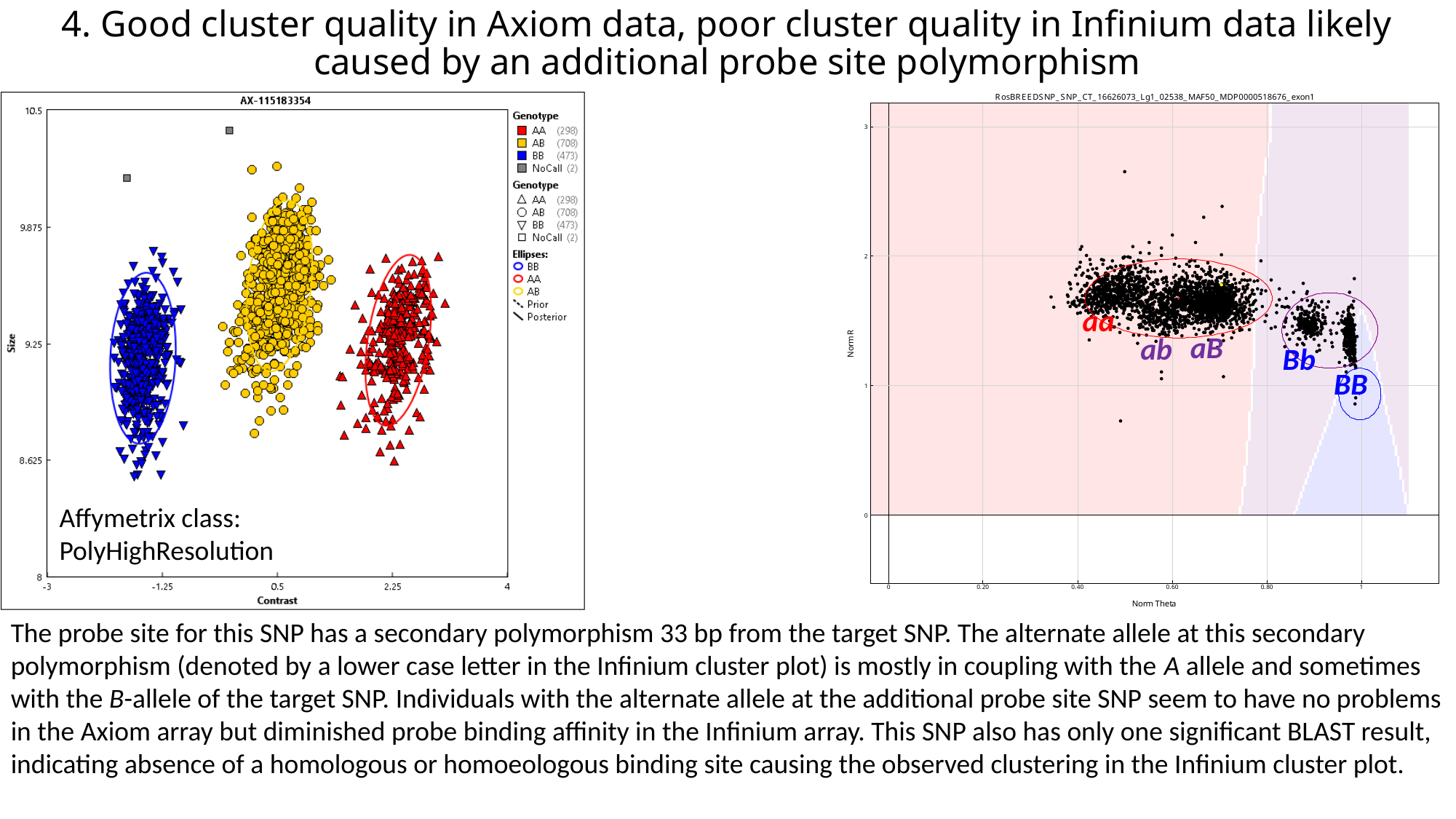

#### 4. Good cluster quality in Axiom data, poor cluster quality in Infinium data likely caused by an additional probe site polymorphism
aa
aB
ab
Bb
BB
Affymetrix class:
PolyHighResolution
The probe site for this SNP has a secondary polymorphism 33 bp from the target SNP. The alternate allele at this secondary polymorphism (denoted by a lower case letter in the Infinium cluster plot) is mostly in coupling with the A allele and sometimes with the B-allele of the target SNP. Individuals with the alternate allele at the additional probe site SNP seem to have no problems in the Axiom array but diminished probe binding affinity in the Infinium array. This SNP also has only one significant BLAST result, indicating absence of a homologous or homoeologous binding site causing the observed clustering in the Infinium cluster plot.

### Slide 9
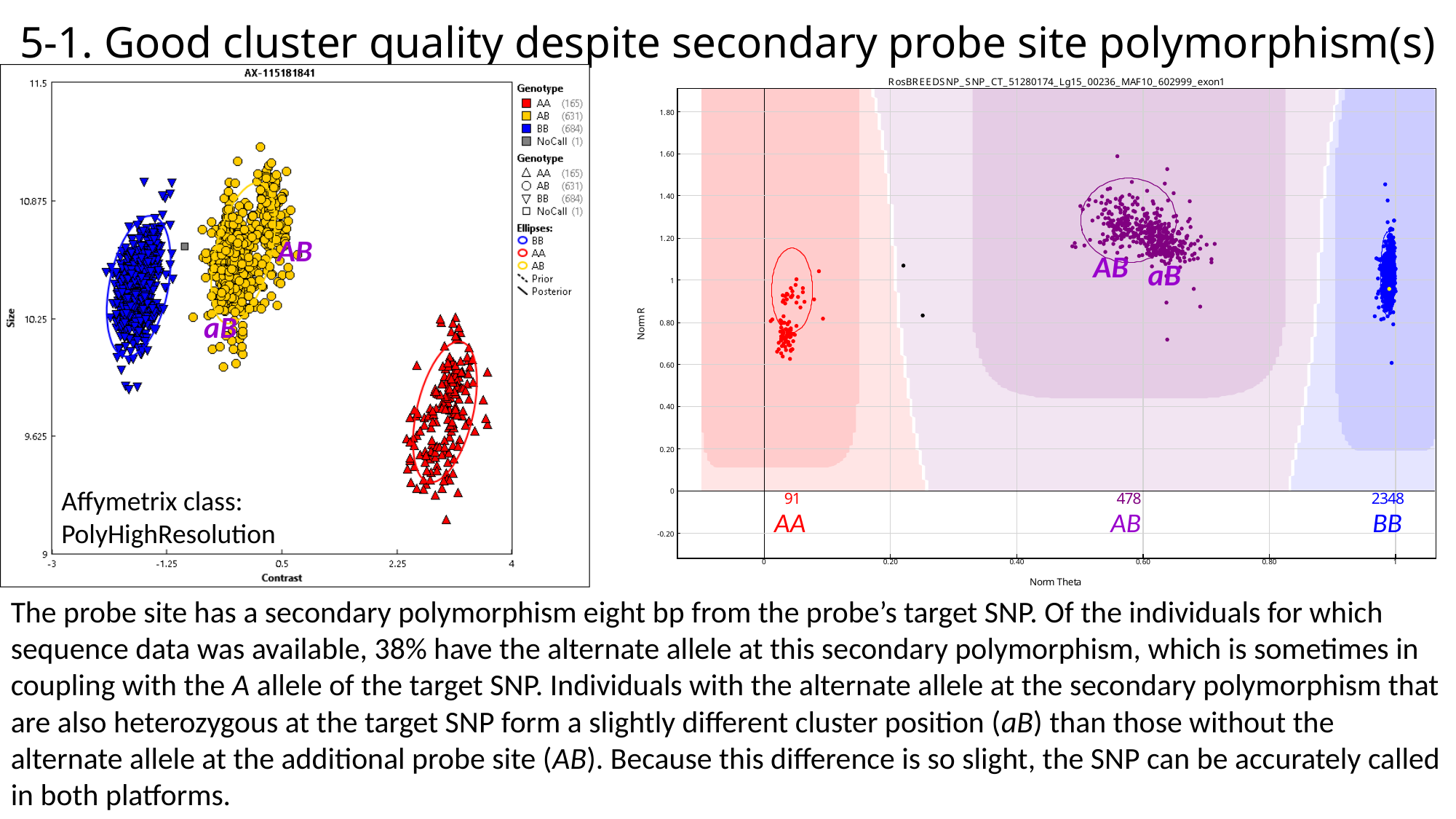

#### 5-1. Good cluster quality despite secondary probe site polymorphism(s)
AB
AB
aB
aB
Affymetrix class:
PolyHighResolution
AA
AB
BB
The probe site has a secondary polymorphism eight bp from the probe’s target SNP. Of the individuals for which sequence data was available, 38% have the alternate allele at this secondary polymorphism, which is sometimes in coupling with the A allele of the target SNP. Individuals with the alternate allele at the secondary polymorphism that are also heterozygous at the target SNP form a slightly different cluster position (aB) than those without the alternate allele at the additional probe site (AB). Because this difference is so slight, the SNP can be accurately called in both platforms.

### Slide 10
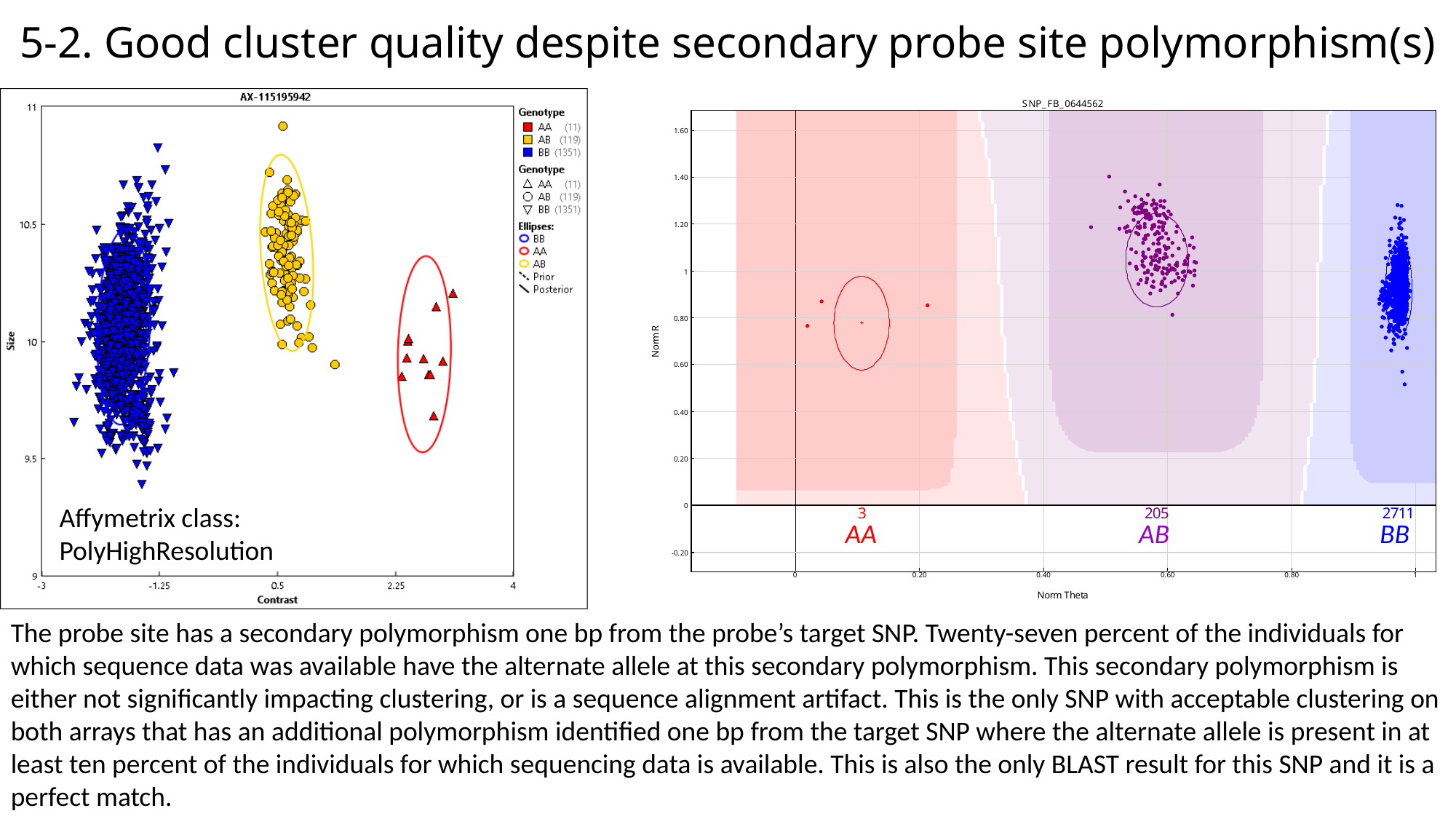

#### 5-2. Good cluster quality despite secondary probe site polymorphism(s)
Affymetrix class:
PolyHighResolution
AA
AB
BB
The probe site has a secondary polymorphism one bp from the probe’s target SNP. Twenty-seven percent of the individuals for which sequence data was available have the alternate allele at this secondary polymorphism. This secondary polymorphism is either not significantly impacting clustering, or is a sequence alignment artifact. This is the only SNP with acceptable clustering on both arrays that has an additional polymorphism identified one bp from the target SNP where the alternate allele is present in at least ten percent of the individuals for which sequencing data is available. This is also the only BLAST result for this SNP and it is a perfect match.
